# Mechanistic basis for GPCR phosphorylation-dependent allosteric signaling specificity of *β*-arrestin 1 and 2

**DOI:** 10.1101/2025.04.29.649777

**Authors:** Midhun K Madhu, Rajesh K Murarka

## Abstract

*β*-arrestins (*β*arr1 and *β*arr2) are key transducers of G protein-coupled receptor (GPCR) signaling, orchestrating both shared and isoform-specific intracellular pathways. Phosphorylation of the receptor C-terminal tail by GPCR kinases encodes regulatory “barcodes” that modulate *β*-arrestin conformations and interactions with downstream effectors. However, how distinct phosphorylation patterns shape *β*-arrestin structure and function remains poorly understood. Here, we combine all-atom molecular dynamics simulations with machine learning and graph neural networks to systematically map the barcode-dependent conformational landscapes of *β*-arrestins bound to the phosphorylated vasopressin receptor 2 tail (V2Rpp). We find that V2Rpp engages *β*arr1 more stably than *β*arr2, mediated by isoform-specific residue contacts that trigger distinct allosteric responses. These include differential interdomain rotations and rearrangements in key structural motifs, potentially facilitating selective effector protein engagement. Furthermore, we identify critical residue networks that transmit phosphorylation signals to effector-binding interfaces in a barcode- and isoform-specific manner. Notably, *β*arr1 exhibits stronger allosteric coupling between V2Rpp and c-edge loop 2 compared to *β*arr2, which is consistent with its enhanced membrane association. Together, these findings advance our understanding of the molecular mechanisms by which *β*-arrestins interpret GPCR phosphorylation signatures, offering a framework that could aid in designing pathway-selective therapeutics.

## Introduction

*β*-arrestins (*β*arrs) play a pivotal role as signal transducers for G protein-coupled receptors (GPCRs), the largest family of cell surface receptors, which are the targets of over a third of currently marketed drugs. ^1–5^ Phosphorylation by G protein receptor kinases (GRKs) in the C-terminal tail (C-tail) of GPCRs enables the binding of *β*arrs, causing desensitization of G protein-mediated signaling and receptor internalization. ^6^ *β*arrs further act as scaffolds for a diverse range of effector proteins, thereby initiating G protein-independent downstream signaling cascades involved in cell growth, proliferation, and differentiation. ^2,7^

Unlike visual arrestins (arrestin-1 and arrestin-4), the *β*arr isoforms, *β*arr1 and *β*arr2 (arrestin-2 and arrestin-3, respectively; Figure 1A, B) exhibit remarkable promiscuity, engaging with hundreds of non-visual GPCRs to elicit both overlapping and distinct cellular responses. ^2,8^ The functional diversity of *β*arrs is driven by GRK-mediated receptor C-tail phosphorylation, where unique phospho-patterns stabilize distinct active *β*arr conformations, leading to selective effector interactions and enabling precise modulation of downstream signaling pathways. ^9–11^ GPCRs and *β*arrs engage in a bimodal interaction mechanism, where the phosphorylated C-tail of the receptor interacts with a crevice in the N-domain of *β*arrs containing several positively charged residues, forming a ‘hanging complex’ (partially engaged) ^12^ and further the finger loop of *β*arrs inserts into the receptor core, while the c-edge loops anchor to the membrane, forming a ‘core complex’ (fully engaged), ^13,14^ as shown in Figure S1A. Both of these modes can activate *β*arrs to varying degrees, thereby regulating downstream functional responses, as demonstrated by previous experimental studies. ^15,16^ The core complex has been shown to be essential for desensitizing G protein signaling by sterically occluding the receptor’s intracellular binding cavity, whereas the hanging complex is sufficient for receptor internalization via clathrin-mediated endocytosis. Additionally, the hanging complex exposes the central crest region of *β*arrs involved in receptor binding, making it accessible to other effectors. ^17^

**Figure 1.**
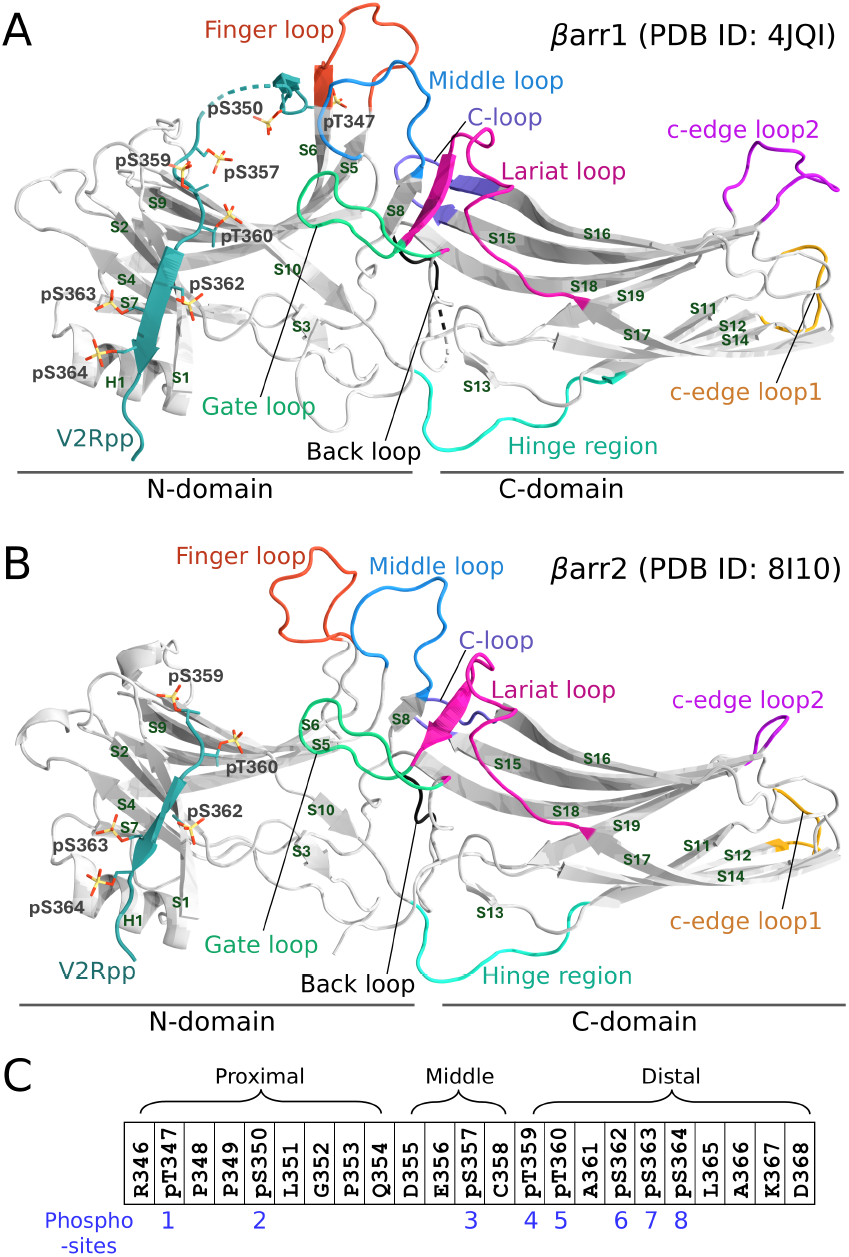
Cartoon representations of the active states of **(A)** *β*arr1 and **(B)** *β*arr2 bound to the phosphorylated V2R C-tail (full phospho-pattern, with all serine and threonine residues phosphorylated). Key structural elements in *β*arr1 (and in *β*arr2) are highlighted in distinct colors and are described as follows: the lariat loop (residues L274-L300 in *β*arr1; L275-L301 in *β*arr2), which contains the gate loop (D290-N299 in *β*arr1; D291-N300 in *β*arr2); the finger loop (G64-T74; G65-T75); the middle loop (Q130-A139; Q131-A140); the C-loop (I241-Q248; I242-Q249); the back loop (R312-L315; R313-L316); the c-edge loop1 (L191-P196; L192-S197); the c-edge loop2 (S330-S340; S331-G334); and the hinge region (A174-T183; A175-S184). **(C)** V2R C-tail phosphopeptide, with phosphosites 1 to 8 highlighted.

Numerous structural and spectroscopic investigations have explored the atomistic mechanisms underlying phospho-pattern-dependent signaling by *β*arrs. ^17–19^ Recent molecular dynamics (MD) simulations, complemented by experimental validation, have provided compelling evidence that the spatial arrangement of phosphates within the C-tail of the vasopressin receptor 2 (V2Rpp), rather than the mere number of phosphorylated residues, plays a crucial role in dictating *β*arr1 conformations and subsequent signaling outcomes. ^20^ Furthermore, this study reveals that distinct phospho-patterns induce unique conformational changes in *β*arr1, including differential rotation of its C-domain relative to the N-domain and altered positioning of key loop regions connecting *β*-sheets and spanning the interdomain crevice. Importantly, these conformational changes occur independently of one another, highlighting the intricate nature of phospho-pattern-dependent *β*arr1 activation. ^20^ X-ray crystallography has been instrumental in elucidating critical phospho-patterns within GPCRs, particularly the PXPXXP and PXXPXXP motifs (where P denotes phosphoserine or -threonine, and X represents any other residue), which are essential for *β*arr recruitment. ^21^ Recently, the structure determination of *β*arr1 and *β*arr2 in complex with the C-terminus of various GPCRs identified a specific PXPP motif as crucial for *β*arr binding and activation when present in the interacting receptor. It interacts with a spatially organized lysine-arginine motif (KKRRKK sequence) in the N-domain of *β*arrs. ^22,23^ Sequence analysis across the human GPCRome indicated that the PXPP motif is conserved among many receptors, confirming its role in *β*arr activation and functional engagement. Additionally, a series of X-ray structures of *β*arr1 were determined to investigate how arrestins recognize distinct phospho-patterns on GPCRs and translate them into selective functions. ^24^ This study also employed FRET and NMR to reveal how small changes in phospho-interactions can cause significant alterations in *β*arr1 conformations. Furthermore, recent studies utilizing a range of biophysical approaches have revealed that the phospho-patterns on different regions of the GPCR C-tail can elicit distinct functions of *β*arrs. ^25,26^

Despite significant progress in elucidating the activation and signaling mechanisms of *β*arrs, critical knowledge gaps remain in our understanding of how diverse phospho-patterns on GPCRs govern the complex conformational landscapes of *β*arr isoforms and ultimately determine their functional outcomes. While structural insights from X-ray crystallography and cryo-EM have been invaluable, these techniques provide limited information about the dynamics, particularly the interactions among GPCRs, *β*arrs, and downstream effector proteins. Moreover, the reliance on stabilizing antibodies in many structural studies raises questions about the physiological relevance of the observed conformations. While prior research on *β*arr1 has laid the foundation for understanding *β*arr activation, ^19,20^ the mechanistic details of phosphorylation-dependent signaling in both isoforms remain incompletely characterized. A comprehensive understanding of the allosteric mechanisms governing *β*arr1 and *β*arr2 signaling requires a detailed investigation of their dynamic behavior in response to interactions with diverse phosphorylation barcodes on GPCRs. However, achieving this through experimental approaches, such as spectroscopic methods, remains challenging, as these techniques typically provide localized information and often fail to accurately quantify the relationship between probe responses and precise conformational changes. To address these challenges, we employed an integrative computational framework that combines molecular dynamics (MD) simulations and machine learning (ML) to investigate the functional dynamics of *β*arrs induced by distinct phospho-patterns on the C-terminal tail of V2R. Our findings reveal phosphorylation-dependent conformational changes in both *β*arr1 and *β*arr2, underscoring their residue-specific differences. Notably, distinct phospho-patterns selectively modulate key structural motifs and effector-interacting regions in *β*arrs through unique allosteric communication pathways originating from the V2R C-tail.

## Results

To elucidate the molecular basis of phospho-pattern-dependent regulation of *β*arr isoforms, we conducted extensive MD simulations of *β*arr1 and *β*arr2 bound to five functionally distinct, differentially phosphorylated C-terminal peptides of the vasopressin receptor 2 (V2Rpp; residues 346-368). ^22,24,27^ The fully phosphorylated (FP) system for *β*arr1 was modeled based on an active X-ray crystal structure complexed with V2Rpp, ^27^ in which all serine and threonine residues on V2R C-tail are phosphorylated: pS347, pS350, pS357, pT359, pY360, pS362, pS363, and pS364, labeled P1-P8 (Figure 1C). The other systems were modeled using X-ray structures with distinct phospho-patterns, each lacking phosphorylation at one (P1, P3, or P5) or two (P7/8) specific sites relative to the FP state ^24^ (Figure S1B). These *β*arr1 systems are designated as *β*arr1_V2R(FP), *β*arr1_V2R(–P1), *β*arr1_V2R(–P3), *β*arr1_V2R(–P5), and *β*arr1_V2R(–P7/8). For the *β*arr2_V2R(FP) system, initial coordinates were adapted from an active cryo-EM structure. ^22^ Phospho-patterns similar to those in the *β*arr1 systems were incorporated in silico to generate *β*arr2_V2R(–P1), *β*arr2_V2R(–P3), *β*arr2_V2R(–P5), and *β*arr2_V2R(–P7/8). Seven independent MD trajectories, each 1 *µ*s in duration, were generated per system, totaling 70 *µ*s of simulation data (see SI Methods for details). Unless otherwise specified, results are based on average properties computed from the final 600 ns of each trajectory. The names of the *β*arr structural elements follow the ArrestinDb nomenclature: ^28^ *β*-sheets (S) and helices (H) are identified by their positional numbers, while loops connecting them are named according to their adjacent elements. For example, H1 represents helix 1, S4 denotes *β*-sheet 4, and s5s6 refers to the loop between S5 and S6.

### V2Rpp phosphorylation pattern dictates isoform-specific *β*arr binding characteristics

The binding stability and contact formation of phosphopeptide with *β*arr1 or *β*arr2 vary depending on the phospho-pattern, significantly influencing the conformational landscape of *β*arrs, as shown in previous studies. ^20,22,24,29^ To quantify the binding stability of V2Rpp, we calculated a ‘stability score’ as previously defined ^20^ and detailed in the SI Methods. For *β*arr1, the absence of specific phosphorylation sites on V2Rpp did not markedly affect overall binding stability, as indicated by the presence of prominent peaks at high stability values across all variants (Figure 2). This observation aligns with prior experimental findings demonstrating stable complex formation between *β*arr1 and all V2Rpp variants. ^24^ Notably, the –P3 and –P5 variants exhibited broader distributions and secondary peaks at moderately lower values, suggesting increased fluctuations in V2Rpp binding stability. These findings suggest that the phosphate groups at sites P3 and P5 (pS357 and pT360) play pivotal roles in stabilizing V2Rpp, particularly when supported by additional phosphorylations on the V2Rpp. The reduced stability in the –P3 and –P5 systems likely stems from the loss of key intermolecular contacts in the middle region of the complex due to missing phosphate groups (Figure S3; Tables S2, S3).

**Figure 2.**
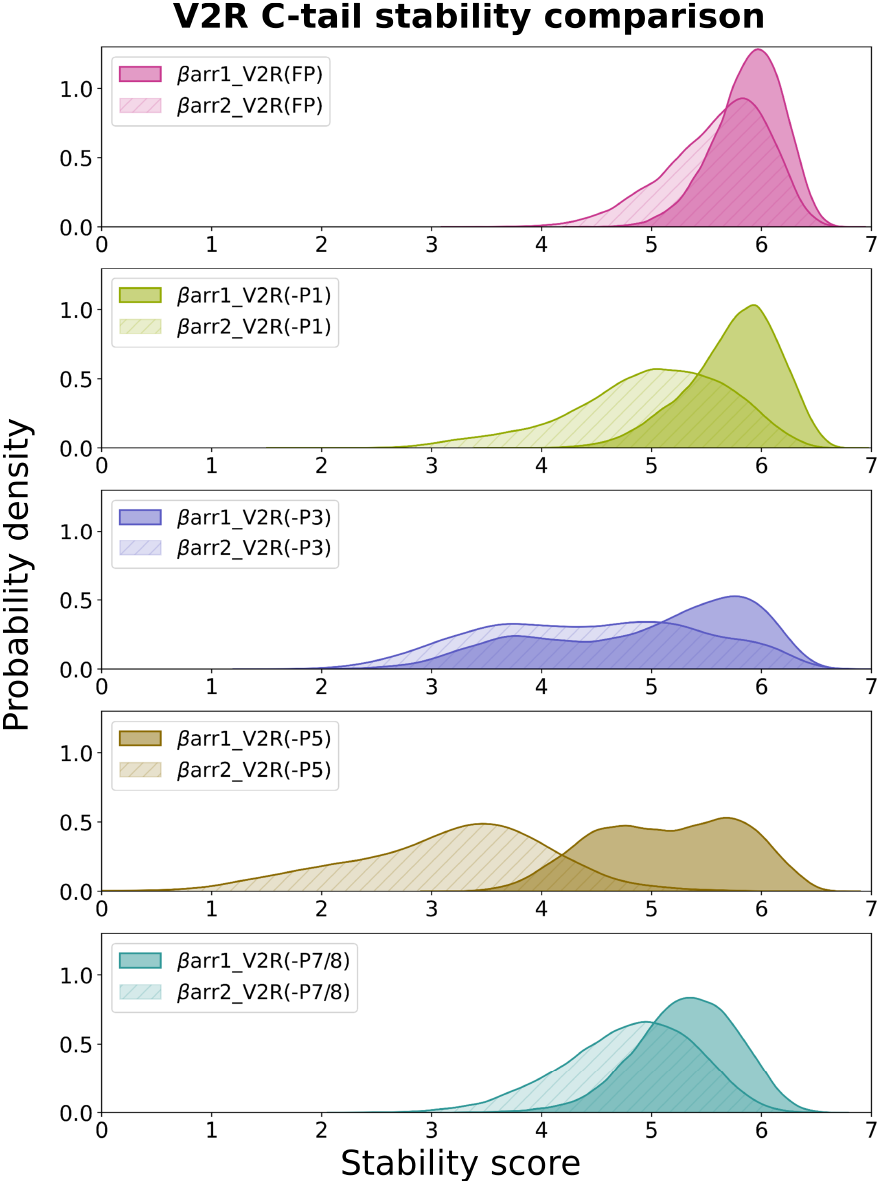
Variation in binding stability of V2Rpp with distinct phospho-patterns in complex with *β*arr1 and *β*arr2. The stability score is calculated as the 7 Å – r.m.s.d. of V2Rpp relative to its initial conformation. ^20^

In all *β*arr2 systems except FP, V2Rpp stability is markedly reduced compared to *β*arr1 (Figure 2), due to fewer residue-level contacts, particularly within the proximal and middle regions of the phosphopeptide (Figures S2A–S2D). The proximal and middle regions of V2Rpp (residues 346–357) are inherently flexible in the case of *β*arr2, which remained unresolved in the fully phosphorylated *β*arr2-V2Rpp structure. ^22^ Consequently, the modeled proximal and middle segments of V2Rpp failed to establish critical contacts in our simulations. Notably, the absence of T360 phosphorylation in the –P5 variant results in the lowest stability score, underscoring the critical role of pT360 in stabilizing *β*arr2-V2Rpp interactions. pT360, the first phosphorylated residue within the PXPP motif–previously identified as essential for *β*arr binding and activation ^22,23^–shows reduced interaction with K295 in *β*arr2 (Figure S2E), whereas it engages more strongly with the corresponding lysine residue (K294) in *β*arr1. In the –P5 system, the absence of pT360 causes a shift in interaction sites from K294 to K11, K25, and K10 in *β*arr1, and from K295 to K12, K26, and K11 in *β*arr2. These alternative interactions are notably weaker in *β*arr2 than in *β*arr1 (Figure S2E), consistent with the reduced V2Rpp stability observed in *β*arr2.

Further analysis revealed that phosphorylated residues in V2Rpp primarily engage in polar, charge-charge, and hydrogen bonding interactions with *β*arr1, consistent with prior structural studies ^24,27^ (Figure S3; Tables S1-S4). In contrast, *β*arr2 exhibits significant differences, including altered interaction partners for the phosphorylated residues and the loss of several key contacts (Figures S4; Tables S5-S8). These variations underlie isoform-specific V2Rpp binding and correlate with the observed differences in V2Rpp peptide stability. The phospho-pattern-dependent differential interactions in *β*arr isoforms suggest unique allosteric triggers from V2Rpp that influence distal functional regions and regulate the binding of effector proteins.

### Key structural motifs in *β*arrs adopt phosphorylation-dependent conformations

To investigate the conformational changes induced by differential phospho-patterns on V2Rpp in *β*arrs, we examined both global and local alterations, including interdomain rotation and the dynamics of key structural motifs. Interdomain rotation is a hallmark of *β*arr activation, typically reaching *∼* 20^*°*^ in fully active structures. ^2^ Previous experiments and simulations have shown that *β*arrs adopt a spectrum of activation states, with the degree of rotation varying based on the phosphorylation pattern of the bound phosphopeptide. ^20,22–24,27,29,30^ To evaluate the impact of phospho-patterns on *β*arr activation, we quantified the rotation of the C-domain relative to the N-domain in both *β*arr1 and *β*arr2 systems. In line with previous reports, ^20^ the removal of the active-state-stabilizing antibody (Fab30) in our simulations led to a substantial reduction in domain rotation in *β*arr1 (Figure 3A), shifting the distributions toward lower angles, with averages ranging from 8.1^*°*^ in –P1 to 14.5^*°*^ in –P3. Phosphorylation at P3 (S357) has been shown to suppress activation by displacing K294 in the gate loop—a segment of the lariat loop connecting *β*-sheets S17 and S18—thus disrupting its interactions with pT359 and pT360. ^20^ Accordingly, the enhanced interdomain rotation observed in the –P3 variant, which lacks phosphorylation at S357, is consistent with previous findings (Figure S3). In contrast, the reduced rotation observed in the –P1 variant may stem from the absence of the pT347 phosphate group, which normally interacts with residues in the finger loop that connects *β*-sheets S5 and S6 (Figure S3). The finger loop is a critical receptor-interacting element and serves as an activation sensor for *β*arrs in receptor core-bound complexes. ^31^ Local perturbations in this region may, therefore, influence the extent of global domain rotation.

**Figure 3.**
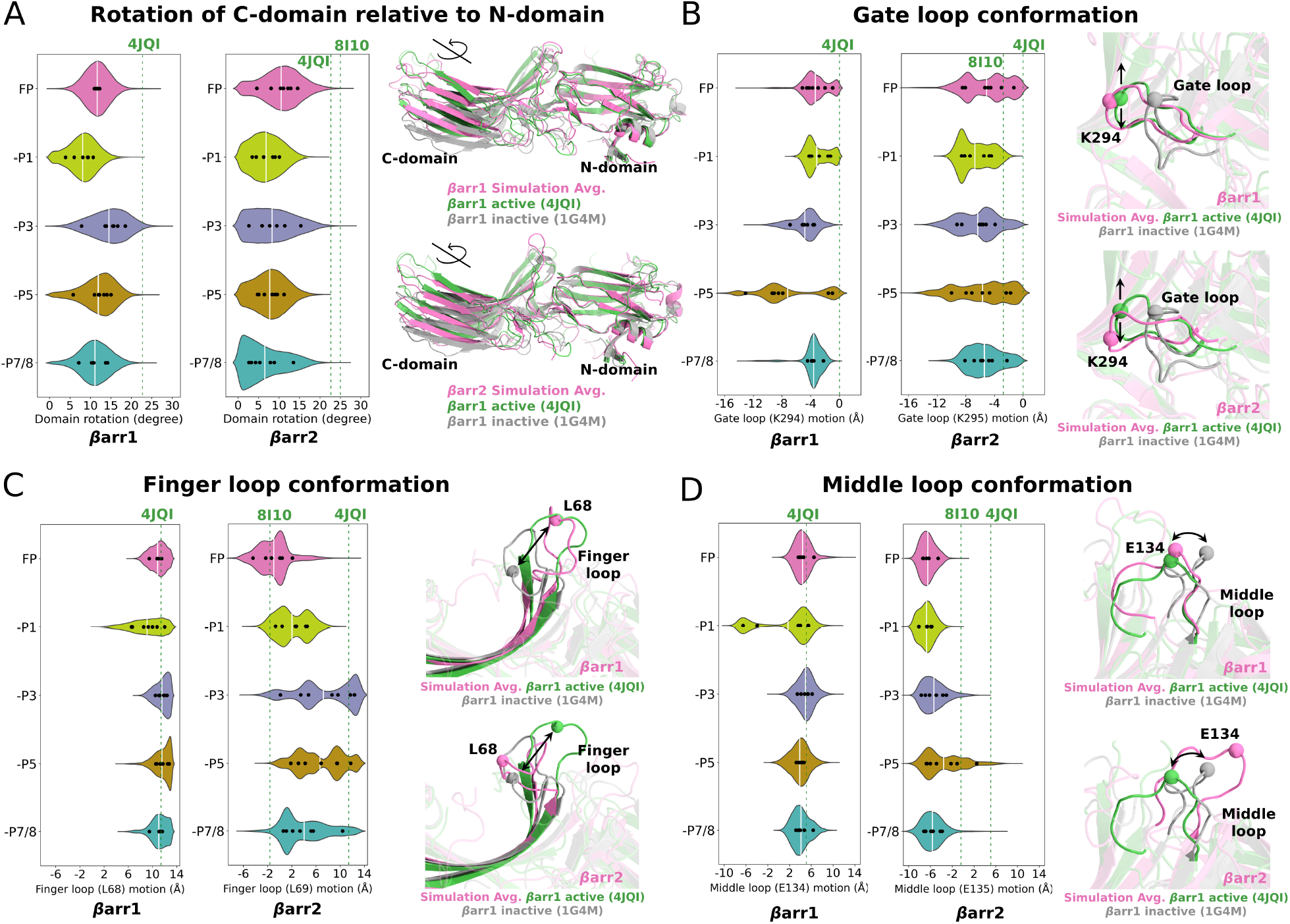
Conformational changes in *β*arr1 and *β*arr2 induced by distinct V2Rpp phospho-patterns. **(A)** Interdomain rotation as a measure of *β*arr activation. Violin plots show the distribution of domain rotation, with white lines representing mean values and black dots indicating individual trajectory averages. Cartoon representations illustrate domain rotations in experimental structures as well as in simulated *β*arr1_V2R(FP) and *β*arr2_V2R(FP) systems. Similarly, the conformational dynamics of critical structural motifs—**(B)** gate loop, **(C)** finger loop, and **(D)** middle loop—are represented using violin plots and cartoon depictions. The matrices used to quantify these motions are detailed in the SI Methods.

For *β*arr2, the –P1 and –P7/8 systems display reduced interdomain rotation, consistent with the trend observed in *β*arr1. Overall, all *β*arr2 variants exhibit lower interdomain rotation angles, with averages ranging from 6.5^*°*^ to 10.5^*°*^, the highest being in the FP system (Figure 3A). As previously discussed, this reduction is attributed to relatively weaker interactions between *β*arr2 and V2Rpp in the middle and distal regions (Figures S2A-S2D). Notably, residues T360/pT360 and pT359 in V2Rpp exhibit weaker interactions with K295 in *β*arr2 compared to K294 in *β*arr1 (Figure S2E). Unlike in *β*arr1, where the –P3 system exhibited the largest domain rotation, a similar trend is not observed in *β*arr2. This discrepancy is likely due to the inherent flexibility of the proximal and middle regions of V2Rpp, which resulted in poor engagement of pS357 with K295 in *β*arr2, thereby minimizing its influence on domain rotation (Figure S4).

Besides global changes, such as domain rotation, localized conformational differences in key structural motifs influence distinct arrestin conformations, ^20,24^ prompting our analysis of their motion in simulated systems. The dynamics of the gate loop were assessed by measuring the deviation of K294/K295 from *β*arr1 active conformation (PDB ID: 4JQI). Values near zero indicate alignment with the active state, while increasingly negative values reflect greater displacement, as described previously ^20^ and in the SI Methods. Both *β*arr1 and *β*arr2 exhibit a wide range of deviations in their distributions. However, *β*arr2 shows larger negative values, indicating greater gate loop mobility (Figure 3B). Interestingly, a similar variation was observed in a recently resolved cryo-EM structure of *β*arr2. ^22^ Strikingly, the –P5 systems in both isoforms show broader distributions, with pronounced gate loop fluctuations in the absence of T360 phosphorylation. This further underscores the stabilizing role of phosphorylation at T360 in maintaining gate loop conformation through interactions with K294/K295. It is important to note that distinct gate loop conformational changes have functional significance, as this region is involved in the binding of effectors such as MAP kinases, ^32^ RAPGEF3, ^33^ and PDE4D. ^34^

Next, we examined the conformational dynamics of the finger loop and the middle loop (Figure 1), which interact extensively with the receptor in the core complex, by analyzing the motions of their central residues, L68/L69 and E134/E135, as described in the SI Methods. The *β*arr1 systems exhibit finger loop motions comparable to those of the reference structure of active *β*arr1 (Figure 3C). A notable variation is observed in the –P1 system, which exhibits a broader distribution due to the absence of pT347 phosphate interactions (Figure S3; Table S1). In contrast, the motions are significantly larger in the *β*arr2 systems (Figure 3C), likely due to inherent fluctuations in the proximal V2Rpp, which lacks crucial finger loop interactions. Notably, a similar structural variation is observed in active *β*arr2 relative to *β*arr1. ^22^ Furthermore, the middle loop dynamics are correlated with those of the finger loop owing to their spatial proximity. We observed that the finger and middle loops form multiple contacts in *β*arr1, which are largely disrupted in *β*arr2 (Figure S5). As a result, the middle loop of *β*arr1 remains closer to the reference structure compared to *β*arr2 (Figure 3D). We also analyzed conformational variations in the C-loop and the back loop (Figure 1), regions important for calmodulin and RAF1 binding. ^35,36^ The C-loop exhibits significantly larger deviations in *β*arr1 than in *β*arr2, with concurrent changes observed in the back loop (Figure S6).

Furthermore, the dynamics of key structural motifs are largely uncorrelated and remain decoupled from global rotation, as indicated by their correlation values from mutual information (MI) analysis (Figure S7). Notably, a previous simulation study of *β*arr1 reported a similar trend, demonstrating that domain rotation can occur independently of other structural elements, such as the finger loop and interdomain crevice. ^20^

### Distinct phosphopeptides modulate effector-binding interfaces in *β*arr1 and *β*arr2

Allosteric signals originating from distal regions within a protein can induce localized conformational changes, including alterations in residue interactions, main-chain and side-chain dihedral angles (*φ, ψ*, and *χ*s), and solvent exposure. ^40–43^ These changes are typically assessed using metrics such as inter-residue contacts and interaction energies, solvent-accessible surface area (SASA), and information-theoretic or network-based approaches. ^44–52^ Recently, ML methods have been employed to predict allosteric sites and pathways by identifying key structural and dynamic features, offering improved accuracy and efficiency over traditional approaches. ^53–56^ To leverage ML for identifying allosteric sites modulated by differentially phosphorylated V2Rpp, we formulated a classification problem for each phosphorylation pattern. Class labels correspond to conformational clusters of *β*arr1 and *β*arr2, while input features represent the structural and energetic properties of individual residues. Because *β*arr conformational changes arise from V2Rpp interactions, accurate classification enables the identification of key residues influenced by specific phospho-patterns. Our approach involved two steps: (i) clustering *β*arr conformations to define class labels and (ii) applying ML classification with feature importance analysis to identify residues perturbed by each phospho-pattern, as detailed in SI Methods.

UMAP-based dimensionality reduction followed by *k*-means clustering identified 58 clusters within the *β*arr1 structural ensemble and 48 within *β*arr2, indicating greater conformational heterogeneity in *β*arr1 (Figure S8). This observation suggests that tail-only engagement of the receptor promotes a broader range of *β*arr1 conformations, potentially enabling more diverse effector interactions compared to *β*arr2. Moreover, clusters corresponding to different phospho-systems are largely separable (Figure 4A), consistent with prior findings that specific phosphorylation patterns drive distinct conformational states. ^20,24,26,30^

**Figure 4.**
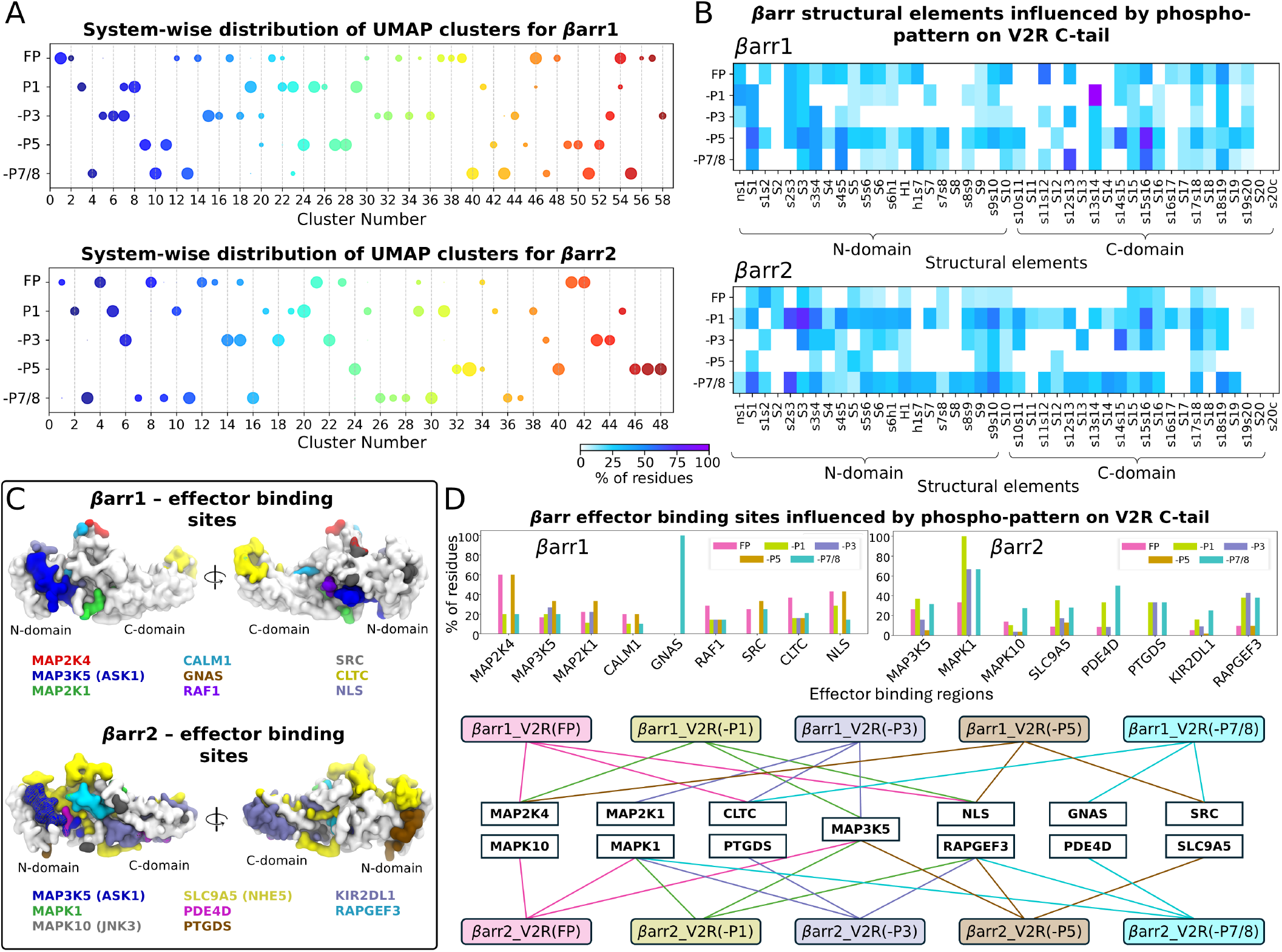
Identification of *β*arr residues/regions allosterically influenced by specific phospho-patterns on V2Rpp. **(A)** Populations of conformations within clusters formed by *β*arr1 and *β*arr2 systems. Cluster identities are used as labels for ML classification of each system to identify key residues affected by V2Rpp variants. **(B)** Heatmaps depicting the percentage of residues influenced in each structural element for *β*arr1 (top) and *β*arr2 (bottom). **(C)** Experimentally observed effector-binding regions in *β*arr1 (top) and *β*arr2 (bottom). ^24,37–39^ **(D)** Percentages of residues influenced in each effector-binding region, shown as bar plots for both *β*arr isoforms (top). The three most important regions are highlighted for clarity (bottom).

We further identified residues in *β*arr1 and *β*arr2 affected by V2Rpp phosphorylation using a two-phase ML approach. In the feature elimination phase, conformations corresponding to individual phospho-systems were classified using six ML algorithms: support vector machines (SVM), decision tree (DT), random forest (RF), extra trees (ET), extreme gradient boosting (XGB), and multilayer perceptron (MLP). Classification was performed separately for each feature set—dihedral angles, inter-residue interaction energies, and SASA (see SI Methods). After evaluating prediction performance (Figure S11), feature importance was assessed using local interpretable model-agnostic explanations (LIME), and features contributing to the top 99% of cumulative importance were retained (Figure S12). In the subsequent feature selection phase, the top-ranked features from each set were combined and classified using the best-performing models (MLP, ET, and XGB), followed by model evaluation (Figure S12). From these, the top 90% of important features—again determined via LIME—were extracted, and the corresponding residues were identified as significantly affected in each phospho-system. To evaluate how uniquely phosphorylated V2Rpp variants impact different structural elements of *β*arrs, we calculated the fraction of significant residues within each element (*β*-sheets, loops, and helices) relative to its total length (Figure 4B). These results are summarized in Table 1.

**Table 1:** Top five *β*arr structural elements most influenced by distinct V2Rpp phospho-patterns, ranked by the percentage of residues relative to their total length identified as important in ML-based classification of each system into conformational clusters.

| βarr1 | | | | | | | | | |
| --- | --- | --- | --- | --- | --- | --- | --- | --- | --- |
| FP |  | -P1 |  | -P3 |  | -P5 |  | -P7/8 |  |
| Structural element | % of res | Struct element | % of res | Struct element | % of res | Struct element | % of res | Struct element | % of res |
| s11s12<br>(c-edge loop1) | 57.1 | s13s14 | 100.0 | s3s4 | 42.9 | s15s16<br>(C-loop) | 80.0 | s12s13 | 66.7 |
| s18s19<br>(c-edge loop2) | 45.6 | ns1 | 42.9 | S1 | 40.0 | s14s15, S1 | 60.0 | s15s16<br>(C-loop) | 60.0 |
| s15s16<br>(C-loop) | 40.0 | S1 | 40.0 | s2s3, s12s13 | 33.3 | s4s5, S3 | 50.0 | s4s5 | 50.0 |
| S10 | 36.4 | s2s3 | 33.3 | s18s19<br>(c-edge loop2) | 27.3 | s3s4 | 42.9 | s3s4 | 42.9 |
| s2s3, s12s13, h1s7 | 33.3 | s18s19<br>(c-edge loop2) | 27.3 | S3, s13s14 | 25.0 | s9s10 | 36.4 | S1 | 40.0 |
| βarr2 | | | | | | | | | |
| Structural element | % of res | Struct element | % of res | Struct element | % of res | Struct element | % of res | Struct element | % of res |
| s1s2 | 40.0 | S3 | 75.0 | s14s15 | 60.0 | s9s10 | 27.3 | s2s3 | 66.7 |
| S3 | 25.0 | s2s3 | 66.7 | S3 | 50.0 | S5 | 25.0 | S1 | 60.0 |
| S15 | 21.4 | s15s16<br>(C-loop) | 60.0 | s15s16<br>(C-loop) | 40.0 | S1 | 20.0 | s9s10 | 54.6 |
| S2, S1, s15s16<br>(C-loop) | 20.0 | s3s4 | 57.1 | S17s18 | 35.7 | s5s6<br>(Finger loop) | 18.2 | s18s19<br>(c-edge loop2) | 50.0 |
| S5, S16 | 16.7 | s9s10 | 54.6 | S13 | 33.3 | S9 | 16.7 | S14 | 44.4 |

We found that the structural elements within the C-domain of *β*arr1 are predominantly influenced by differentially phosphorylated V2Rpp (Figure 4B, Table 1). Notably, the c-edge loop2 (s18s19) emerges as a key structural element in systems where distal V2Rpp phosphorylations (pT360, pS362, pS363, and pS364) are retained (FP, –P1, and –P3), highlighting the critical role of these distal sites in driving conformational changes in the c-edge loop2. This observation is consistent with recent biosensor-based findings. ^25^ Although the –P5 and –P7/8 systems also affect this loop, the impact is comparatively less pronounced (Figure 4B). The greater significance of c-edge loop2 in *β*arr1 aligns with previous findings showing that this loop mediates more effective membrane engagement in *β*arr1 relative to *β*arr2^25,57^ and plays a critical role in GPCR internalization. ^58^ In addition to c-edge loop2, other C-domain regions—specifically the C-loop (in FP, –P5, and –P7/8) and s12s13 (in FP, –P1, and –P7/8)—were identified as key sites (Table 1). Among these, the C-loop is particularly notable for its involvement in calmodulin binding. ^35^ Beyond the C-domain, elements of the N-domain such as S1 and s2s3 were also found to be important in *β*arr1 systems (Table 1), consistent with their previously established role in interacting with MAP3K5. ^59^

Unlike *β*arr1, the key structural elements in *β*arr2 are primarily concentrated within the N-domain (Figure 4B, Table 1), which serves as a scaffold for a range of downstream effectors, including MAP3K5, MAPK10 (JNK3), the Na^+^/H^+^ exchanger SLC9A5, phosphodiesterase PDE4D, and prostaglandin D2 synthase (PTGDS) (Table S10). In contrast, structural components within the C-domain—such as the C-loop, S15, and the s14s15 region—while still relevant, appear to play a less dominant role in *β*arr2 (Figure 4B). This distinction highlights a functional divergence between *β*arr1 and *β*arr2, consistent with their distinct physiological roles and mechanisms of action. ^8,25^

Inferring effector-binding sites influenced by phospho-patterns based solely on the proportion of residues within structural elements may overlook key functional subtleties, as not all residues contribute equally to effector interactions. Effector-binding sites often span residues from multiple regions (Figure 4C; Tables S9 and S10), ^24,37–39^ making the identification of key residues at each site essential for understanding their specific roles. In the fully phosphorylated *β*arr1-V2Rpp complex, we identified MAP2K4, clathrin, RAF1, MAP2K1, and the nuclear localization signal (NLS) as critical interaction sites (Figures 4D, S13, S14), emphasizing the importance of this pattern in mediating diverse cellular processes. Other *β*arr1-V2Rpp variants preferentially engage distinct sites: NLS is most affected in –P1, MAP3K5 and MAP2K1 in –P3, and MAP2K4 and SRC in –P5, consistent with structural and NMR data. ^24^ In *β*arr2, MAP kinases such as MAPK1, MAP3K5, and MAPK10 are preferentially targeted in the FP system, with MAPK1 (ERK2) emerging as the dominant site across all variants except –P5, where SLC9A5 is favored (Figure 4D, S13, S15). This deviation likely stems from the absence of phosphorylation at T360 in –P5, which compromises its interaction with K295 in the gate loop—a key site for MAPK1 binding. ^32^ Together, our results demonstrate the effectiveness of an ML-based approach in accurately identifying allosteric regions in *β*arrs and reveal how distinct phosphorylation patterns regulate their downstream signaling specificity.

### Phosphorylation directs distinct allosteric pathways connecting V2Rpp to distal functional sites

To investigate how differently phosphorylated V2Rpp influences distal functional regions in *β*arr1 and *β*arr2, we identified allosteric pathways using dynamic residue networks (DRNs), with edge weights learned via graph neural network-based autoencoders (GNN-AEs). GNN-AE inputs consisted of MD snapshots represented as unweighted graphs, where residues served as nodes and inter-residue contacts as edges. Node features included positional, energetic, and SASA metrics (see SI Methods). The encoder-decoder architecture employed message-passing operations to reconstruct input features with minimal error (Figure S16). The resulting edge weights, embedded in the latent space, were averaged across snapshots to generate a final network representation for each system. We then performed suboptimal path (SOP) analysis on the DRNs to characterize allosteric communication. Although the shortest path is often biologically meaningful, slightly longer SOPs of comparable length can also contribute to allosteric signaling. ^60^ SOPs were identified using a defined distance cutoff (Figure S17), and their number (*n*_SOP_) was used to quantify the strength of allosteric communication. ^60,61^

The c-edge loop2, known for its role in *β*arr membrane association, also serves as a clathrin binding site for *β*arr1. ^58^ Notably, it exhibits isoform-specific interactions with V2Rpp, as discussed in the previous section. In *β*arr1_V2R(FP), the strongest communication to c-edge loop2 originates from distal V2Rpp residues, relayed primarily through the lariat loop (D297, R282, F277); S15 (K232); S16 (M255); S18 (V328); and s19s20 (R363) in the C-domain (Figures 5A, B). Other systems, except –P1, involve largely the same structural elements, albeit with varying communication strengths (Figure S18). In –P1, the absence of phosphorylation at T347 markedly alters the communication network, redirecting the signal through s19s20 (R363, H362, P356); S19 (T350, F345); the back loop (L305); s12s13 (E206); and S18 (K322, K324, K326) (Figures 5B and S20). Involvement of the back loop is consistent with prior X-ray and NMR studies that reported significant conformational changes in this region for *β*arr1_V2R(–P1). ^24^ Interestingly, the allosteric strength is substantially reduced in –P7/8 (Figure S20), suggesting that distal phosphorylations at pS363 and pS364 are critical for driving allosteric changes in the c-edge loop2. In contrast, allosteric communication in *β*arr2 is primarily mediated by the lariat loop (L294, H296); S9 (C141); S15 (K233, Y239); S16 (Y250, Q256); and S18 (V329) in FP, with distinct pathways emerging under other phosphorylation patterns (Figures 5C, D, S20). Remarkably, overall communication strength is significantly lower in *β*arr2 than in *β*arr1 (Figures 5B, S20), consistent with the observed weaker coupling between the phosphopeptide and c-edge loop2 in *β*arr2. ^25^ This difference likely arises from fewer interactions between *β*arr2 residues and phosphate groups compared to *β*arr1 (Figure S2).

**Figure 5.**
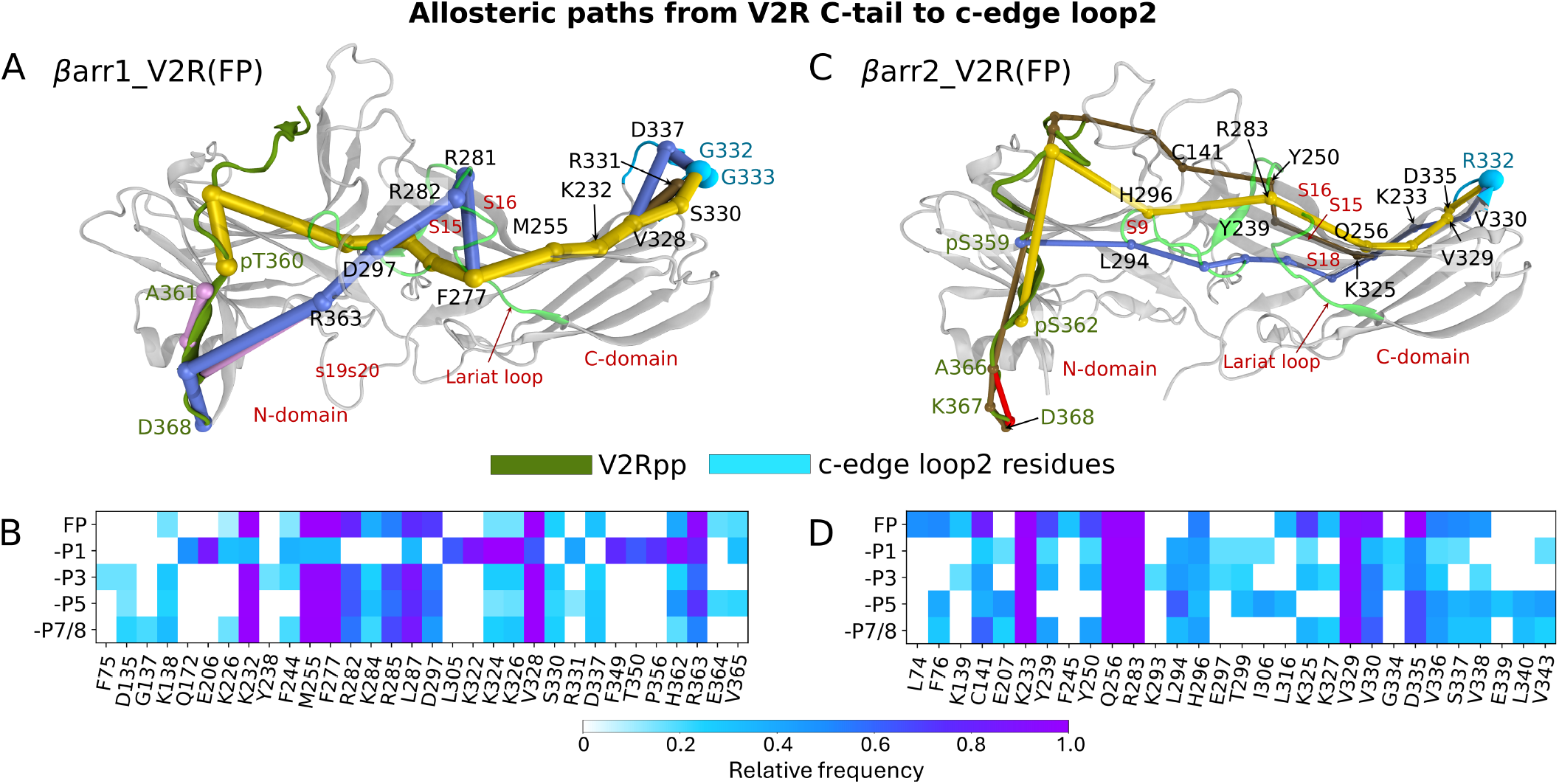
Allosteric communication pathways from V2Rpp to the c-edge loop2. The top five suboptimal paths, ranked by thickness, in the fully phosphorylated system and heatmaps of the top 20 residues, ranked by their occurrence in suboptimal paths for all systems, are displayed for **(A–B)** *β*arr1 and **(C-D)** *β*arr2. The cartoon representations are based on the fully phosphorylated structures (PDB IDs: 4JQI and 8I10), with missing residues modeled. Suboptimal paths are shown in yellow, blue, brown, red, and pink (note that some of them overlap at non-terminal residues). V2Rpp is shown in dark green, and c-edge loop2 in cyan. Key residues identified by a relative frequency *≥* 0.2 are highlighted in A and C.

SRC kinase, a key downstream effector of *β*arrs, plays an essential role in regulating diverse cellular processes, including growth, differentiation, migration, and survival. ^62,63^ Recent high-resolution structures have revealed that SRC binds to two distinct interfaces on *β*arr1: a polyproline site in the s6h1 loop (P88–P91) within the N-domain and a non-proline site in the central crest region; complex formation also depends on an additional polyproline motif (P120–P121) in the s7s8 loop. ^39^ Understanding how V2Rpp allosterically modulates SRC–*β*arr1 interactions is crucial for elucidating their functional implications. Allosteric pathways leading to the P88–P91 motif across all *β*arr1 phospho-variants primarily involve residues in H1 (L100, Q101, R103, I105), S7 (P114), and s19s20 (R363, E364, P366 (Figures 6A, 6B, S19A-S19D). Additionally, residue L48 in s4s5 is identified as an allosteric hub in the –P7/8 system. Interestingly, pathways to the central crest also involve s19s20 (Figures 6C, Figures 6D, S19E-S19H), highlighting the critical nature of this loop in modulating SRC-*β*arr1 interactions. Additional structural elements are engaged in other phospho-variants, including H1 (L100, Q101, R103) and S7 (P114, F115) in –P1; the lariat loop (R282, D297) in –P3; S6 (K77) in –P1, –P3, –P5, and –P7/8; s3s4 (D35) in –P1 and –P7/8; and hinge regions (Q172) in –P3 and –P7/8.

**Figure 6.**
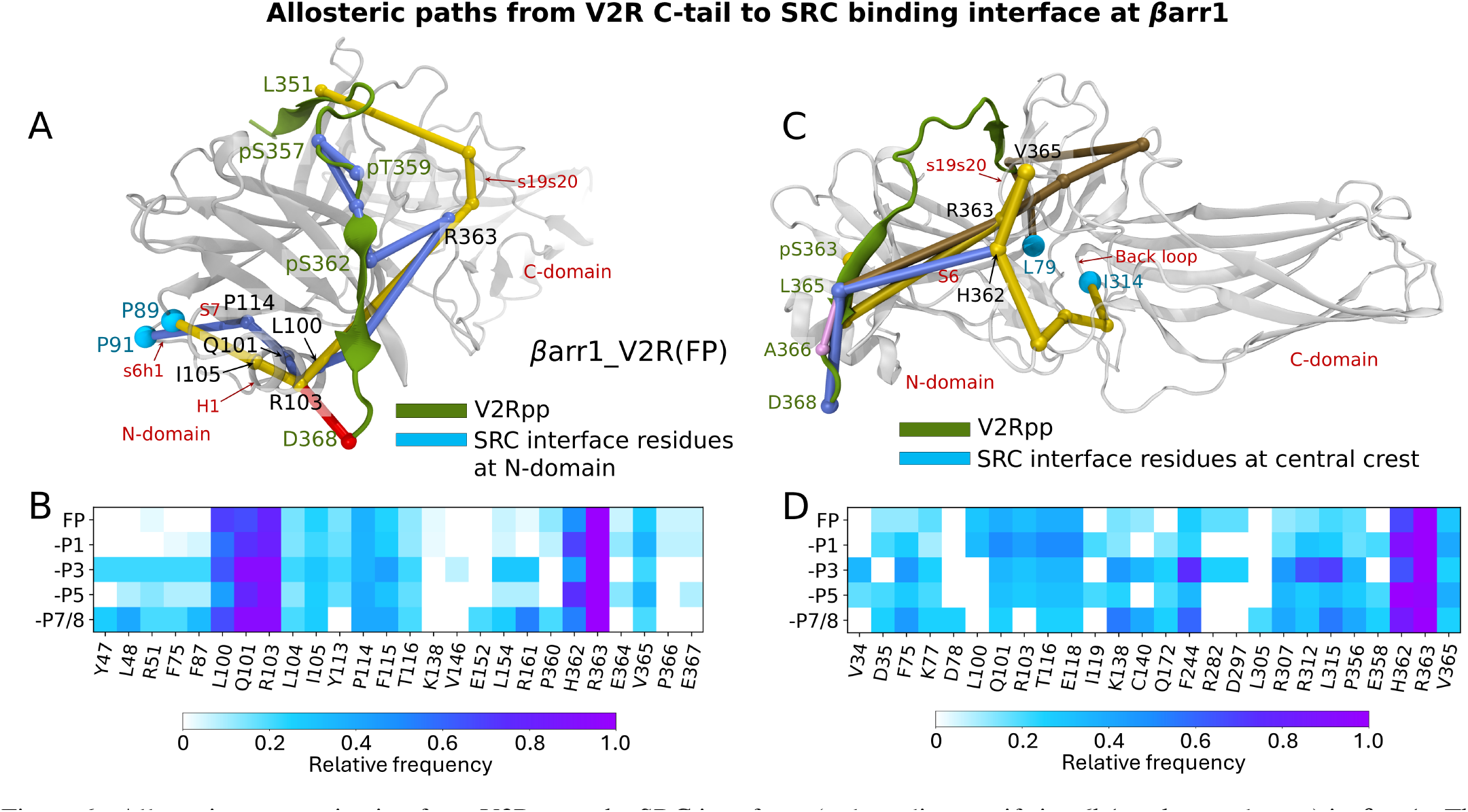
Allosteric communication from V2Rpp to the SRC interfaces (polyproline motifs in s6h1 and central crest) in *β*arr1. The top five suboptimal paths, ranked by thickness, in the fully phosphorylated system and heatmaps of the top 20 residues, ranked by their occurrence in suboptimal paths for all systems, are displayed for **(A–B)** the polyproline motifs in s6h1 and **(C–D)** the central crest. The cartoon representations are based on the fully phosphorylated structure (PDB ID: 4JQI), with missing residues modeled. Suboptimal paths are shown in yellow, blue, brown, red, and pink (note that some of them overlap at non-terminal residues). V2Rpp is shown in dark green, and SRC interface residues are shown in cyan. Key residues identified by a relative frequency*≥* 0.2 are highlighted in A and C.

We examined allosteric communication from V2Rpp to key effector-binding sites distal to the phosphopeptide interface of *β*arr1. The results, which highlight critical residues and structural elements involved in these communications, are presented in Figures S20 and S21. The binding site of the RAF1 protein, a key member of the RAS/MAPK pathway that regulates cell growth and differentiation, lies in the back loop of *β*arr1^36^ and is allosterically connected through residues in the lariat loop, C-loop, hinge region, and s19s20 (Figure S20). MAP2K1 (MEK1), enzymatically activated by RAF1 and known to interact with *β*-sheets S3 and S19, ^64,65^ primarily engages s19s20, S6, the C-loop, and the back loop (Figure S21). Notably, a marked reduction in allosteric communication strength is observed when phosphorylation at the middle or distal sites (P3, P5, or P7/8) is absent, suggesting that these specific phosphate groups collectively contribute to *β*arr1-MAP2K1 complex formation.

In the case of *β*arr2, the killer cell immunoglobulin-like receptor 2DL1 (KIR2DL1), which prevents cell lysis in the human immune response, interacts with distal regions of the C-domain, particularly *β*-sheets S11–S15 and their connecting loop. ^66^ V2Rpp is allosterically connected to these regions primarily through multiple *β*-sheets in the N-domain (S6 and S9) and the C-domain (S16, S18, and S19) (Figure S22). Notably, phospho-pattern-specific differences are observed, including involvement of the lariat loop in –P1, –P3, and –P7/8; the back loop in –P5 and –P7/8; and the C-loop in –P5. The pathways leading to the binding site of cAMP-degrading phosphodiesterase PDE4D at *β*-sheet S14^34^ involve multiple structural elements, including *β*-sheets (S6, S9, S16, S18, and S19) and key loops (the lariat loop, C-loop, and back loop) in the FP system (Figure S23). However, in other variants, the contributions of several structural elements are notably reduced. Specifically, N-domain *β*-sheets are absent in –P1, S9 is not involved in –P3 and –P5, and S6 is absent in –P7/8. Interestingly, –P3 engages a larger number of residues from the lariat loop than the other variants. Furthermore, the binding site of prostaglandin-H2 D-isomerase (PTGDS), located in the N-domain at S6, s6h1, and H1, ^67^ is primarily connected via S2 and S7 in the FP and –P7/8 systems, while S2 is involved in –P1 and –P3, and S7 in –P5 (Figure S24).

## Discussion

The integration of MD simulations, machine learning, and graph neural networks allowed us to dissect the distinct functional dynamics of *β*arrs upon V2Rpp binding. Our findings reveal that *β*arr1 forms a more stable complex with V2Rpp than *β*arr2, with differences in phosphate interactions indicating isoform-specific allosteric triggers. Phosphorylation patterns on V2Rpp drive different conformational changes in *β*arr1 and *β*arr2, influencing unique effector-binding interfaces. Key structural elements mediate this allosteric communication, with the specific phospho-pattern dictating effector engagement.

The reduced binding stability of V2Rpp to *β*arr2 compared to *β*arr1 is consistent with prior experimental evidence showing that *β*arr1 forms a more stable ‘hanging complex’ with GPCRs. ^25^ Structural investigations of *β*arr1 have identified unique allosteric triggers induced by specific phospho-patterns on the bound V2Rpp. ^24^ Our results not only corroborate these findings but also extend them to *β*arr2. The reduced interdomain rotation observed in *β*arr2, relative to *β*arr1, while not an inherent feature of *β*arr2, ^29^ is likely driven by interactions with the specific phospho-patterns examined here. Distinct interdomain angles induced by specific phosphopeptides in *β*arr1 and *β*arr2 give rise to unique active conformations, supporting the hypothesis that *β*arrs undergo pattern-dependent interdomain rotations to selectively engage downstream effectors. ^68–71^ Additionally, *β*arrs may achieve this selectivity through conformational variations in key structural motifs, as observed in our analysis. Indeed, a previous study using unnatural amino acid incorporation and ^19^F NMR demonstrated that the movements of the finger and middle loops are modulated by *β*arr1 interactions with different phospho-patterns, thereby stabilizing distinct structural states linked to specific arrestin functions. ^72^ Moreover, changes in finger loop orientation have been correlated with structural variations in arrestin-1, resulting in distinct domain rotations, ^73^ while interactions between the gate loop and specific phosphate groups in V2Rpp have been shown to regulate the extent of *β*arr1 activation. ^20^

The ML-based approach employed in this study enabled a direct and systematic comparison between *β*arr1 and *β*arr2 by identifying regions that are allosterically influenced by similar phosphorylation patterns. Notably, our analysis provides a structural explanation for differences in their C-domains that had previously been suggested by spectroscopic studies, ^8,25^ by pinpointing the specific elements responsible for these distinctions. We also identified structural regions uniquely affected by phosphorylation in each isoform, many of which involve residues known to interact with effectors. With the increasing availability of high-resolution structures of *β*arr–effector complexes, our ML framework could be further applied to explore effector-specific conformational dynamics in *β*arr1 and *β*arr2.

Beyond their canonical scaffolding roles, *β*arrs are known to allosterically activate downstream kinases, including SRC, ^63^ RAF1, ^74^ and ERK2. ^75^ Recent studies have provided critical insights into phospho-pattern-specific effector binding ^24,26,30^ and shown how bound phosphopeptides modulate distal regions of *β*arrs in an isoform-specific manner. ^25^ Moreover, simulations combined with experimental validations have elucidated how receptor core and tail engagement influence arrestin activation, ^76^ and how distinct phosphorylation patterns affect binding stability, activation, and conformational dynamics, particularly in *β*arr1. ^20^ The present study advances our understanding by offering residue-level insights into how diverse phosphorylation patterns in bound phosphopeptides drive allosteric communication to distal functional regions in both *β*arr1 and *β*arr2. Notably, the observed attenuation of allosteric signaling to c-edge loop 2 in *β*arr2 relative to *β*arr1, is consistent with prior experimental findings. ^25^ Furthermore, the distinct allosteric responses elicited by identical phosphorylation patterns underscore the known functional divergence between the isoforms. ^8^

Mapping signaling pathways to the recently identified SRC interfaces—located at the N-domain polyproline site and the central crest region— ^39^ highlights a more nuanced mode of regulation than previously appreciated. ^62^ Our data suggest that distinct phosphorylation patterns can modulate SRC activation to varying degrees, potentially contributing to context-specific signaling outcomes. Moreover, the varied engagement of structural elements across phospho-variants in communicating with effector interfaces suggests that these motifs act as molecular switches, selectively triggered by specific phospho-barcodes. Together, these findings emphasize the adaptability of *β*arrs as scaffolds and allosteric regulators of GPCR signaling, with phosphorylation-dependent conformational changes driving their functional plasticity.

Overall, our residue-level insights provide a useful basis for developing strategies to selectively modulate *β*-arrestin-mediated signaling downstream of GPCRs. This knowledge may aid in the design of biased ligands that preferentially activate specific downstream pathways or support efforts to engineer targeted mutations that disrupt selective effector interactions. Future studies investigating how distinct phosphorylation patterns in the receptor C-tail coordinate with receptor-core engagement in *β*arr1 and *β*arr2 may uncover additional regulatory mechanisms that fine-tune arrestin function.

## Methods

### System preparation and molecular dynamics

Initial coordinates for the *β*arr1 and *β*arr2 systems were obtained from active-state structures resolved via X-ray crystallography and cryo-EM. Both systems were solvated and ionized using CHARMM-GUI, with *β*arr1 comprising approximately 96,000 atoms and *β*arr2 approximately 106,000 atoms. Molecular dynamics simulations were performed using GROMACS, with equilibration in the NVT and NPT ensembles followed by 1 *µ*s production runs, yielding a cumulative 70 *µ*s of simulation data. Temperature and pressure were maintained using standard protocols, with a 2 fs integration time step and trajectory frames saved every 10 ps. Detailed procedures for system preparation and simulation are provided in the SI Methods.

### Analysis of trajectories and ML-based methods

Root mean square deviation (RMSD) values, distances, and dihedral angles were calculated using the CPPTRAJ module of AMBER18^77^ after converting trajectories to a compatible format using the MDConvert tool from MDTraj. ^78^ Stability calculations for V2Rpp binding were performed according to established protocols. ^20^ Inter-residue interactions, mutual information, and conformational dynamics (interdomain rotation angles and motions of key structural motifs) were analyzed using custom Python scripts. Machine-learning-based approaches, including dimensionality reduction, clustering, feature importance analysis, and allosteric pathway identification via graph neural networks coupled with suboptimal path analysis, were applied to interpret dynamic data. Detailed methodological descriptions are provided in the SI Methods.

## Data Availability

All the essential input files required to reproduce the simulations and ML analysis codes are available at https://github.com/rkmlabiiserb/barr_phos.

## Acknowledgment

M.K.M. thanks Kunal Shewani for his assistance in the initial stage of simulations. R.K.M. acknowledges the financial support from the Science and Engineering Research Board (SERB), Department of Science and Technology, India (file no. CRG/2023/000970). The support and the resources provided by PARAM Sanganak under the National Supercomputing Mission, Government of India, at the Indian Institute of Technology, Kanpur, are gratefully acknowledged.

## Notes

The authors declare no competing interests.

## Notes

### Competing Interest Statement

The authors have declared no competing interest.

## References

[1] Wess, J.; Oteng, A.-B.; Rivera-Gonzalez, O.; Gurevich, E. V.; Gurevich, V. V. Pharmacological Reviews 2023, 75, 854–884.

[2] Gurevich, V. V.; Gurevich, E. V. Cellular and Molecular Life Sciences 2019, 76, 4413–4421.

[3] Dror, R. O.; Arlow, D. H.; Maragakis, P.; Mildorf, T. J.; Pan, A. C.; Xu, H.; Borhani, D. W.; Shaw, D. E. Proceedings of the National Academy of Sciences U.S.A 2011, 108, 18684– 18689.

[4] Alhadeff, R.; Vorobyov, I.; Yoon, H. W.; Warshel, A. Proceedings of the National Academy of Sciences U.S.A 2018, 115, 10327–10332.

[5] Mafi, A.; Kim, S.-K.; Goddard III, W. A. Proceedings of the National Academy of Sciences U.S.A 2022, 119, e2110085119.

[6] Komolov, K. E.; Benovic, J. L. Cellular Signalling 2018, 41, 17–24.

[7] Peterson, Y. K.; Luttrell, L. M. Pharmacological Reviews 2017, 69, 256–297.

[8] Ghosh, E.; Dwivedi, H.; Baidya, M.; Srivastava, A.; Kumari, P.; Stepniewski, T.; Kim, H. R.; Lee, M.-H.; van Gastel, J.; Chaturvedi, M.; others Cell Reports 2019, 28, 3287–3299.

[9] Tobin, A. British Journal of Pharmacology 2008, 153, S167– S176.

[10] Nobles, K. N.; Xiao, K.; Ahn, S.; Shukla, A. K.; Lam, C. M.; Rajagopal, S.; Strachan, R. T.; Huang, T.-Y.; Bressler, E. A.; Hara, M. R.; others Science Signaling 2011, 4, ra51–ra51.

[11] Butcher, A. J.; Prihandoko, R.; Kong, K. C.; McWilliams, P.; Edwards, J. M.; Bottrill, A.; Mistry, S.; Tobin, A. B. Journal of Biological Chemistry 2011, 286, 11506–11518.

[12] Thomsen, A. R.; Plouffe, B.; Cahill, T. J.; Shukla, A. K.; Tarrasch, J. T.; Dosey, A. M.; Kahsai, A. W.; Strachan, R. T.; Pani, B.; Mahoney, J. P.; others Cell 2016, 166, 907–919.

[13] Kang, Y.; Zhou, X. E.; Gao, X.; He, Y.; Liu, W.; Ishchenko, A.; Barty, A.; White, T. A.; Yefanov, O.; Han, G. W.; others Nature 2015, 523, 561–567.

[14] Staus, D. P.; Hu, H.; Robertson, M. J.; Kleinhenz, A. L.; Wingler, L. M.; Capel, W. D.; Latorraca, N. R.; Lefkowitz, R. J.; Skiniotis, G. Nature 2020, 579, 297–302.

[15] Kumari, P.; Srivastava, A.; Banerjee, R.; Ghosh, E.; Gupta, P.; Ranjan, R.; Chen, X.; Gupta, B.; Gupta, C.; Jaiman, D.; others Nature Communications 2016, 7, 13416.

[16] Kumari, P.; Srivastava, A.; Ghosh, E.; Ranjan, R.; Dogra, S.; Yadav, P. N.; Shukla, A. K. Molecular Biology of the Cell 2017, 28, 1003–1010.

[17] Gurevich, V. V.; Gurevich, E. V. Trends in Pharmacological Sciences 2024, 45, 639–650.

[18] Mayer, D.; Damberger, F. F.; Samarasimhareddy, M.; Feldmueller, M.; Vuckovic, Z.; Flock, T.; Bauer, B.; Mutt, E.; Zosel, F.; Allain, F. H.; others Nature Communications 2019, 10, 1261.

[19] Maharana, J.; Banerjee, R.; Yadav, M. K.; Sarma, P.; Shukla, A. K. Current Opinion in Structural Biology 2022, 75, 102406.

[20] Latorraca, N. R.; Masureel, M.; Hollingsworth, S. A.; Heydenreich, F. M.; Suomivuori, C.-M.; Brinton, C.; Townshend, R. J.; Bouvier, M.; Kobilka, B. K.; Dror, R. O. Cell 2020, 183, 1813–1825.

[21] Zhou, X. E.; He, Y.; de Waal, P. W.; Gao, X.; Kang, Y.; Van Eps, N.; Yin, Y.; Pal, K.; Goswami, D.; White, T. A.; others Cell 2017, 170, 457–469.

[22] Maharana, J.; Sarma, P.; Yadav, M. K.; Saha, S.; Singh, V.; Saha, S.; Chami, M.; Banerjee, R.; Shukla, A. K. Molecular Cell 2023, 83, 2091–2107.

[23] Isaikina, P.; Petrovic, I.; Jakob, R. P.; Sarma, P.; Ranjan, A.; Baruah, M.; Panwalkar, V.; Maier, T.; Shukla, A. K.; Grzesiek, S. Molecular Cell 2023, 83, 2108–2121.

[24] He, Q.-T.; Xiao, P.; Huang, S.-M.; Jia, Y.-L.; Zhu, Z.-L.; Lin, J.-Y.; Yang, F.; Tao, X.-N.; Zhao, R.-J.; Gao, F.-Y.; others Nature Communications 2021, 12, 2396.

[25] Haider, R. S.; Matthees, E. S.; Drube, J.; Reichel, M.; Zabel, U.; Inoue, A.; Chevigné, A.; Krasel, C.; Deupi, X.; Hoffmann, C. Nature Communications 2022, 13, 5638.

[26] Gareri, C.; Pfeiffer, C. T.; Jiang, X.; Paulo, J. A.; Gygi, S. P.; Pham, U.; Chundi, A.; Wingler, L. M.; Staus, D. P.; Stepniewski, T. M.; others Science Signaling 2024, 17, eadk5736.

[27] Shukla, A. K.; Manglik, A.; Kruse, A. C.; Xiao, K.; Reis, R. I.; Tseng, W.-C.; Staus, D. P.; Hilger, D.; Uysal, S.; Huang, L.-Y.; others Nature 2013, 497, 137–141.

[28] Herrera, L. P. T.; Andreassen, S. N.; Caroli, J.; Rodríguez-Espigares, I.; Kermani, A. A.; Keserű, G. M.; Kooistra, A. J.; Pándy-Szekeres, G.; Gloriam, D. E. Nucleic Acids Research 2025, 53, D425–D435.

[29] Min, K.; Yoon, H.-J.; Park, J. Y.; Baidya, M.; Dwivedi-Agnihotri, H.; Maharana, J.; Chaturvedi, M.; Chung, K. Y.; Shukla, A. K.; Lee, H. H. Structure 2020, 28, 1014–1023.

[30] Dwivedi-Agnihotri, H.; Chaturvedi, M.; Baidya, M.; Stepniewski, T. M.; Pandey, S.; Maharana, J.; Srivastava, A.; Caengprasath, N.; Hanyaloglu, A. C.; Selent, J.; others Science Advances 2020, 6, eabb8368.

[31] Vishnivetskiy, S. A.; Huh, E. K.; Gurevich, E. V.; Gurevich, V. V. Journal of Neurochemistry 2021, 157, 1138–1152.

[32] Xu, T.-R.; Baillie, G. S.; Bhari, N.; Houslay, T. M.; Pitt, A. M.; Adams, D. R.; Kolch, W.; Houslay, M. D.; Milligan, G. Biochemical Journal 2008, 413, 51–60.

[33] Berthouze-Duquesnes, M.; Lucas, A.; Saulière, A.; Sin, Y. Y.; Laurent, A.-C.; Galés, C.; Baillie, G.; Lezoualc’h, F. Cellular Signalling 2013, 25, 970–980.

[34] Baillie, G. S.; Adams, D. R.; Bhari, N.; Houslay, T. M.; Vadrevu, S.; Meng, D.; Li, X.; Dunlop, A.; Milligan, G.; Bolger, G. B.; others Biochemical Journal 2007, 404, 71–80.

[35] Wu, N.; Hanson, S. M.; Francis, D. J.; Vishnivetskiy, S. A.; Thibonnier, M.; Klug, C. S.; Shoham, M.; Gurevich, V. V. Journal of Molecular Biology 2006, 364, 955–963.

[36] Coffa, S.; Breitman, M.; Spiller, B. W.; Gurevich, V. V. Biochemistry 2011, 50, 6951–6958.

[37] Xiao, K.; McClatchy, D. B.; Shukla, A. K.; Zhao, Y.; Chen, M.; Shenoy, S. K.; Yates III, J. R.; Lefkowitz, R. J. Proceedings of the National Academy of Sciences U.S.A 2007, 104, 12011–12016.

[38] Crépieux, P.; Poupon, A.; Langonné-Gallay, N.; Reiter, E.; Delgado, J.; Schaefer, M. H.; Bourquard, T.; Serrano, L.; Kiel, C. Frontiers in Endocrinology 2017, 8, 32.

[39] Pakharukova, N.; Thomas, B. N.; Bansia, H.; Li, L.; Abzalimov, R. R.; Kim, J.; Kahsai, A. W.; Pani, B.; Bassford, D. K.; Liu, S.; others BioRxiv preprint bioRxiv: 10.1101/2024.07.31.605623 2024,

[40] Buchenberg, S.; Sittel, F.; Stock, G. Proceedings of the National Academy of Sciences U.S.A 2017, 114, E6804–E6811.

[41] Wu, N.; Barahona, M.; Yaliraki, S. N. Current Opinion in Structural Biology 2024, 84, 102737.

[42] Guo, J.; Zhou, H.-X. Chemical Reviews 2016, 116, 6503–6515.

[43] Porter, J. R.; Moeder, K. E.; Sibbald, C. A.; Zimmerman, M. I.; Hart, K. M.; Greenberg, M. J.; Bowman, G. R. Biophysical Journal 2019, 116, 818–830.

[44] Doshi, U.; Holliday, M. J.; Eisenmesser, E. Z.; Hamelberg, D. Proceedings of the National Academy of Sciences U.S.A 2016, 113, 4735–4740.

[45] Vijayabaskar, M.; Vishveshwara, S. Biophysical Journal 2010, 99, 3704–3715.

[46] Sethi, A.; Eargle, J.; Black, A. A.; Luthey-Schulten, Z. Proceedings of the National Academy of Sciences U.S.A 2009, 106, 6620–6625.

[47] Hacisuleyman, A.; Erman, B. Proteins: Structure, Function, and Bioinformatics 2017, 85, 1056–1064.

[48] Madhu, M. K.; Debroy, A.; Murarka, R. K. The Journal of Physical Chemistry B 2022, 126, 1917–1932.

[49] Madhu, M. K.; Shewani, K.; Murarka, R. K. Journal of Chemical Information and Modeling 2024, 64, 449–469.

[50] Miao, Y.; Caliman, A. D.; McCammon, J. A. Biophysical Journal 2015, 108, 1796–1806.

[51] Liu, J.; Nussinov, R. Proceedings of the National Academy of Sciences U.S.A 2008, 105, 901–906.

[52] Clark, L. J.; Krieger, J.; White, A. D.; Bondarenko, V.; Lei, S.; Fang, F.; Lee, J. Y.; Doruker, P.; Böttke, T.; Jean-Alphonse, F.; others Nature Chemical Biology 2020, 16, 1096–1104.

[53] Agajanian, S.; Alshahrani, M.; Bai, F.; Tao, P.; Verkhivker, G. M. Journal of Chemical Information and Modeling 2023, 63, 1413–1428.

[54] Nerín-Fonz, F.; Cournia, Z. Current Opinion in Structural Biology 2024, 85, 102774.

[55] Ferraro, M.; Moroni, E.; Ippoliti, E.; Rinaldi, S.; Sanchez-Martin, C.; Rasola, A.; Pavarino, L. F.; Colombo, G. The Journal of Physical Chemistry B 2020, 125, 101–114.

[56] Zhu, J.; Wang, J.; Han, W.; Xu, D. Nature Communications 2022, 13, 1661.

[57] Lally, C. C. M.; Bauer, B.; Selent, J.; Sommer, M. E. Nature Communications 2017, 8, 14258.

[58] Kang, D. S.; Kern, R. C.; Puthenveedu, M. A.; von Zastrow, M.; Williams, J. C.; Benovic, J. L. Journal of Biological Chemistry 2009, 284, 29860–29872.

[59] Li, X.; MacLeod, R.; Dunlop, A. J.; Edwards, H. V.; Advant, N.; Gibson, L. C.; Devine, N. M.; Brown, K. M.; Adams, D. R.; Houslay, M. D.; others FEBS Letters 2009, 583, 3310–3316.

[60] Van Wart, A. T.; Durrant, J.; Votapka, L.; Amaro, R. E. Journal of Chemical Theory and Computation 2014, 10, 511–517.

[61] Eargle, J.; Luthey-Schulten, Z. Bioinformatics 2012, 28, 3000– 3001.

[62] Yang, F.; Xiao, P.; Qu, C.-x.; Liu, Q.; Wang, L.-y.; Liu, Z.-x.; He, Q.-t.; Liu, C.; Xu, J.-y.; Li, R.-r.; others Nature Chemical Biology 2018, 14, 876–886.

[63] Pakharukova, N.; Masoudi, A.; Pani, B.; Staus, D. P.; Lefkowitz, R. J. Journal of Biological Chemistry 2020, 295, 16773–16784.

[64] Meng, D.; Lynch, M. J.; Huston, E.; Beyermann, M.; Eichhorst, J.; Adams, D. R.; Klussmann, E.; Houslay, M. D.; Baillie, G. S. Journal of Biological Chemistry 2009, 284, 11425–11435.

[65] Cassier, E.; Gallay, N.; Bourquard, T.; Claeysen, S.; Bockaert, J.; Crepieux, P.; Poupon, A.; Reiter, E.; Marin, P.; Vandermoere, F. Elife 2017, 6, e23777.

[66] Yu, M.-C.; Su, L.-L.; Zou, L.; Liu, Y.; Wu, N.; Kong, L.; Zhuang, Z.-H.; Sun, L.; Liu, H.-P.; Hu, J.-H.; others Nature Immunology 2008, 9, 898–907.

[67] Mathurin, K.; Gallant, M. A.; Germain, P.; Allard-Chamard, H.; Brisson, J.; Iorio-Morin, C.; de Brum Fernandes, A.; Caron, M. G.; Laporte, S. A.; Parent, J.-L. Journal of Biological Chemistry 2011, 286, 2696–2706.

[68] Chen, Q.; Iverson, T. M.; Gurevich, V. V. Trends in Biochemical Sciences 2018, 43, 412–423.

[69] Ahmed, M. R.; Zhan, X.; Song, X.; Kook, S.; Gurevich, V. V.; Gurevich, E. V. Biochemistry 2011, 50, 3749–3763.

[70] Song, X.; Coffa, S.; Fu, H.; Gurevich, V. V. Journal of Biological Chemistry 2009, 284, 685–695.

[71] Zhan, X.; Perez, A.; Gimenez, L. E.; Vishnivetskiy, S. A.; Gurevich, V. V. Cellular Signalling 2014, 26, 766–776.

[72] Yang, F.; Yu, X.; Liu, C.; Qu, C.-X.; Gong, Z.; Liu, H.-D.; Li, F.-H.; Wang, H.-M.; He, D.-F.; Yi, F.; others Nature Communications 2015, 6, 8202.

[73] Zhuang, T.; Chen, Q.; Cho, M.-K.; Vishnivetskiy, S. A.; Iverson, T. M.; Gurevich, V. V.; Sanders, C. R. Proceedings of the National Academy of Sciences U.S.A 2013, 110, 942–947.

[74] Zang, Y.; Kahsai, A. W.; Pakharukova, N.; Huang, L.-y.; Lefkowitz, R. J. Journal of Biological Chemistry 2021, 297.

[75] Kahsai, A. W.; Shah, K. S.; Shim, P. J.; Lee, M. A.; Shreiber, B. N.; Schwalb, A. M.; Zhang, X.; Kwon, H. Y.; Huang, L.-Y.; Soderblom, E. J.; others Proceedings of the National Academy of Sciences U.S.A 2023, 120, e2303794120.

[76] Latorraca, N. R.; Wang, J. K.; Bauer, B.; Townshend, R. J.; Hollingsworth, S. A.; Olivieri, J. E.; Xu, H. E.; Sommer, M. E.; Dror, R. O. Nature 2018, 557, 452–456.

[77] Roe, D. R.; Cheatham III, T. E. Journal of Chemical Theory and Computation 2013, 9, 3084–3095.

[78] McGibbon, R. T.; Beauchamp, K. A.; Harrigan, M. P.; Klein, C.; Swails, J. M.; Hernández, C. X.; Schwantes, C. R.; Wang, L.-P.; Lane, T. J.; Pande, V. S. Biophysical Journal 2015, 109, 1528–1532.

