## Supplemental Information for "Mechanistic basis for GPCR phosphorylation-dependent allosteric signaling specificity of *β*-arrestin 1 and 2"

---

#### Supplementary Methods

##### Model preparation and system setup

The initial coordinates for the $\beta$arr1 systems (residues 5 to 367) were derived from active-state X-ray crystal structures with bound phosphopeptide (V2Rpp) having distinct phosphorylation patterns. The fully phosphorylated $\beta$arr1 system ($\beta$arr1_V2R(FP)), modeled from PDB ID 4JQI, contained phosphorylation at all serine and threonine residues of V2Rpp (pS347, pS350, pS357, pT359, pY360, pS362, pS363, and pS364), labeled as P1 through P7. Other systems, with no phosphorylation at one or two specific sites, were modeled from PDB IDs 7DF9 ($\beta$arr1_V2R(–P1), lacking phosphorylation at P1), 7DFC ($\beta$arr1_V2R(–P3), lacking phosphorylation at P3), 7DFA ($\beta$arr1_V2R(–P5), lacking phosphorylation at P5), and 7DFB ($\beta$arr1_V2R(–P7/8), lacking phosphorylation at P7 and P8). For the $\beta$arr2 systems (residues 7 to 350), initial coordinates were derived from a cryo-EM structure (PDB ID 8I10). However, the residues 346–357 in the proximal part of the phosphopeptide were unresolved in the structure, necessitating their modeling to resemble the $\beta$arr1–V2Rpp systems. The $\beta$arr2 systems were designated as $\beta$arr2_V2R(FP), $\beta$arr2_V2R(–P1), $\beta$arr2_V2R(–P3, $\beta$arr2_V2R(–P5), $\beta$arr2_V2R(–P7/8). MODELLER v9.17 was used to add the missing residues and prepare the final models.

All systems were prepared using CHARMM36m force field for proteins and solvated with the CHARMM-modified TIP3P water model via the CHARMM-GUI web server. To create a physiologically relevant environment, sodium ($Na^+$) and chloride ($Cl^-$) ions were added to achieve an ionic concentration of 150 mM. The resulting systems contained, on average, 96,388 atoms and measured an initial volume of $\sim$1030 nm$^3$ for $\beta$arr1, and on average, 105,969 atoms with an initial volume of $\sim$1130 nm$^3$ for $\beta$arr2.

##### Molecular dynamics simulations details

Molecular dynamics (MD) simulations were performed using GROMACS version 2019.5. Each system underwent energy minimization for 5000 steps using the steepest descent algorithm to relax the structure before further processing. Subsequently, the temperature of each system was gradually increased from 0 K to 310 K over a duration of 125 ps in the NVT ensemble, with positional restraints applied to the heavy atoms of the proteins. This was followed by a 100 ns NPT equilibration phase, allowing the system to reach equilibrium without restraints. Production runs were carried out under the NPT ensemble at a constant temperature of 310 K using a Nose-Hoover thermostat with a coupling constant of 1 ps. The pressure was maintained at 1 bar using a Parrinello-Rahman barostat with a coupling constant of 5 ps. Long-range electrostatic interactions were calculated via the particle mesh Ewald (PME) method with a grid spacing of 1 Å. A cutoff of 12 Å was applied for both the real space electrostatic interactions and Lennard-Jones potential for van der Waals interactions, with a force switch applied at 10 Å. The LINCS algorithm was employed to constrain bond lengths involving hydrogen atoms. A time step of 2 fs was utilized throughout the simulations and trajectory frames were saved every 10ps. Seven independent MD trajectories, each lasting 1 $\mu$s, were generated for all systems, resulting in a total of 70 $\mu$s of simulation data.

##### Analysis of trajectories

We assess the stability of V2Rpp binding to $\beta$arrs in a manner reported previously.[1] The stability score is calculated as 7 Å – root mean square deviation (r.m.s.d.) of $C_\alpha$ atoms of V2Rpp, such that higher scores indicate stronger binding and lower scores reflect weaker binding. The 7 Å reference value is chosen to ensure all resulting scores are positive. The r.m.s.d. values, distances, and dihedral angles were calculated using the CPPTRAJ module of AMBER18,[2] after converting the trajectories to a compatible format using the MDConvert program from MDTraj.[3] Trajectories were visualized using VMD, and images were rendered with VMD and PyMOL. Plots were generated using custom Python scripts. Further, an array of analyses was performed as outlined below.

##### Inter-domain rotation angle

The inter-domain rotation angle serves as a measure of $\beta$arr activation.[1] The rotation is calculated by measuring the angle of the C-domain relative to the N-domain between the inactive (PDB ID 1G4M) and active (PDB ID 4JQI) $\beta$arr1 crystal structures, as outlined previously.[4] The same reference structures were used for both $\beta$arr1 and $\beta$arr2 to facilitate easier comparison. Naomi Latorraca generously provided the corresponding script.[4]

##### Inter-residue contacts and interaction type analysis

A contact between two residues is established if any two heavy atoms of the residues fall within a 4.5 Å distance for at least 75% of the simulation time. The contacts were determined using Trajcontacts program.[5] An in-house Python script was used to determine interaction types such as polar, charge-charge, van der Waals (vdW), hydrogen bonds (H-bond), and $\pi$-cation based on previously described geometric criteria.[6]

##### Analysis of structural motifs' movement

To analyze the conformation of the gate loop, we evaluated the projection of the $N_\zeta$ atom of K294 in $\beta$arr1 (K295 in $\beta$arr2) onto a stationary vector defined by the $C_\alpha$ atoms of R7 (R8 in $\beta$arr2) and R165 (R166 in $\beta$arr2) in the active-state $\beta$arr1 crystal structure (PDB ID 4JQI). The absolute value of the difference between the projection and the vector representing the initial position in 4JQI was negated, assigning a value of 0 to conformations close to the crystallographic active conformation, while increasingly negative values indicated deviations from the active state, as explained previously.[1] To assess the conformations of the finger loop, middle loop, C-loop, and back loop, we evaluated the projection of the $C_\alpha$ atoms of the central residues (L68, E134, F244, and I314 in $\beta$arr1; L69, E135, F245, and V315 in $\beta$arr2) onto vectors defined by the corresponding atoms in the inactive (PDB ID 1G4M) and active crystallographic positions.

##### Mutual information (MI)

Mutual Information (MI) is often used to quantify the correlation between two processes described by time series **X** and **Y**. It is defined as:

$$I_{\mathbf{XY}} = H(\mathbf{X}) + H(\mathbf{Y}) - H(\mathbf{X},\mathbf{Y}), \tag{1}$$

where **X** and **Y** represent two different time series. Here, $H(\mathbf{X})$ and $H(\mathbf{X},\mathbf{Y})$ are the Shannon entropies, given by:

$$H(\mathbf{X}) = -\int p(\mathbf{X}) \ln p(\mathbf{X})\, d\mathbf{X}, \tag{2}$$

$$H(\mathbf{X},\mathbf{Y}) = -\iint p(\mathbf{X},\mathbf{Y}) \ln p(\mathbf{X},\mathbf{Y})\, d\mathbf{X}\, d\mathbf{Y}. \tag{3}$$

Here, $p(\mathbf{X})$ is the marginal probability distribution of **X**, and $p(\mathbf{X},\mathbf{Y})$ is the joint probability distribution of **X** and **Y**. The correlation coefficient $r$ can be estimated from MI as:

$$r = \sqrt{1 - e^{-2I_{\mathrm{XY}}}}. \tag{4}$$

Time series of rotational angles and structural motif movements are considered as **X** and **Y** here. MI was computed using the adaptive partitioning method,[7] as implemented in the Python module `minfo`.[8]

##### $k$-means clustering algorithm

The $k$-means clustering algorithm partitions a given set of data points into $k$ clusters, where $k$ is a predefined number.[9] It works by iteratively assigning each data point to the closest centroid, which is the mean of the data points in the cluster. After all data points are assigned to the closest centroid, the algorithm updates the centroids by re-estimating the mean of each cluster. This process continues until the centroids no longer change or a maximum number of iterations is reached. We used $k$-means implemented in the SciPy Python library for our analysis.

##### Kernel principal component analysis (kernel PCA)

Kernel principal component analysis (kernel PCA) is a nonlinear dimensionality reduction technique that extends traditional principal component analysis (PCA) by incorporating kernel methods. This extension allows kernel PCA to capture complex nonlinear patterns within the data.[10,11] In kernel methods, the data is mapped to a higher-dimensional feature space through a nonlinear transformation, denoted by $\phi$.

$$\phi : \mathbb{R}^p \rightarrow \mathscr{F} \tag{5}$$

Here, $\mathscr{F}$ represents the feature space, which has a higher dimension than the original space, $\mathbb{R}^p$, where $x \in \mathbb{R}^p$ denotes the original input data. The mapping $\phi(x)$ refers to the transformation of the input data into this higher-dimensional feature space.

Typically, the feature space may have very high or even infinite dimensionality, making it challenging to explicitly construct the mapping function $\phi$ and calculate the transformed data. Fortunately, linear methods like PCA can be adapted to compute the inner product directly.[12] This allows the inner product in the feature space to be derived as a function of the original input data. Consider the following example of a mapping:

$$\phi : (x_1, x_2)^{'} \rightarrow (x_1^2, \sqrt{2}x_1x_2, x_2^2)^{'} \tag{6}$$

In this case, the inner product in the feature space can be rewritten as an algebraic expression in the original space:

$$\begin{aligned}
\phi(x)^{'}\phi(y) &= (x_1^2, \sqrt{2}x_1x_2, x_2^2) \cdot (y_1^2, \sqrt{2}y_1y_2, y_2^2)^{'} \\
&= ((x_1, x_2)(y_1, y_2)^{'})^2 \\
&= (x^{'}y)^2 \\
&= k(x, y)
\end{aligned} \tag{7}$$

Here, $k$ is the kernel function. The inner product of the mapped data is replaced by the kernel function in the original space. This means the inner product in the feature space can be calculated in the original space without explicitly applying the nonlinear mapping $\phi$. This technique is known as the *kernel trick*.[13]

Several kernel functions can be employed in kernel PCA, and we utilized the following. The linear kernel is given by $k(x, y) = x^{'}y$. The polynomial kernel is defined as $k(x, y) = (x^{'}y + c)^d$, where $c$ and $d$ are hyperparameters that influence the shape of the kernel. Lastly, the radial basis function (RBF) kernel is expressed as $k(x, y) = \exp\left(-\frac{\|x-y\|^2}{2\sigma^2}\right)$, where $\sigma$ denotes the width of the kernel.

After transforming the data into a higher-dimensional space using the aforementioned kernel functions, linear PCA is subsequently applied to the transformed feature space. The efficacy of the dimensionality reduction and the quality of the resulting low-dimensional representations were evaluated through clustering analysis using $k$-means and silhouette scores. All kernel PCA computations were implemented using the Scikit-learn library, ensuring efficient execution and consistency in the results.

##### t-distributed stochastic neighbor embedding (t-SNE)

t-SNE is an unsupervised nonlinear dimensionality reduction method,[14] effectively applied to analyze high-dimensional MD simulation trajectories.[15] It focuses on preserving the local neighborhood structure, ensuring that points close to each other in the original high-dimensional space remain close in the lower-dimensional embedding. In the original space, the probability $p(j|i)$ represents the likelihood of point $x_j$ being a neighbor of point $x_i$ rather than any other point $x_k$. This probability is modeled using a Gaussian distribution centered at $x_i$ with a standard deviation of $\sigma_i$ as,

$$p_{j|i} = \frac{\exp\left(-\|x_i - x_j\|^2/2\sigma_i^2\right)}{\sum_{k\neq i}\exp\left(-\|x_i - x_k\|^2/2\sigma_i^2\right)} \tag{8}$$

Similarly, the conditional probability in the embedded space, $q(j|i)$, with the same $n$ points initialized randomly, is computed but now based on a $t$-distribution. Having a longer tail than the Gaussian distribution, the $t$-distribution moves dissimilar points farther apart to ensure less crowding in the reduced space. Probability is in the lower dimensional space is represented as,

$$q_{i|j} = \frac{(1 + \|y_i - y_j\|^2)^{-1}}{\sum_{k\neq i}(\|y_i - y_k\|^2)^{-1}} \tag{9}$$

where $y_i$ and $y_j$ represent the low-dimensional embeddings. The aim of t-SNE is to minimize the dissimilarity between the high- and low-dimensional probability distributions $P_{ij}$ and $Q_{ij}$. This is achieved by minimizing the Kullback-Leibler (KL) divergence, defined as:

$$\mathrm{KL}(P_i \parallel Q_j) = \sum_i \sum_j p_{i|j} \log \frac{p_{i|j}}{q_{i|j}} \tag{10}$$

where $P_i$ and $Q_i$ are the joint probability distributions in the high- and low-dimensional space over all of the data points.

The primary hyperparameters that can be tuned in t-SNE are perplexity, learning rate, and the number of iterations. The perplexity parameter, $P$, determines the width of the Gaussian kernel, $\sigma_i$, in Eq 8. It is defined such that $\log_2 P = H(P_i) = -\sum_j p(j|i)\log_2 p(j|i)$, where $H(P_i)$ is the Shannon entropy for point $i$. This parameter loosely governs how many nearest neighbors each point is influenced by, thereby striking a balance between capturing local and global patterns. Lower perplexity values emphasize finer local structures, while higher values focus on the broader global distribution. To find the optimal perplexity, we experimented with a range of values, starting with an initial guess of perp $= \sqrt{n}$, and selected the value that produced the highest silhouette score.[15] We used the default learning rate of 200, which balances stability and convergence, and to prevent random fragmentation of clusters, the number of iterations was set to 3500, as previously described.[15,16] The choice of the distance metric, here was the Euclidean distance.

For t-SNE calculations, we relied on the Scikit-learn implementation, leveraging the Barnes–Hut approximation to enhance computational efficiency for gradient calculations.

###### Uniform manifold approximation and projection (UMAP)

UMAP is a dimensionality reduction technique based on fuzzy topology, designed to preserve the local structure of data while projecting it into a lower-dimensional space.[17] It operates by constructing probability distributions that describe the relationships between high-dimensional data points, similar to t-SNE but with key differences in its formulation.

In UMAP, the high-dimensional probability distribution between points is defined as:

$$p_{ij} = \exp\left(-\frac{d(x_i, x_j)}{\sigma_i}\right) \tag{11}$$

where $p_{ij}$ represents the probability of connection between points $x_i$ and $x_j$, and $\sigma_i$ is a local scale parameter specific to each point. The distance function $d(x_i, x_j)$ measures the distance between two data points in the original high-dimensional space.

In contrast to t-SNE, UMAP uses a unique local distance metric for each pair of points, which allows for a more accurate representation of the local topology. The probability distribution in the low-dimensional space is given by:

$$q_{ij} = \frac{1}{1 + \left\|y_i - y_j\right\|^2} \tag{12}$$

where $y_i$ and $y_j$ are the low-dimensional coordinates of the data points $x_i$ and $x_j$, and the distance function is squared Euclidean distance. The parameter $b$ controls the spread of the low-dimensional probability distribution.

A key difference between UMAP and t-SNE lies in the loss function used during optimization. While t-SNE minimizes the Kullback-Leibler (KL) divergence between the high-dimensional and low-dimensional probability distributions, UMAP minimizes the cross-entropy (CE) loss function, which is defined as:

$$\mathrm{CE}(X,Y) = -\sum_{i,j}\left[p_{ij}\log q_{ij} + (1-p_{ij})\log(1-q_{ij})\right] \tag{13}$$

This cross-entropy loss function is advantageous as it better preserves the relative distances between points in both the high- and low-dimensional spaces, particularly for both small and large distances. UMAP's approach facilitates a more robust and flexible dimensionality reduction method compared to t-SNE.

The primary hyperparameters that can be tuned in UMAP are the number of neighbors `n_neighbors` and the minimum distance `mindist`. The `n_neighbors` parameter controls how many nearest neighbors each point considers when constructing the local neighborhood, balancing local versus global structure. Smaller values of `n_neighbors` emphasize local details, while larger values capture broader global patterns. The `mindist` parameter determines the minimum distance between points in the low-dimensional space, affecting the tightness of the clusters. Smaller values of `mindist` result in more compact clusters, while larger values spread the points out. We experimented with various values for both `n_neighbors` (ranging from 100 to 1000) and `mindist` (0.01, 0.02, 0.05, 0.1, 0.25, 0.5, 1) to assess their effects on the resulting embeddings. Additionally, we chose the distance metric to be the Manhattan distance. The optimal combination of these hyperparameters was selected based on maximizing the silhouette score.

UMAP projections were performed using the Python implementation available at `https://github.com/lmcinnes/umap`.

For both $\beta$arr1 and $\beta$arr2, UMAP performed best in terms of high silhouette scores for clustering in the reduced dimensional space compared to tSNE and kernel PCA, as shown in Figures S9 and S10.

**Machine learning classification and feature importance analysis**

We employed a two-phase approach for the classification task aimed at predicting the $\beta$arr conformational clusters of phospho-systems based on a combination of structural and energetic features. For this we gathered three distinct feature sets relevant to the protein structure and dynamics.The feature sets are as follows: (i) dihedral angles, including that of backbone and side-chain—$\phi$, $\psi$, and $\chi$s; (ii) inter-residue interaction energies, calculated as the sum of electrostatic and van der Waals energies between residues maintaining at least 10% contact throughout the simulation; and (iii) solvent-accessible surface area (SASA), which measures the exposure of each residue to the solvent environment. All features were normalized using standard normalization to ensure consistent scaling across features. Standard normalization transforms the data to have a mean of zero and a standard deviation of one, making the features dimensionless and comparable. The transformation is performed as:

$$z = \frac{x-\mu}{\sigma} \tag{14}$$

where $z$ is the normalized value, $x$ is the original feature value, $\mu$ is the mean of the feature, and $\sigma$ is the standard deviation of the feature.

In the initial phase, we focused on eliminating irrelevant or redundant features to retain only those that contributed meaningfully to the classification of the $\beta$arr conformations. The features were evaluated by applying several machine learning (ML) algorithms, including support vector machines (SVM), decision trees (DT), random forests (RF), extra trees (ET), extreme gradient boosting (XGB), and multilayer perceptron (MLP). The classification performance for each feature set was assessed independently, and accuracies exceeding 98% were achieved (Figure S11), indicating that the selected features were highly informative for the classification task.

To interpret the significance of individual features and understand their contribution to the model's decisions, we utilized local interpretable model-agnostic explanations (LIME). LIME is an explainable artificial intelligence (XAI) method designed to provide transparency into complex ML models by approximating them with simpler, interpretable models on a local level. LIME works by perturbing the input features and observing the effect of these changes on the model's predictions. For each perturbed data point, LIME trains a surrogate model (typically a simple linear/explainable model) to approximate the behavior of the complex model in that region of the data. The importance of each feature is then assessed by measuring how much its perturbation influences the output of the surrogate model. Here, to identify the most important features, we retained only the top 99% of features consistently identified as important by at least three out of six ML models using the LIME explainer.

In the second phase, we integrated the top-ranked features from each of the three feature sets (dihedral angles, inter-residue interaction energies, and SASA) into a refined feature set. This integrated set was then used to train three machine learning models: extra trees (ET), extreme gradient boosting (XGB), and multilayer perceptron (MLP). These classifiers were selected based on their superior performance in the first phase, as evidenced by their high accuracy and low log loss (Figure S11). When trained on the refined feature set, the classifiers achieved average accuracies of approximately 99 % (Figure S12), demonstrating the effectiveness of our feature selection strategy in identifying a minimal yet highly informative subset of features. We further refined the features by selecting only those contributing to the top 90% of the overall feature importance across all three models. This selection significantly reduced the dimensionality, while retaining the most important features. The corresponding residues were identified and used for downstream analyses.

###### Evaluation of machine learning models

We evaluated classifier performance using 5-fold cross-validation to ensure robustness across data splits. The data was preprocessed by separating features and labels, and class imbalance (over 30%) was addressed by randomly subsampling the majority class.

For assessing the model performance, we utilized log loss (cross-entropy loss) in addition to the commonly used accuracy metric, as the former is effective in evaluating imbalanced class distributions. Accuracy measures the proportion of correctly classified instances out of all predictions, calculated as,

$$\text{accuracy} = \frac{\text{Number of correct predictions}}{\text{Total number of predictions}} \tag{15}$$

Log loss is a metric that penalizes false classifications based on predicted probabilities for each class, estimated as,

$$\text{log loss} = -\frac{1}{N}\sum_{i=1}^{N}\sum_{j=1}^{M} y_{ij}\log(p_{ij}), \tag{16}$$

where $N$ is the number of instances, $M$ is the number of classes, $y_{ij}$ is the indicator function (1 if the instance $i$ belongs to class $j$, 0 otherwise), and $p_{ij}$ is the predicted probability of instance $i$ belonging to class $j$.

###### Marking the effector-binding regions

Effector-binding regions on $\beta$arr1 and $\beta$arr2 were annotated using residue-level data from prior structural and spectroscopic studies compiled previously.[18,19] Only studies identifying specific binding residues were included; those reporting broad residue ranges were excluded to ensure accuracy.

###### Graph neural network-based autoencoder

We utilized a graph neural network (GNN)-based autoencoder to construct allosteric dynamic residue networks (DRNs). A key challenge in creating DRNs is accurately assigning meaningful weights to the edges, as these weights represent the strength of the connection between residues. Standard methods that rely on contacts, inter-residue interaction energies, or correlations may fail to capture important physical properties related to allosteric effects when used individually. To address this, we introduce a deep learning approach that leverages the capabilities of GNNs to learn edge weights from input features such as atomic positions (and thereby, correlations in fluctuations), interaction energies, and SASA. Such an approach also aggregates information from neighboring nodes and edges through message passing, thereby learning relationships between residues, which is highly beneficial in identifying allosteric communications.

**Graph representation**: Before passing though GNN autoencoder, each MD snapshot was converted to an unweighted graph, with residues as nodes and edges defined by inter-residue contacts that persisted for at least 10% of the total simulation time. A graph is represented as $G = (V, E)$, where $V$ denotes the set of $n$ nodes, corresponding to the $n$ residues, and $E$ is the set of edges represented as $\{v_1, v_2\}$ where $v_1, v_2 \in V$. Node features are given as $\mathbf{X} \in \mathbb{R}^{n\times d}$, where $d$ is the number of input features per node. The node features are defined as follows. (i) The position of the $C_\alpha$ atom relative to the origin after aligning the trajectories to the starting conformation. This alignment removes rotations and translations. (ii) The sum of interaction energy weights for a residue with all its interacting residues. Each weight is derived from the total inter-residue interaction energy, which is the sum of electrostatic and van der Waals interactions. These weights are normalized by dividing by the most favorable (i.e., most negative) energy value. Any negative values resulting from this normalization were set to zero to eliminate unfavorable interactions. (iii) The SASA of each residue. To ensure consistent scaling across all features, standard normalization was applied. Each feature was transformed to have a mean of zero and a standard deviation of one.

**GNN autoencoder architecture**: The GNN-based autoencoder consists of two main parts: the encoder and the decoder. Each part involves two key message-passing layers: one that aggregates information from nodes to other nodes (node-to-node) and another that aggregates from nodes to edges (node-to-edge). In the encoder, these layers are stacked to capture complex interdependencies between nodes and edges, while in the decoder, the reverse operations are performed to reconstruct the original input features.

The encoder maps node features $\mathbf{X}$ to latent edge features $\mathbf{Z} \in \mathbb{R}^{m \times l}$, where $l$ is the latent dimensionality. It consists of two main stages: node-to-node message passing and node-to-edge aggregation.

In the node-to-node message passing step, node features are updated by aggregating information from their neighbors:

$$\mathbf{h}_i^{\text{new}} = \mathbf{W}\mathbf{h}_i + \sum_{j \in \mathcal{N}(i)} \frac{1}{\sqrt{\deg(i) \cdot \deg(j)}} \mathbf{W}\mathbf{h}_j, \tag{17}$$

where $\mathbf{h}_i$ is the feature vector of node $i$, $\mathcal{N}(i)$ denotes the neighbors of node $i$, $\deg(i)$ is the degree of node $i$, and $\mathbf{W} \in \mathbb{R}^{d \times d'}$ is a trainable weight matrix. Self-loops are added to ensure each node's information is included in the update.

In the node-to-edge aggregation step, updated node features are transformed into edge features:

$$\mathbf{z}_{ij} = \mathbf{W}_1\mathbf{h}_i + \mathbf{b}, \tag{18}$$

where $\mathbf{z}_{ij}$ is the latent feature vector of edge $(i, j)$, $\mathbf{W}_1 \in \mathbb{R}^{d' \times l}$ is a trainable weight matrix, and $\mathbf{b} \in \mathbb{R}^l$ is a bias term. The output of the encoder is $\mathbf{Z}$, the latent edge features. The latent space is defined by the edge features $\mathbf{Z}$ of dimension $m \times l$, where $l$ is typically much smaller than $d$. $l$ is set to 1 to ensure that each edge is represented by a scalar value, which effectively serves as a weight for the corresponding edge.

The decoder reconstructs the original node features $\mathbf{X}$ from the latent edge features $\mathbf{Z}$ through two steps: edge-to-node aggregation and node reconstruction.

In the edge-to-node aggregation step, the decoder aggregates information from edges back to nodes:

$$\mathbf{h}_i^{\text{agg}} = \sum_{j \in \mathcal{E}(i)} \mathbf{W}_2\mathbf{z}_{ij} + \mathbf{b}_2, \tag{19}$$

where $\mathbf{h}_i^{\text{agg}}$ is the aggregated feature vector for node $i$, $\mathbf{W}_2 \in \mathbb{R}^{l \times d''}$ is a trainable weight matrix, and $\mathbf{b}_2 \in \mathbb{R}^{d''}$ is a bias term.

In the node reconstruction step, the aggregated node features are mapped back to the input space:

$$\mathbf{h}_i^{\text{recon}} = \mathbf{W}_3\mathbf{h}_i^{\text{agg}} + \mathbf{b}_3, \tag{20}$$

where $\mathbf{h}_i^{\text{recon}}$ is the reconstructed feature vector for node $i$, $\mathbf{W}_3 \in \mathbb{R}^{d'' \times d}$ is a trainable weight matrix, and $\mathbf{b}_3 \in \mathbb{R}^d$ is a bias term.

**Loss function**: The reconstruction quality is measured using the Mean Squared Error (MSE) loss:

$$\mathcal{L}_{\text{recon}} = \frac{1}{n}\sum_{i=1}^{n} \|\mathbf{x}_i - \mathbf{h}_i^{\text{recon}}\|_2^2, \tag{21}$$

where $\mathbf{x}_i$ is the original feature vector of node $i$, and $\mathbf{h}_i^{\text{recon}}$ is the reconstructed feature vector of node $i$.

**Training**: The model is trained using the Adam optimizer with a learning rate of $\eta = 0.001$. The encoder and decoder are trained jointly to minimize $\mathcal{L}_{\text{recon}}$. Training is performed for 500 epochs, with early stopping based on validation performance.

**Implementation edge weight optimization**: The input node features were calculated using CPPTRAJ and MDTraj. PyTorch[20] with the Torch Geometric module[21] implemented in Python was used to build the GNN autoencoder. After training, the latent space representing the learned edge weights for each snapshot was averaged for each system to construct the respective DRNs.

##### Dynamic residue networks and suboptimal paths

Dynamic residue networks constructed from MD simulation data are extensively used to study the allosteric communications between distant residues in proteins and nucleic acids.[5,22–24] Networks are constructed by considering $C_\alpha$ atoms of the $\beta$arrs and V2Rpp as nodes. The edges and weights were obtained from the GNN autoencoder, and weights ($w$) were converted to lengths as,

$$d_{XY} = -\log(w_{XY}) \tag{22}$$

The length of a path between two distant nodes is defined as the sum of edge weights along that path:

$$l = \sum_{XY} d_{XY} \tag{23}$$

where $d_{XY}$ is the edge weight of all node pairs in the path.[22,25] An optimal or shortest path $l_0$ between two nodes is the one with the least length and corresponds to the maximum correlation. Furthermore, suboptimal paths (SOP) between any two nodes are the additional paths connecting them, and have lengths:

$$l_{\text{SOP}} \leq l_0 + l_{\text{cut}} \tag{24}$$

where $l_{\text{cut}}$ is an arbitrary distance cutoff. Herein, testing $l_{\text{cut}} = 20$ and 30 yielded similar results (Figure S17). The results presented in the main text and SI are based on $l_{\text{cut}} = 20$. SOPs were calculated using the `subopt` program implemented in the `NetworkView` plugin in VMD.[22,25,26]

.

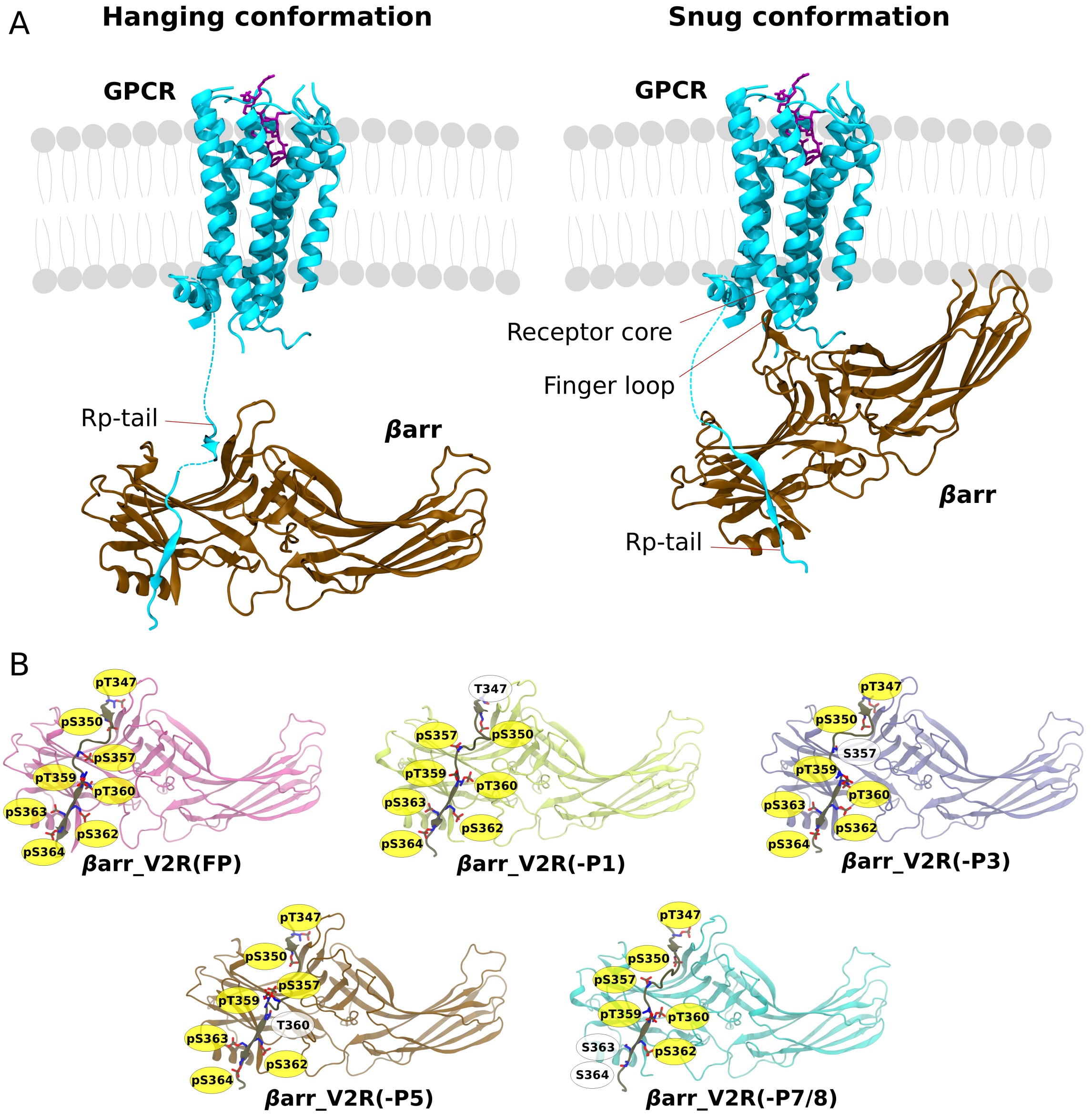


Figure S1: (A) Different modes of $\beta$arr interaction with GPCRs: Hanging and snug conformations are shown as cartoons. (B) Cartoon representation of simulated $\beta$arr1/$\beta$arr2 structures bound with V2Rpp, showing distinct phospho-patterns. Phosphorylated serines/threonines are shown in yellow, and unphosphorylated in white.

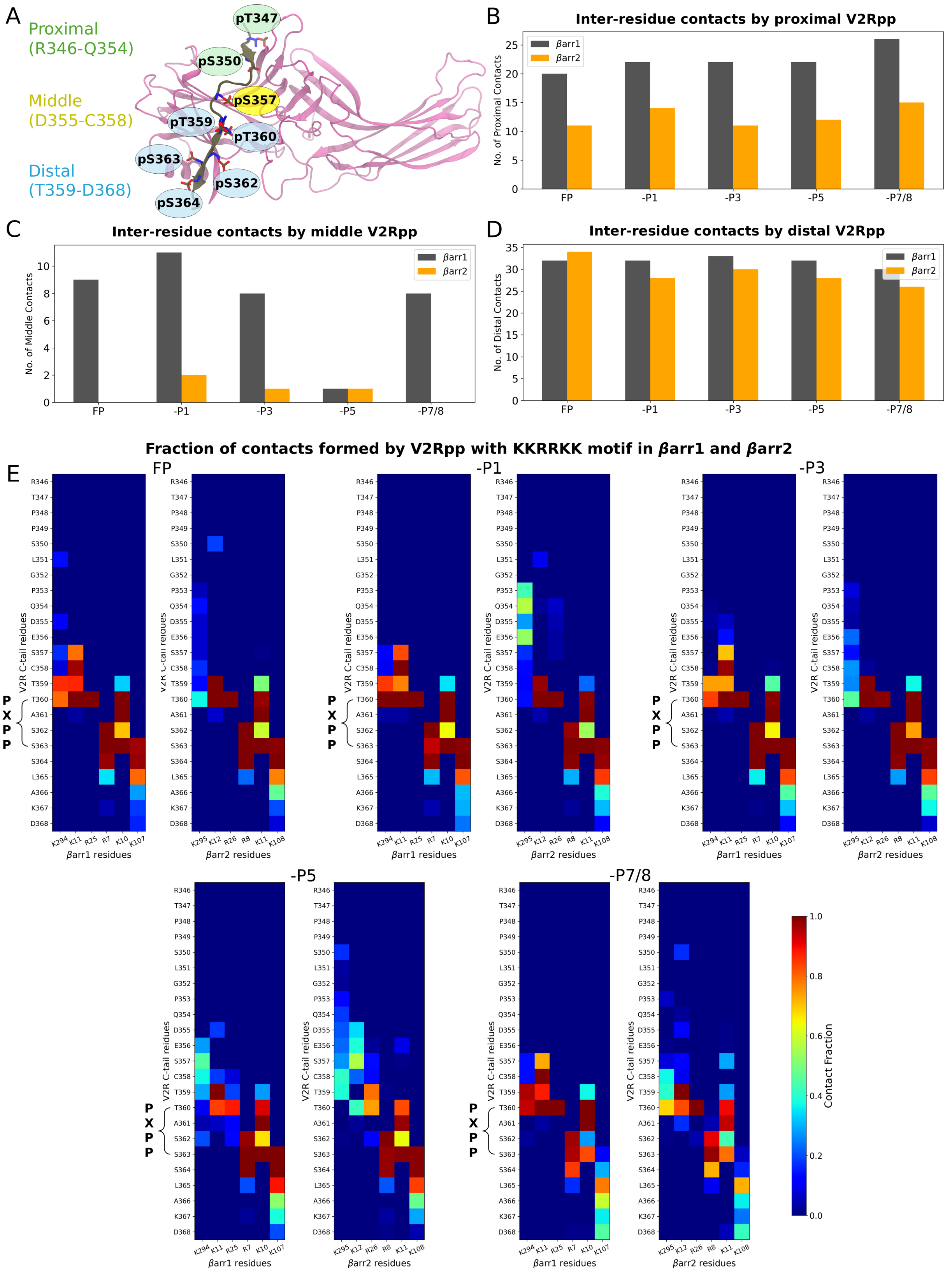


Figure S2: **(A)** $\beta$arr–V2Rpp bound structure showing different parts of V2Rpp: proximal, middle, and distal. Number of inter-residue contacts formed by **(B)** the proximal part of V2Rpp, **(C)** the middle part of V2Rpp, and **(D)** the distal part of V2Rpp for each system of $\beta$arr1 and $\beta$arr2. **(E)** Heatmap showing the fraction of contacts formed by the PxPP motif of V2Rpp with the KKRRKK motif in $\beta$arr1 and $\beta$arr2 for each system.

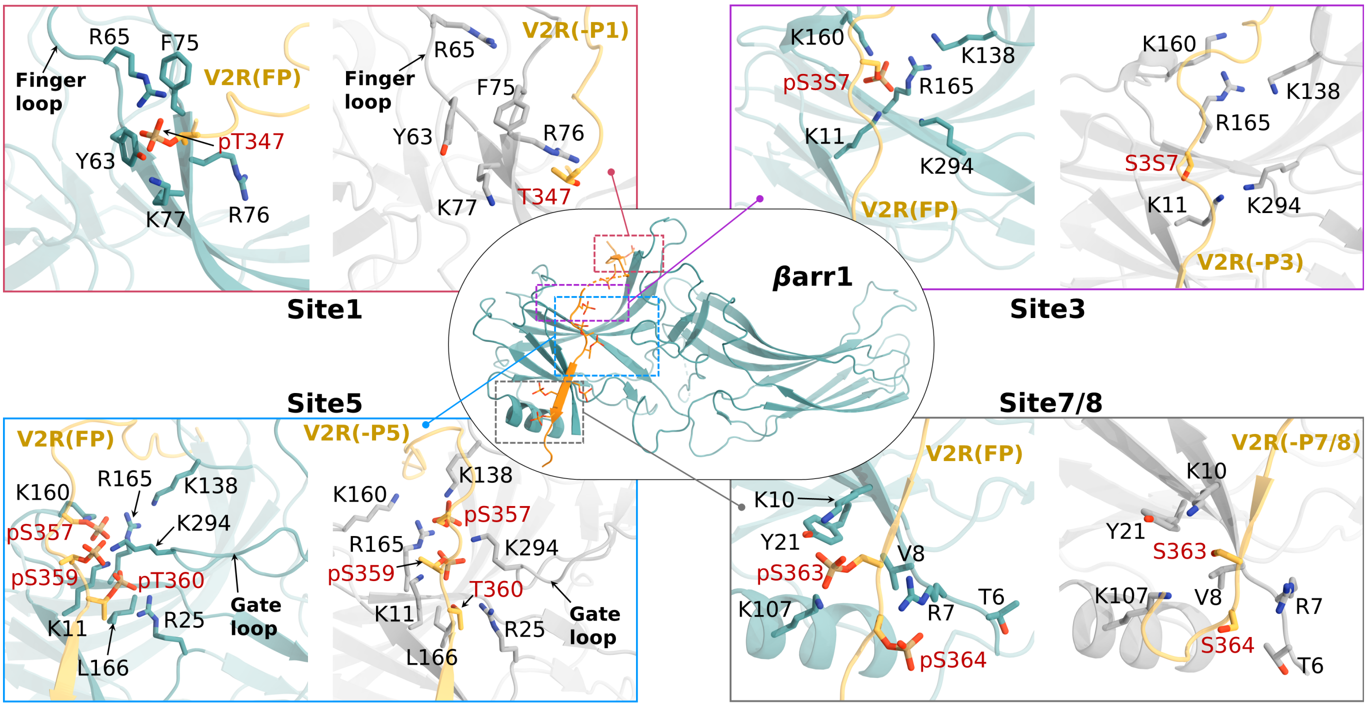


Figure S3: Different phospho-sites on the fully phosphorylated V2Rpp-bound $\beta$arr1 are depicted at the center. Zoomed-in views of each site are shown in the surrounding boxes. Each box highlights the residues that interact with the peptide in the FP system (shown in teal) and the loss of those interactions in specific phospho-systems due to the absence of phosphorylation (shown in white).

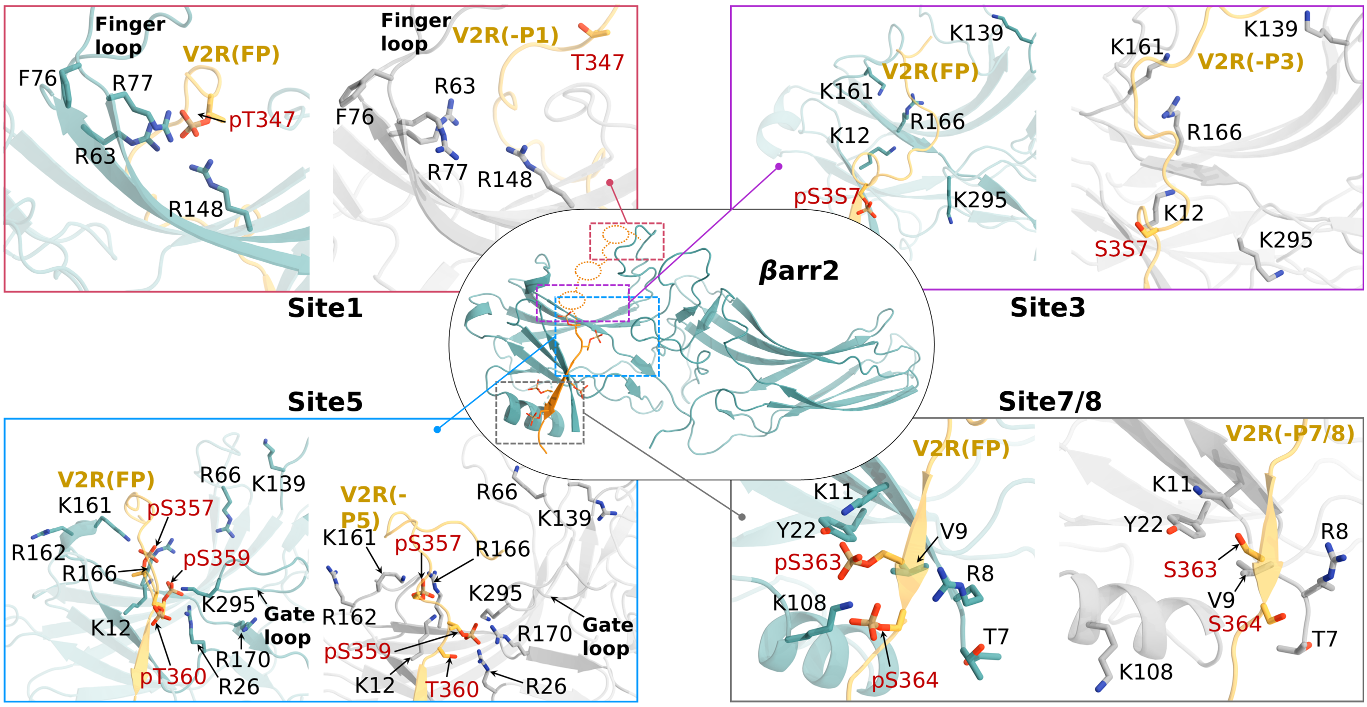


Figure S4: Different phospho-sites on the fully phosphorylated V2Rpp-bound $\beta$arr2 are depicted at the center. Zoomed-in views of each site are shown in the surrounding boxes. Each box highlights the residues that interact with the peptide in the FP system (shown in teal) and the loss of those interactions in specific phospho-systems due to the absence of phosphorylation (shown in white).

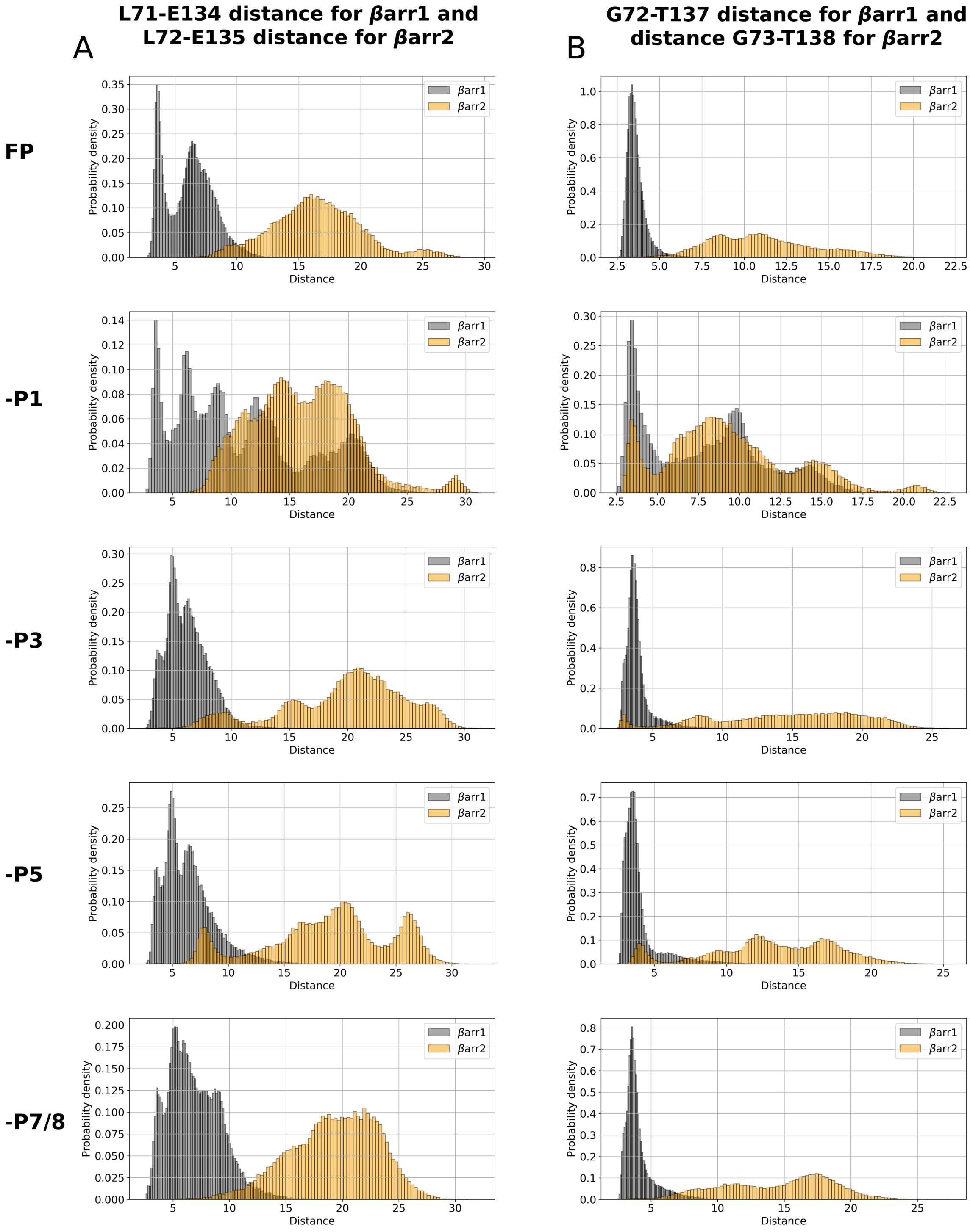


Figure S5: The distributions of interaction distances between the finger loop and the middle loop. Two specific pairs **(A)** L71-E134 and **(B)** G72-T137 are shown for $\beta$arr1, along with the corresponding pairs in $\beta$arr2, for each phospho-system.

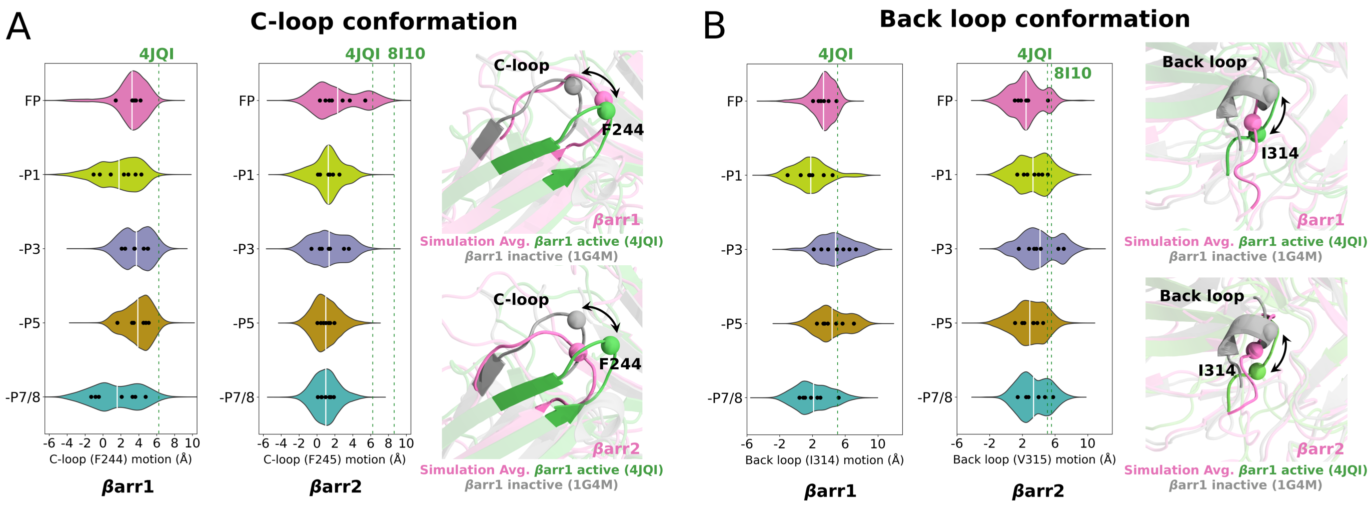


Figure S6: Conformational changes in key structural elements of $\beta$arr1 and $\beta$arr2 induced by unique phospho-patterns on V2Rpp: motion of **(A)** the C-loop and **(B)** the back loop. Violin plots depict the distributions, showing mean values (white lines) and individual trajectory averages (black dots). Cartoon representations of the motions are shown on the right. The matrices used to measure conformational dynamics are described in the SI Methods section.

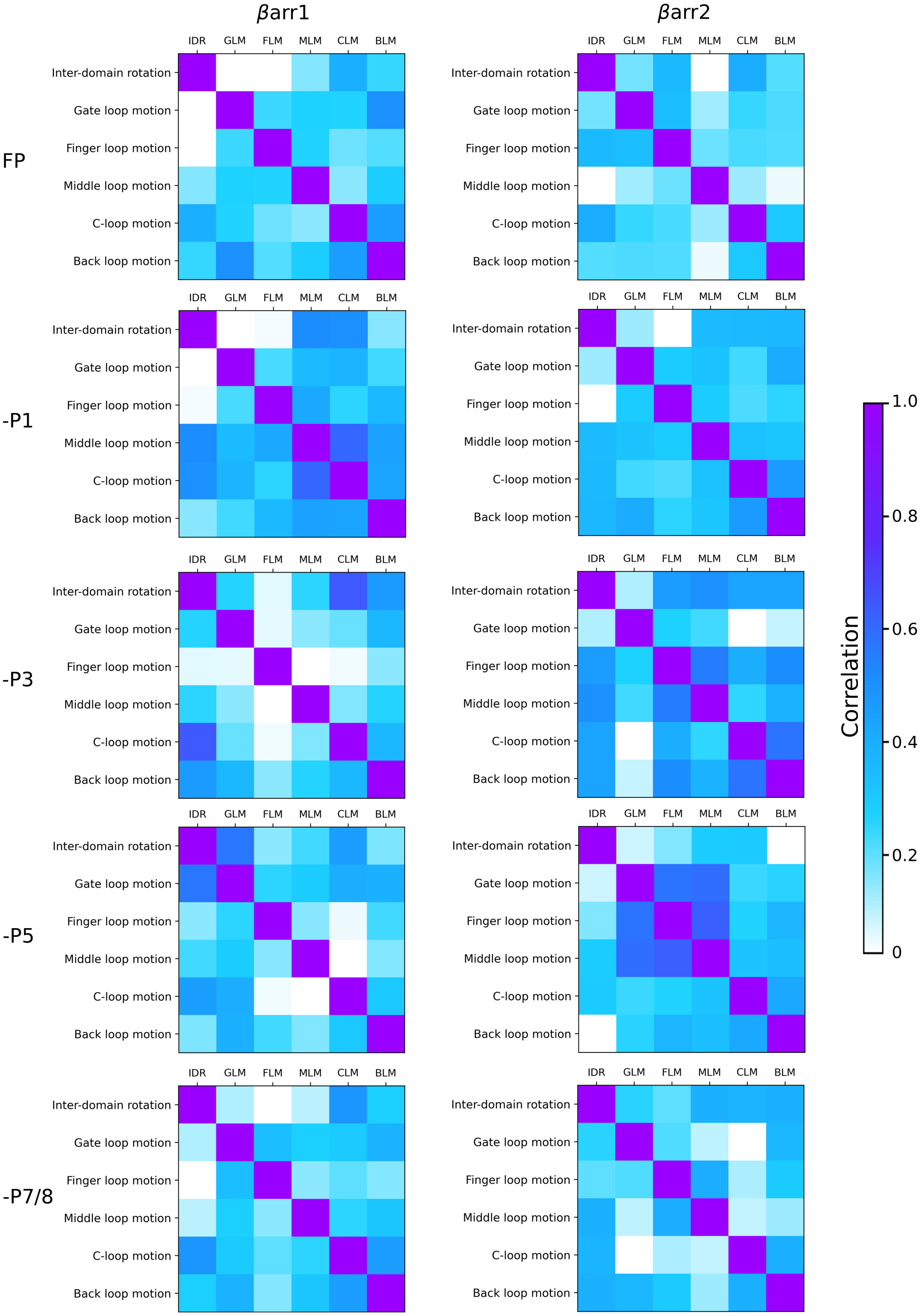


Figure S7: The correlation between structural movements in $\beta$arr1 and $\beta$arr2 for each phospho-system. The metrics used to measure these motions are described in the SI Methods section. All except a few cases show values of non-diagonal entries $< 0.6$, implying very little correlation between the motions in general.

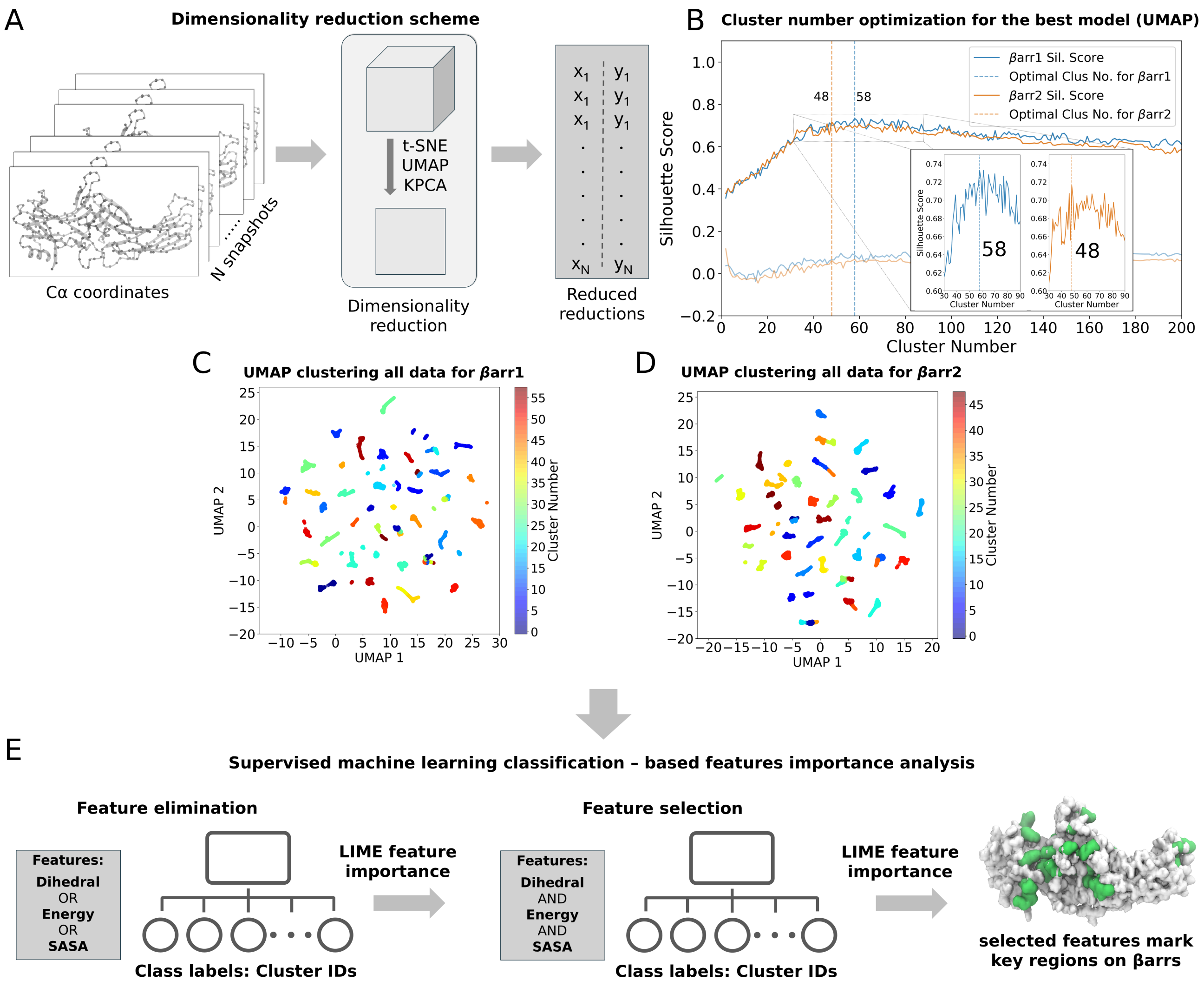


Figure S8: **(A)** The dimensionality reduction scheme involves using $C_\alpha$ coordinates of $\beta$arrs as features, which are subjected to different dimensionality reduction algorithms while varying the hyperparameters of each algorithm (the UMAP resulted in high-quality clusters, as shown in Figures S9 and S10). **(B)** Clustering of the reduced UMAP dimensions is performed, and the optimal number of clusters is selected based on silhouette scores. The product of silhouette scores in the reduced dimension and in the higher dimensions (where higher-dimension clusters are a one-to-one mapping of the lower-dimension clusters) is also reported as a faded line. A value greater than 0 ensures good cluster separation in the higher dimensions as well. Results of clustering UMAP dimensions for **(C)** $\beta$arr1 and **(D)** $\beta$arr2. Clusters are distinctly colored. **(E)** A scheme of supervised learning, where the input features include dihedral angles, inter-residue interaction energies of residues in contact, or SASA of every residue, each considered separately in the first phase of feature elimination. After removing the least important 1% of features, determined using LIME, the remaining features are combined for the second phase of feature selection. Finally, residues corresponding to the top 90% of important features are highlighted on the structure.

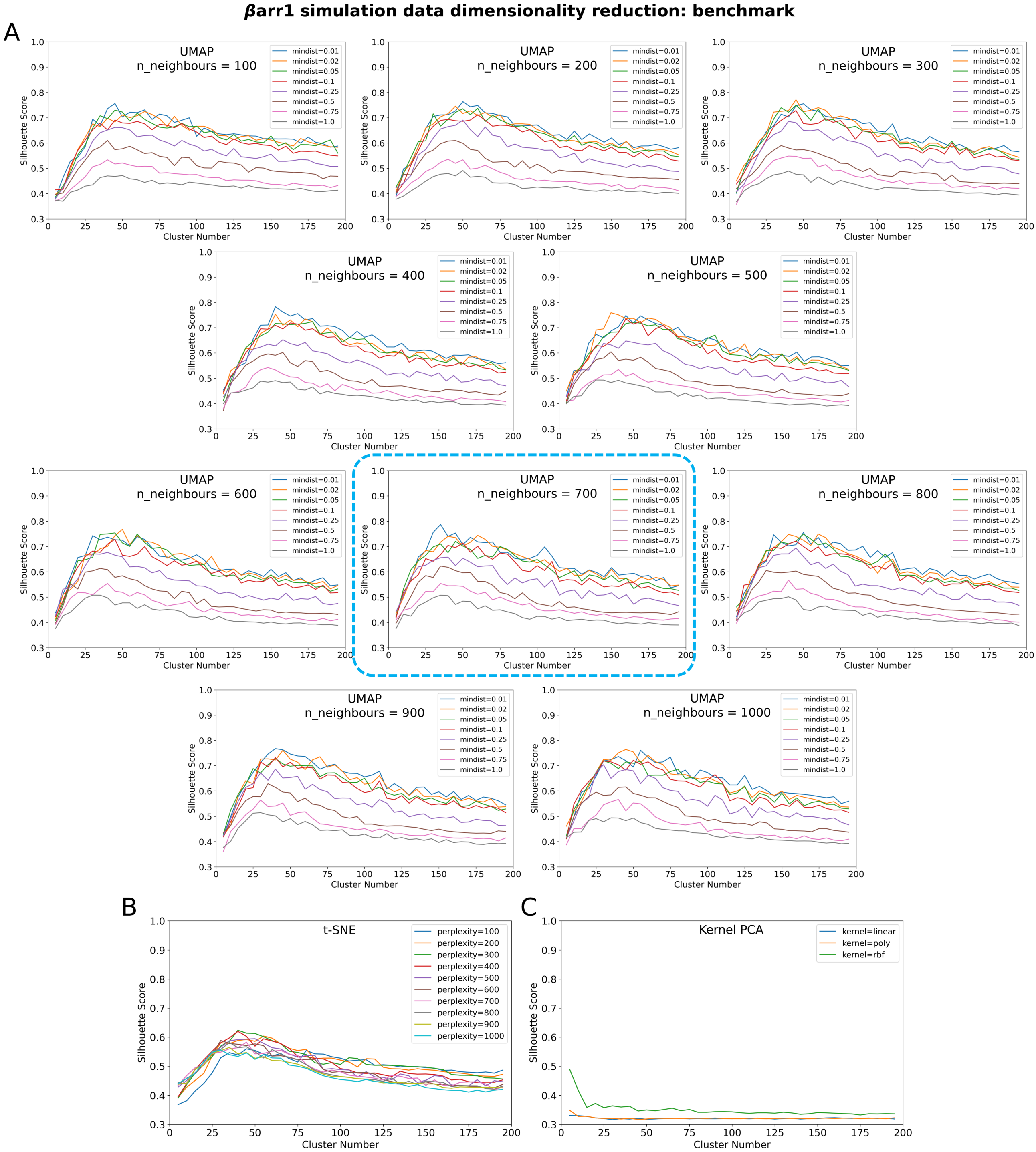


Figure S9: Hyperparameter tuning for dimensionality reduction of $C_\alpha$ coordinates of $\beta$arr1 systems. **(A)** UMAP: the `n_neighbors` and `mindist` parameters are varied, and silhouette scores are reported for each combination after clustering. **(B)** t-SNE: different perplexity values are tested, and the resulting silhouette scores are reported. **(C)** kernel PCA: various kernels are evaluated, and their performance is assessed using silhouette scores.

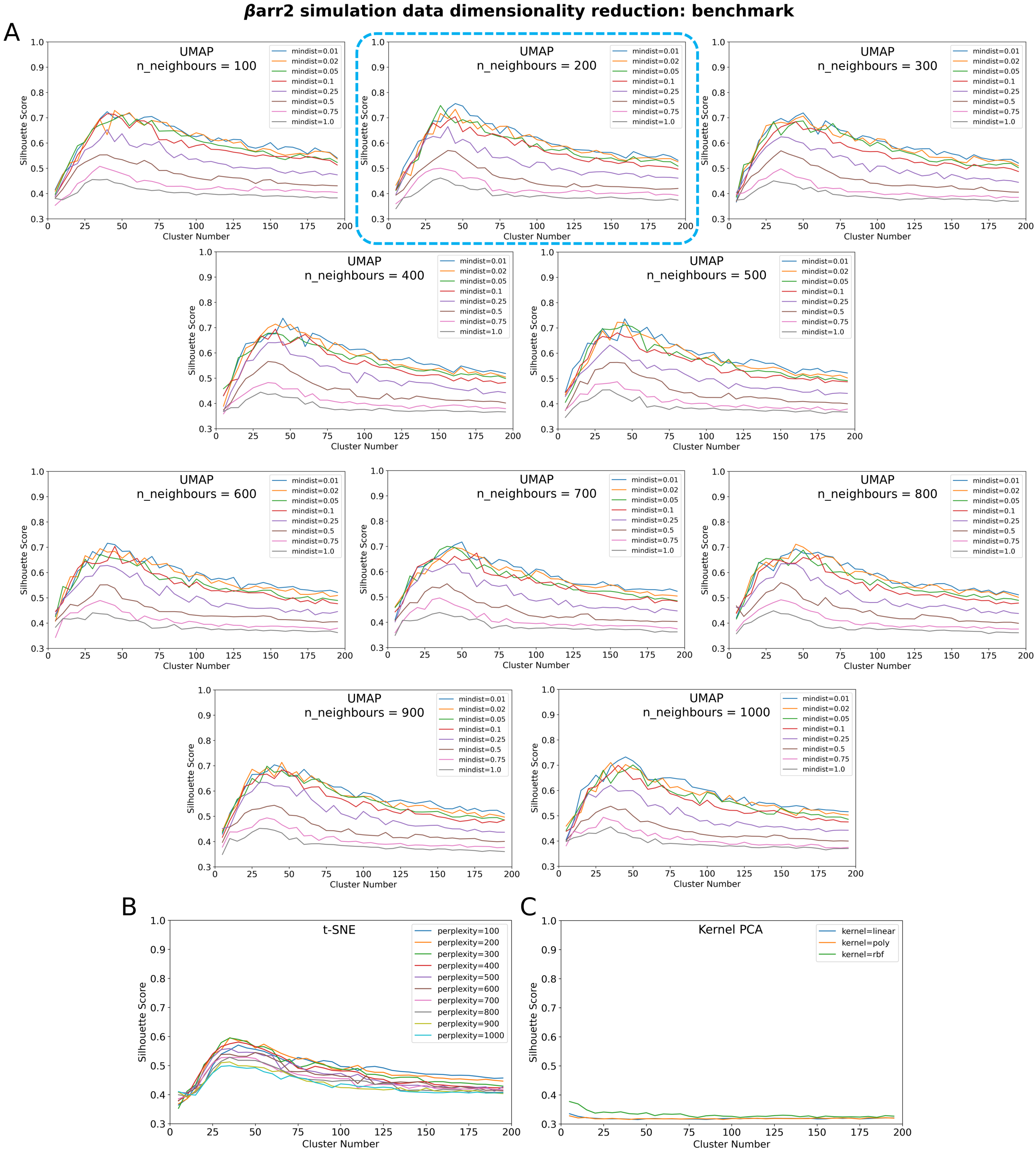


Figure S10: Hyperparameter tuning for dimensionality reduction of $C_\alpha$ coordinates of $\beta$arr2 systems. **(A)** UMAP: the `n_neighbors` and `mindist` parameters are varied, and silhouette scores are reported for each combination after clustering. **(B)** t-SNE: different perplexity values are tested, and the resulting silhouette scores are reported. **(C)** kernel PCA: various kernels are evaluated, and their performance is assessed using silhouette scores.

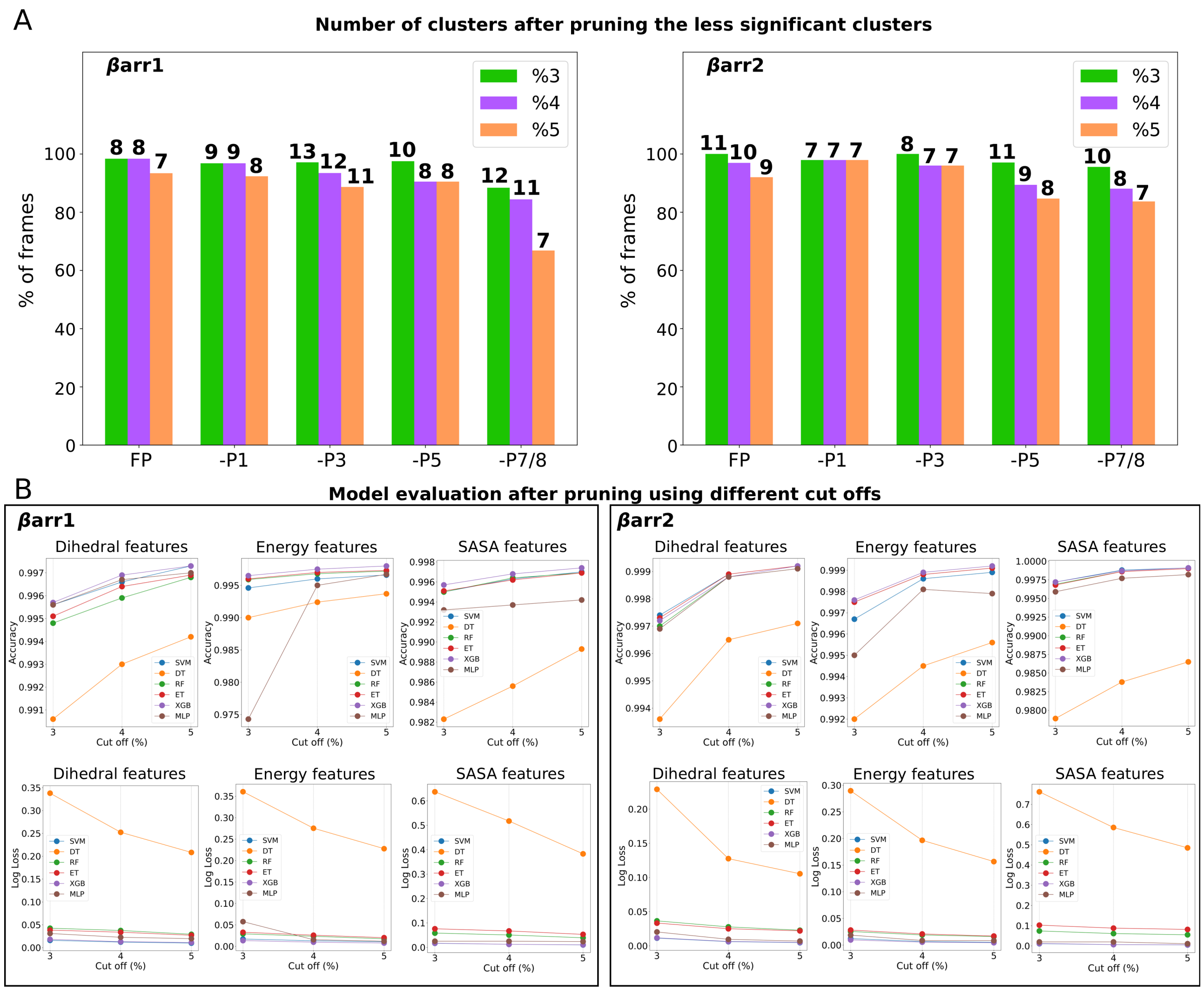


Figure S11: **(A)** The number of clusters obtained for each phospho-system after pruning insignificant ones. Pruning is performed using three different cutoffs, where each cutoff specifies the minimum percentage of snapshots that must be present in all selected clusters for $\beta$arr1 (left) and $\beta$arr2 (right). The final number of selected clusters is indicated above each bar in bold. **(B)** Accuracy and log loss scores for models built using various ML classification algorithms, evaluated on all pruned datasets for $\beta$arr1 (left) and $\beta$arr2 (right) (phase 1 of ML classification).

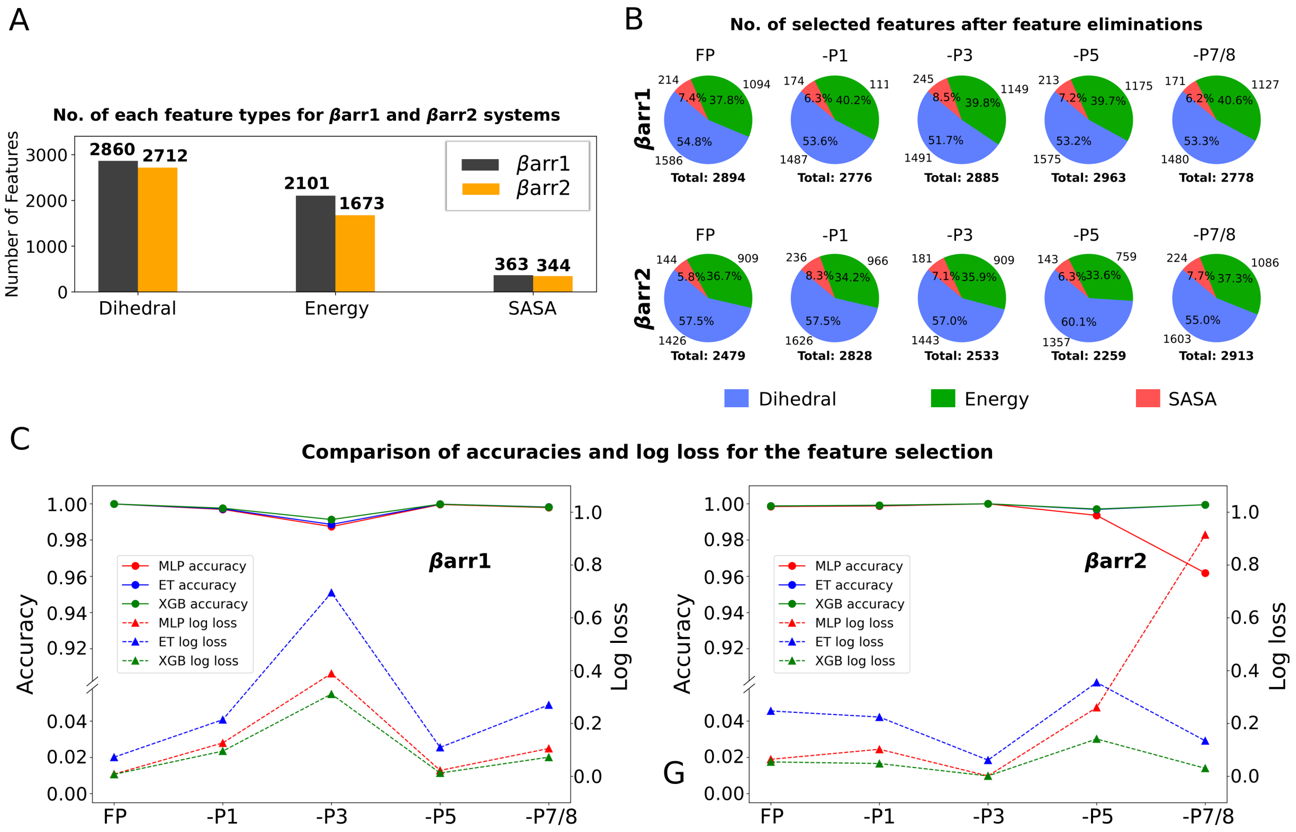


Figure S12: **(A)** Number of features in the dihedral, energy, and SASA feature sets considered during the initial phase of ML classification. **(B)** Number of features retained after feature elimination in the first phase. **(C)** Classification performance in the second phase (feature selection), showing accuracy and log loss scores using the combined feature set derived from phase one.

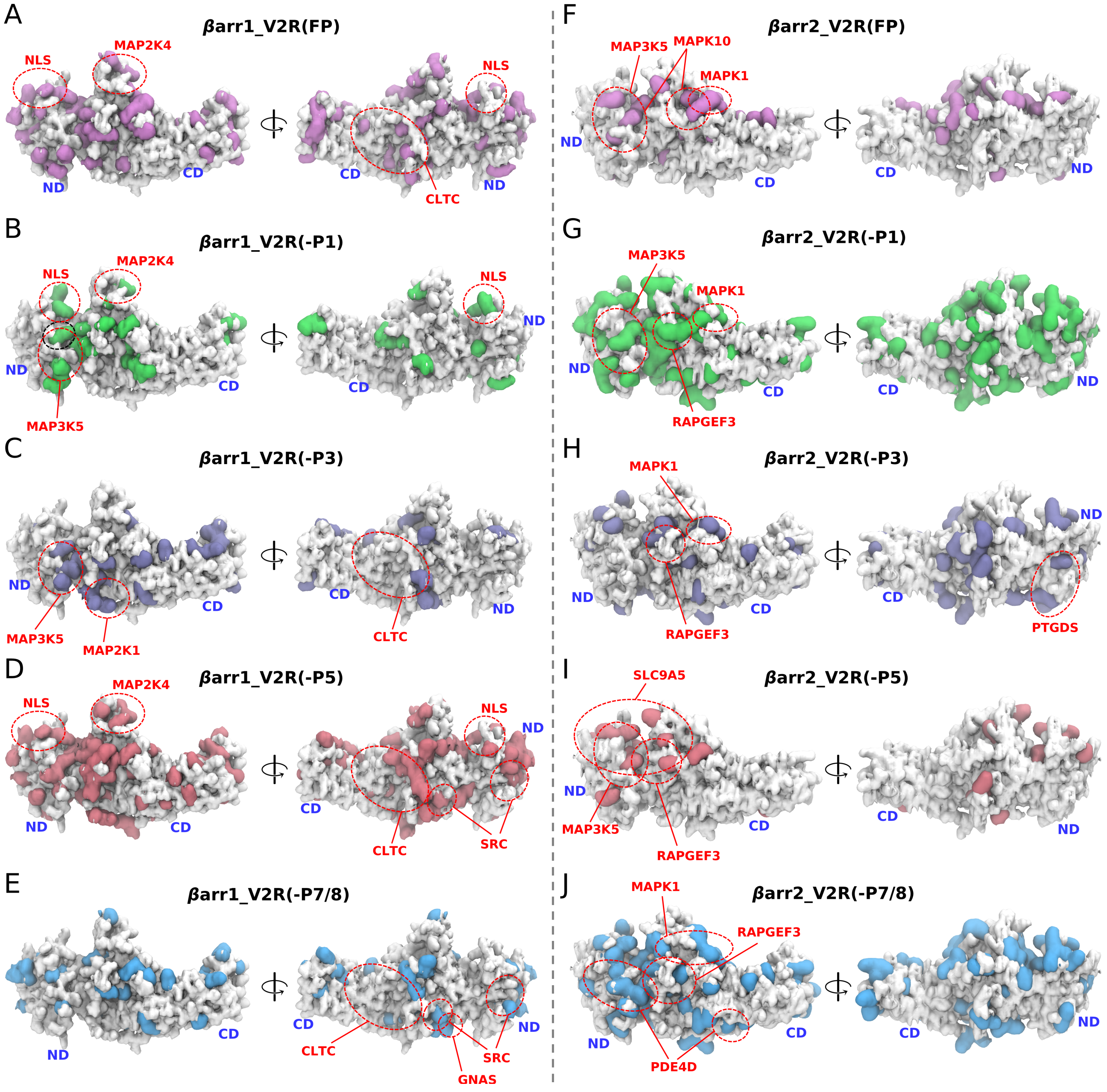


Figure S13: Important residues obtained through ML classifications are marked on the $\beta$arr structures, highlighting the most significant effector binding regions (top 3) for each phospho-system, identified using the percentage of residues for **(A-E)** $\beta$arr1 and **(F-J)** $\beta$arr2.

| Sl No. | $\beta$arr1_V2R(FP) | $\beta$arr1_V2R(-P1) | $\beta$arr1_V2R(-P3) | $\beta$arr1_V2R(-P5) | $\beta$arr1_V2R(-P7/8) |
|---|---|---|---|---|---|
| 1 | 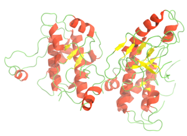 MAP2K4 (60.0%) | Nuclear localization signal<br>NLS (28.6%) | 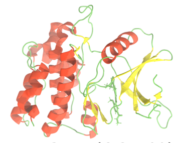 MAP3K5 (26.7%) | 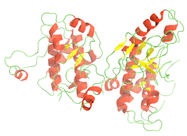 MAP2K4 (60.0%) | 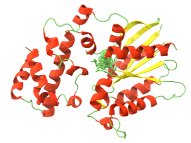 GNAS (100.0%) |
| 2 | Nuclear localization signal<br>NLS (42.9%) | 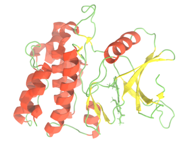 MAP3K5 (20.0%) | 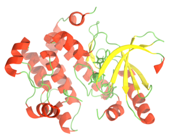 MAP2K1 (22.2%) | Nuclear localization signal<br>NLS (42.9%) | 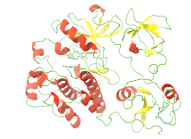 SRC (25.0%) |
| 3 | 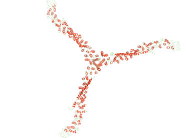 CLTC (36.8%) | 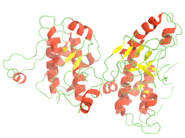 MAP2K4 (20.0%) | 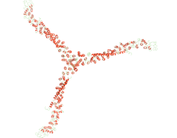 CLTC (15.8%) | 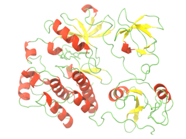 SRC (33.3%) | 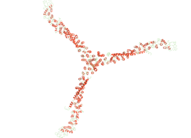 CLTC (21.1%) |
| 4 | 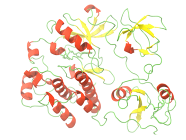 SRC (35.0%) | 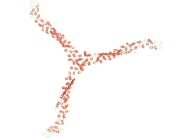 CLTC (15.8%) | 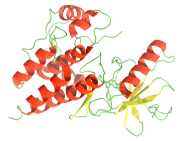 RAF1 (14.3%) | 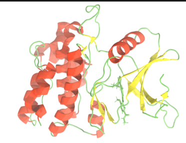 MAP3K5 (33.3%) | 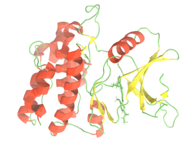 MAP3K5 (20.0%) |
| 5 | 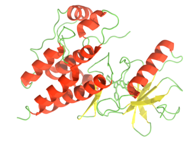 RAF1 (28.6%) | 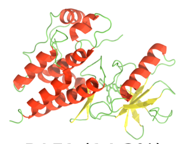 RAF1 (14.3%) | | 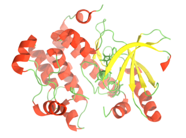 MAP2K1 (33.3%) | 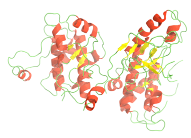 MAP2K4 (20.0%) |
| 6 | 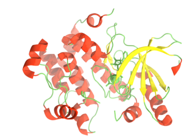 MAP2K1 (22.2%) | 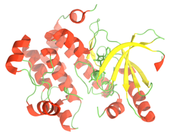 MAP2K1 (11.1%) | | 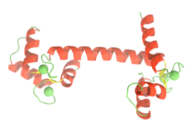 CALM1 (20.0%) | 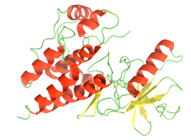 RAF1 (14.3%) |
| 7 | 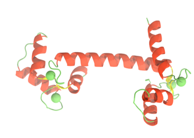 CALM1 (20.0%) | 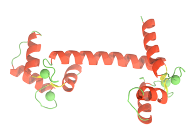 CALM1 (10.0%) | | 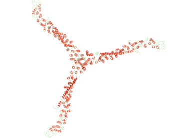 CLTC (15.8%) | Nuclear localization signal<br>NLS (14.3%) |
| 8 | 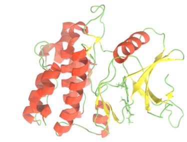 MAP3K5 (16.7%) | | | 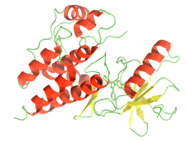 RAF1 (14.3%) | 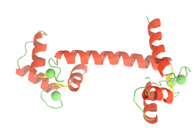 CALM1 (10.0%) |

Figure S14: Cartoon representations of downstream effectors or functional regions, rank-ordered based on their likeliness of preferring each $\beta$arr1 system. The percentage of residues influenced in the corresponding regions is indicated in parentheses.

| Sl No. | $\beta$arr2V2R(FP) | $\beta$arr2_V2R(-P1) | $\beta$arr2_V2R(-P3) | $\beta$arr2_V2R(-P5) | $\beta$arr2_V2R(-P7/8) |
|---|---|---|---|---|---|
| 1 | 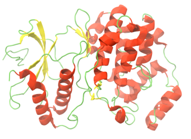 MAPK1 (33.3%) | 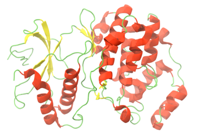 MAPK1 (100.0%) | 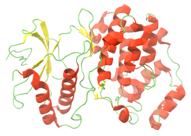 MAPK1 (66.7%) | 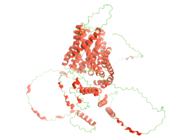 SLC9A5 (12.9%) | 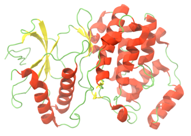 MAPK1 (66.7%) |
| 2 | 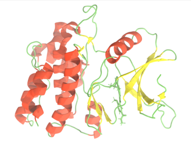 MAP3K5 (26.3%) | 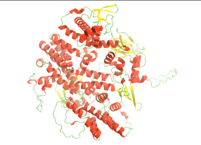 RAPGEF3 (38.1%) | 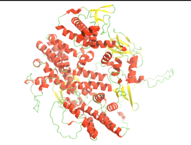 RAPGEF3 (42.9%) | 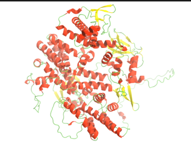 RAPGEF3 (9.5%) | 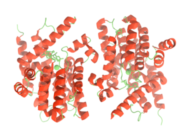 PDE4D (50.0%) |
| 3 | 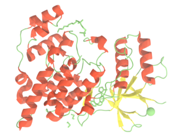 MAPK10 (13.8%) | 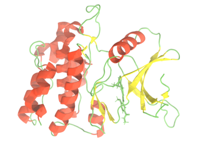 MAP3K5 (36.8%) | 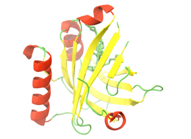 PTGDS (33.3%) | 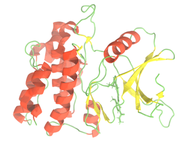 MAP3K5 (5.3%) | 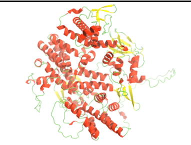 RAPGEF3 (38.1%) |
| 4 | 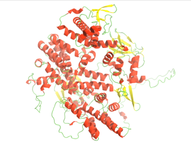 RAPGEF3 (9.5%) | 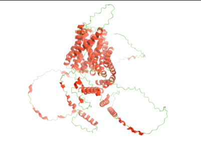 SLC9A5 (35.5%) | 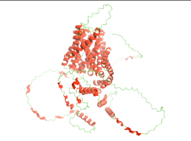 SLC9A5 (17.2%) | 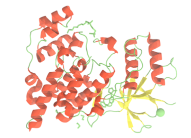 MAPK10 (3.5%) | 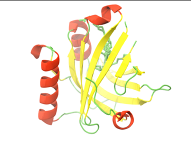 PTGDS (33.3%) |
| 5 | 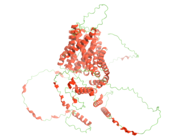 SLC9A5 (8.6%) | 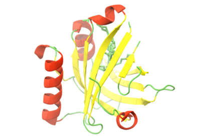 PTGDS (33.3%) | 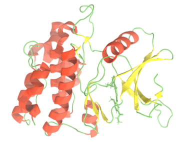 MAP3K5 (15.8%) | 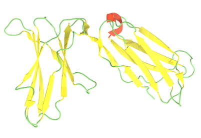 KIR2DL1 (1.8%) | 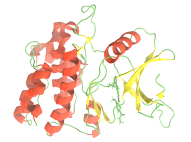 MAP3K5 (31.6%) |
| 6 | 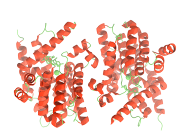 PDE4D (8.3%) | 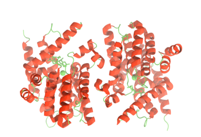 PDE4D (33.3%) | 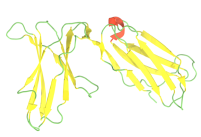 KIR2DL1 (8.9%) | | 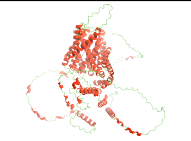 SLC9A5 (28.0%) |
| 7 | 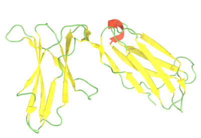 KIR2DL1 (5.4%) | 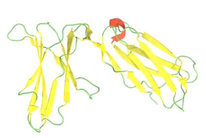 KIR2DL1 (16.1%) | 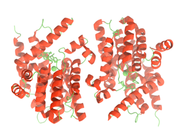 PDE4D (8.3%) | | 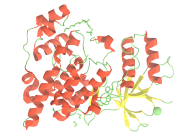 MAPK10 (27.6%) |
| 8 | | 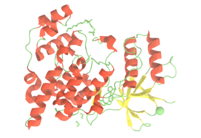 MAPK10 (10.3%) | 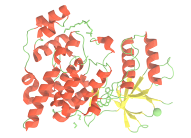 MAPK10 (3.5%) | | 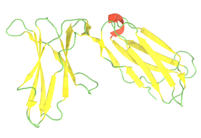 KIR2DL1 (25.0%) |

Figure S15: Cartoon representations of downstream effectors or functional regions, rank-ordered based on their likeliness of preferring each $\beta$arr2 system. The percentage of residues influenced in the corresponding regions is indicated in parentheses.

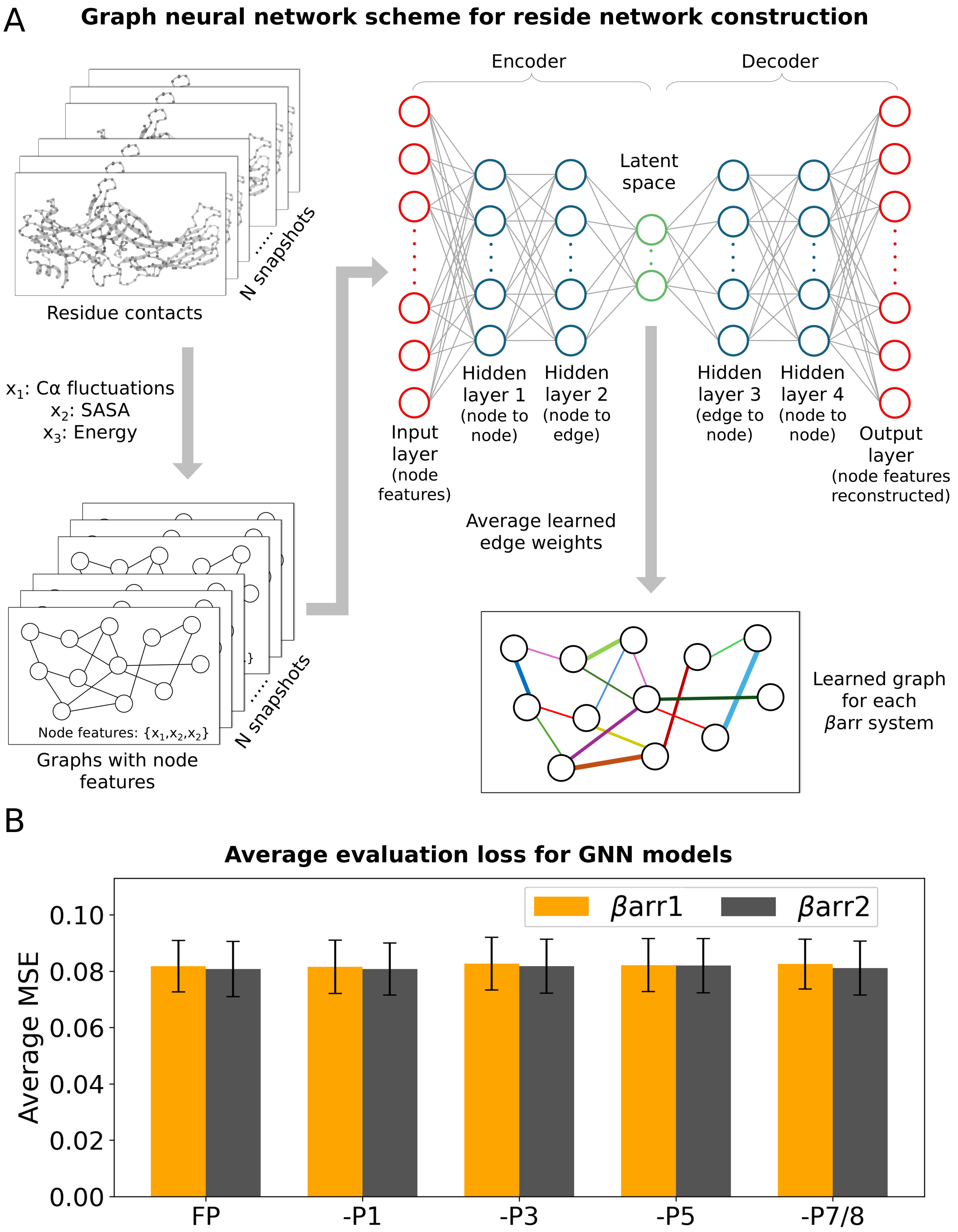


Figure S16: **(A)** Schematic representation of the GNN-based autoencoder. Input graphs, constructed from structural snapshots, use nodes representing residues and include features such as $C_\alpha$ atom fluctuations, SASA, and the sum of interaction energy weights. The architecture consists of an encoder and decoder composed of fully connected layers, connected through a latent space. This latent space captures the learned edge representations for each input graph. **(B)** Average mean squared error (MSE) values obtained for each phospho-system in both $\beta$arr1 and $\beta$arr2.

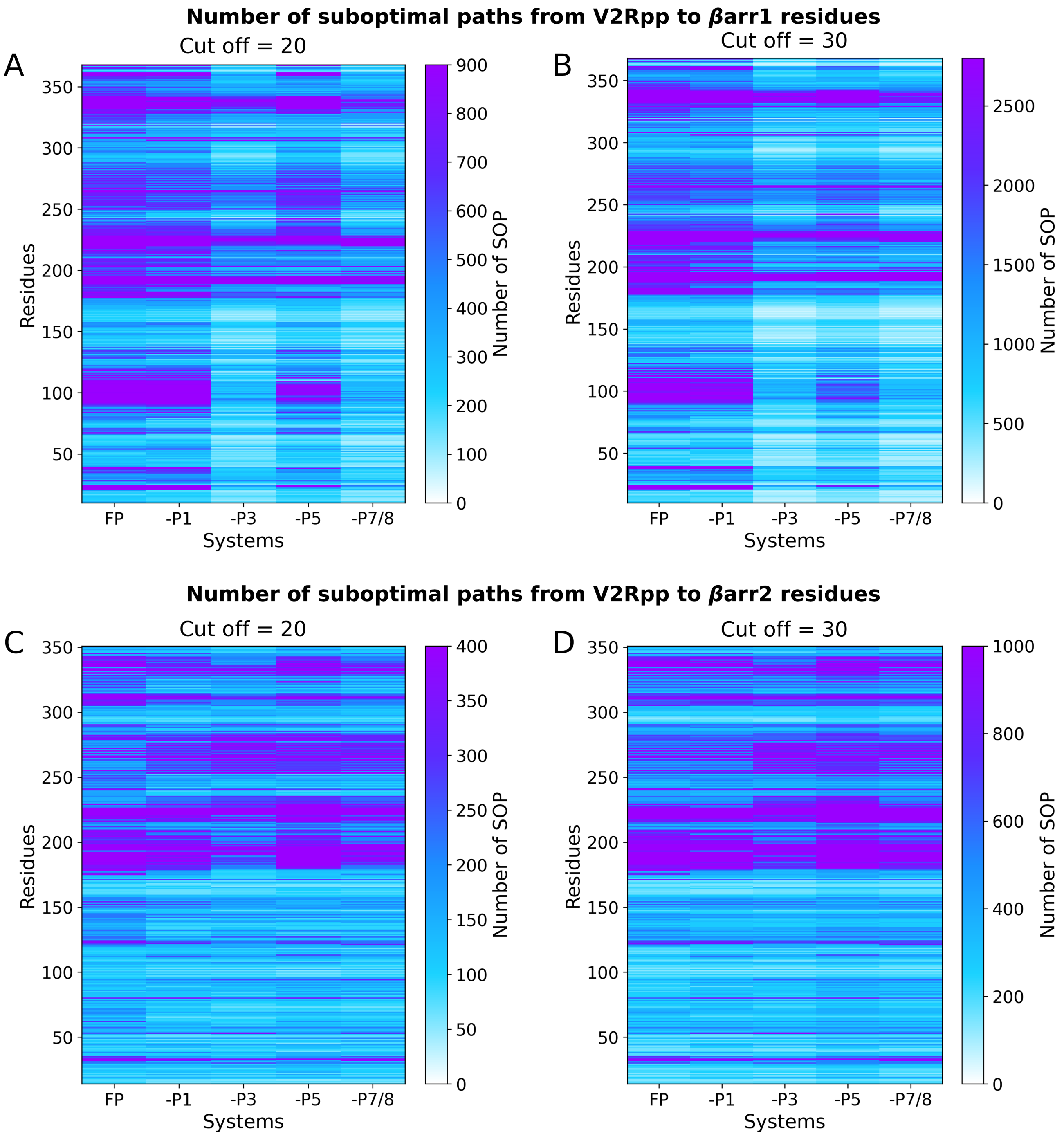


Figure S17: Number of suboptimal paths from V2Rpp residues to $\beta$arr1 residues across all systems for **(A)** $l_{\mathtt{cut}}$ = 20 and **(B)** $l_{\mathtt{cut}}$ = 30. Corresponding results for $\beta$arr2 systems are shown in **(C)** $l_{\mathtt{cut}}$ = 20 and **(D)** $l_{\mathtt{cut}}$ = 30.

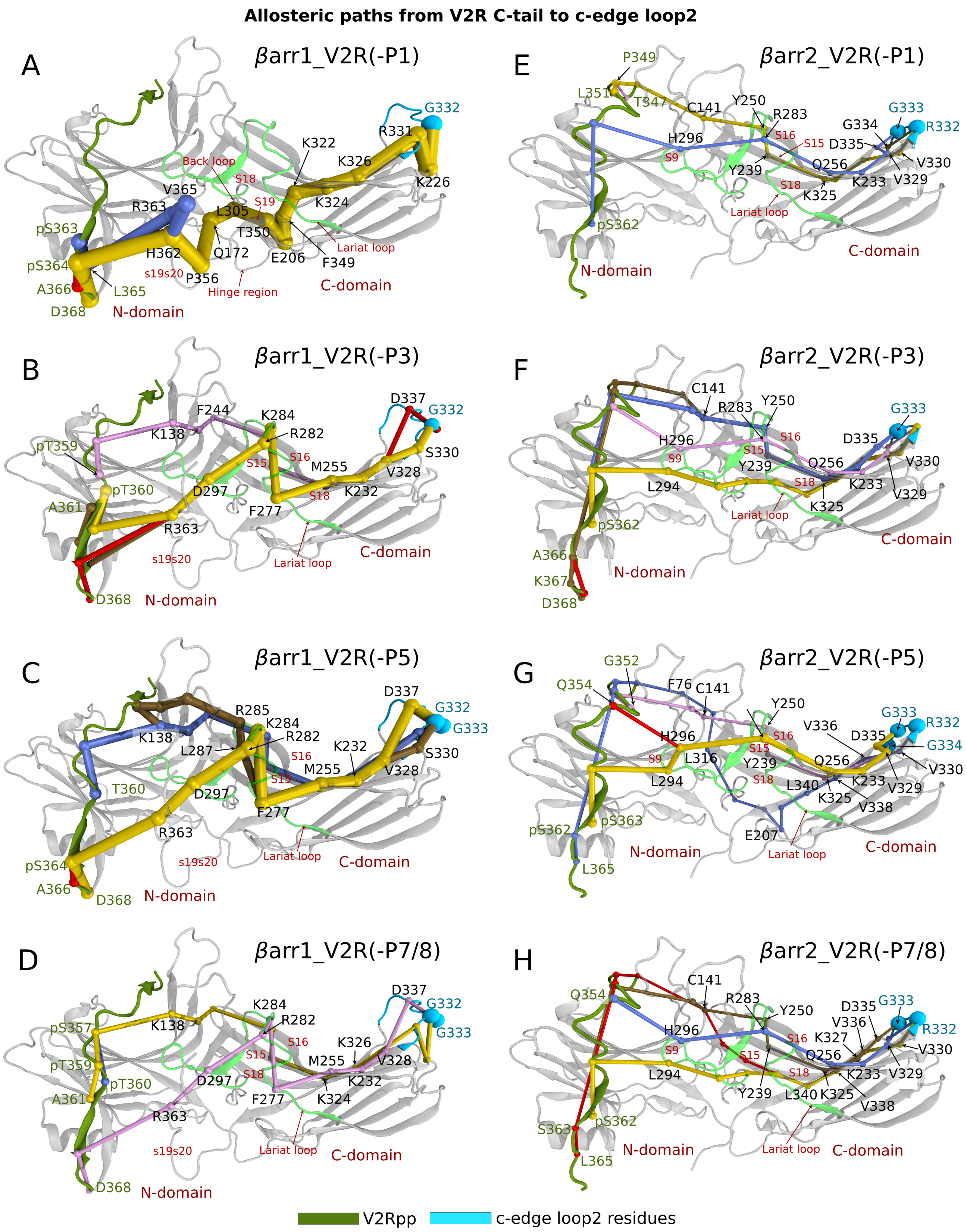


Figure S18: Allosteric communication pathways from V2Rpp to the c-edge loop2. The top five suboptimal paths ranked by thickness are shown in yellow, blue, brown, red, and pink (note that some of them overlap at non-terminal residues) for different phospho-variants of **(A-D)** $\beta$arr1 and **(E-H)** $\beta$arr2. V2Rpp is shown in dark green, and c-edge loop2 in cyan. The representations are based on the fully phosphorylated structures (PDB IDs: 4JQI and 8i10), with missing residues modeled. Key residues for each system, identified by a relative frequency $\geq 0.2$, are highlighted.

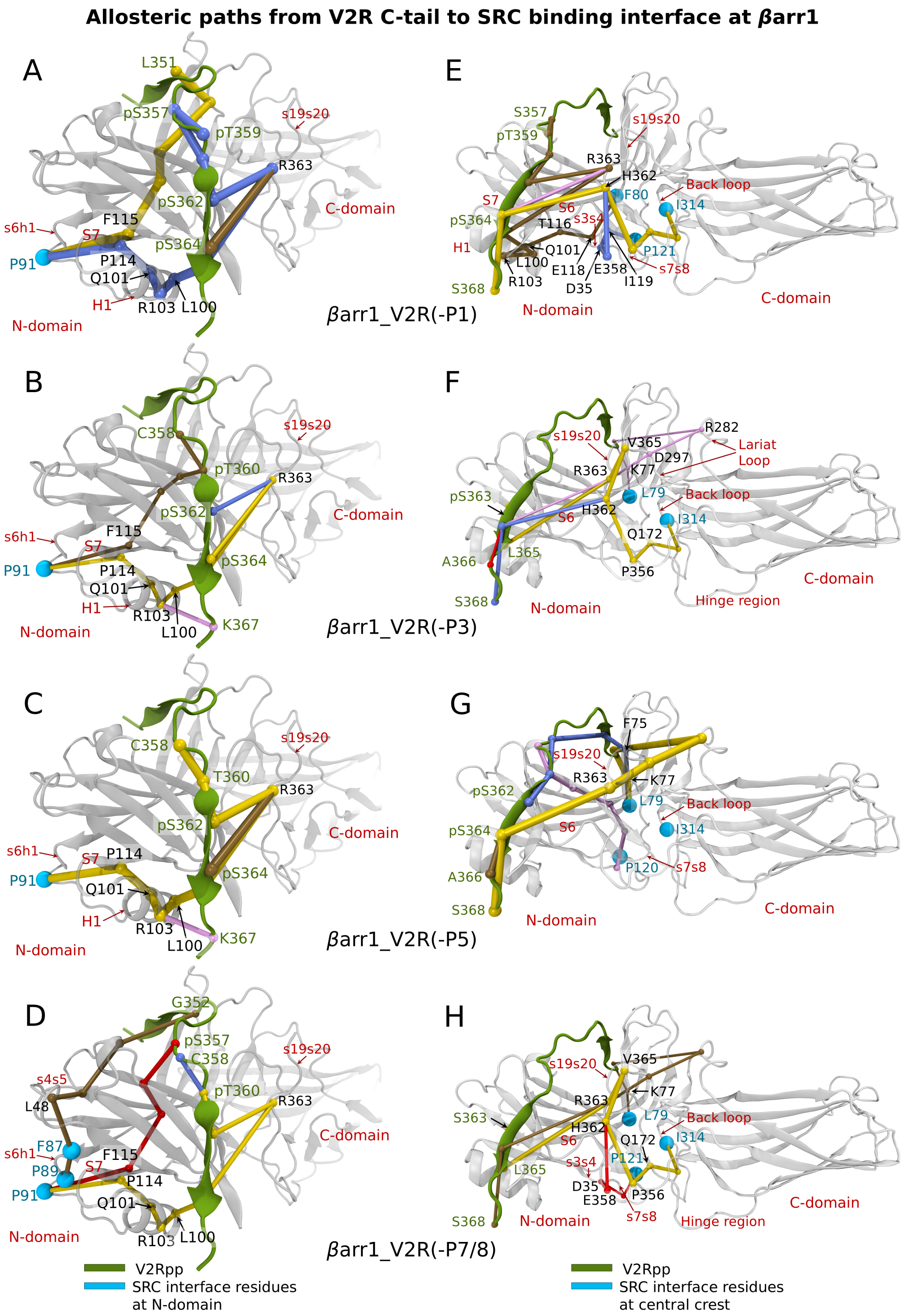


Figure S19: Allosteric communication pathways from V2Rpp to the SRC interfaces at $\beta$arr1: **(A-D)** polyproline motifs in s6h1 and **(E-H)** central crest for the phospho-variants. The top five suboptimal paths ranked by thickness are shown in yellow, blue, brown, red, and pink (note that some of them overlap at non-terminal residues). V2Rpp is shown in dark green, and SRC interface residues in cyan. The representations are based on fully phosphorylated structure (PDB ID 4JQI), with missing residues modeled. Key residues for each system, identified by a relative frequency $\geq 0.2$, are highlighted.

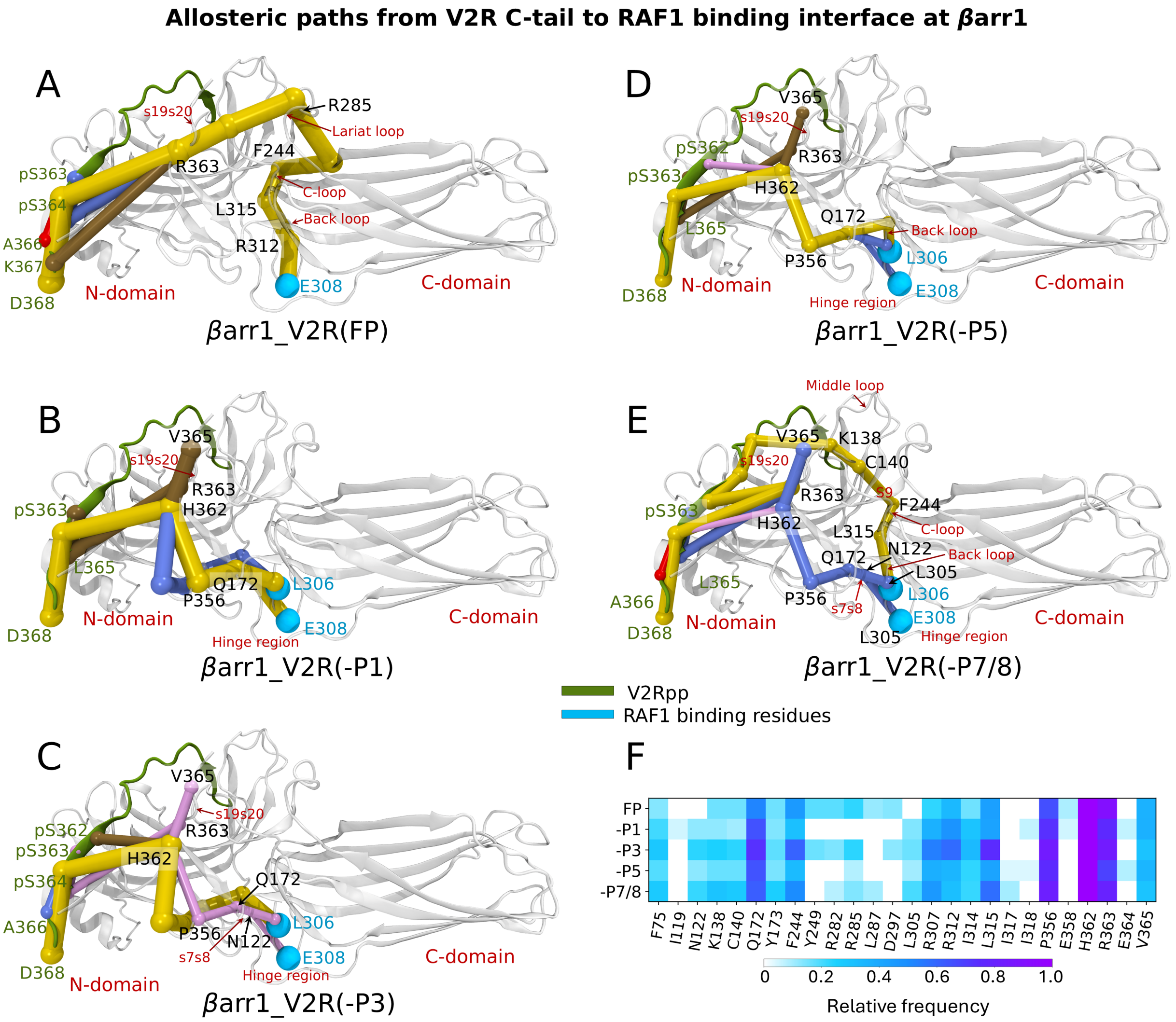


Figure S20: **(A-E)** Allosteric communication pathways from V2Rpp to the RAF1 interface at $\beta$arr1. The top five suboptimal paths ranked by thickness are shown in yellow, blue, brown, red, and pink (note that some of them overlap at non-terminal residues). V2Rpp is shown in dark green, and RAF1 interface residues in cyan. The representations are based on the fully phosphorylated structure (PDB ID 4JQI), with missing residues modeled. **(F)** The top 20 residues, ranked by their occurrence in suboptimal paths, are shown as heatmaps for each system. Key residues, identified by a relative frequency $\geq 0.2$, are highlighted in Figures A-F.

#### Allosteric paths from V2R C-tail to MAP2K1 binding interface at βarr1

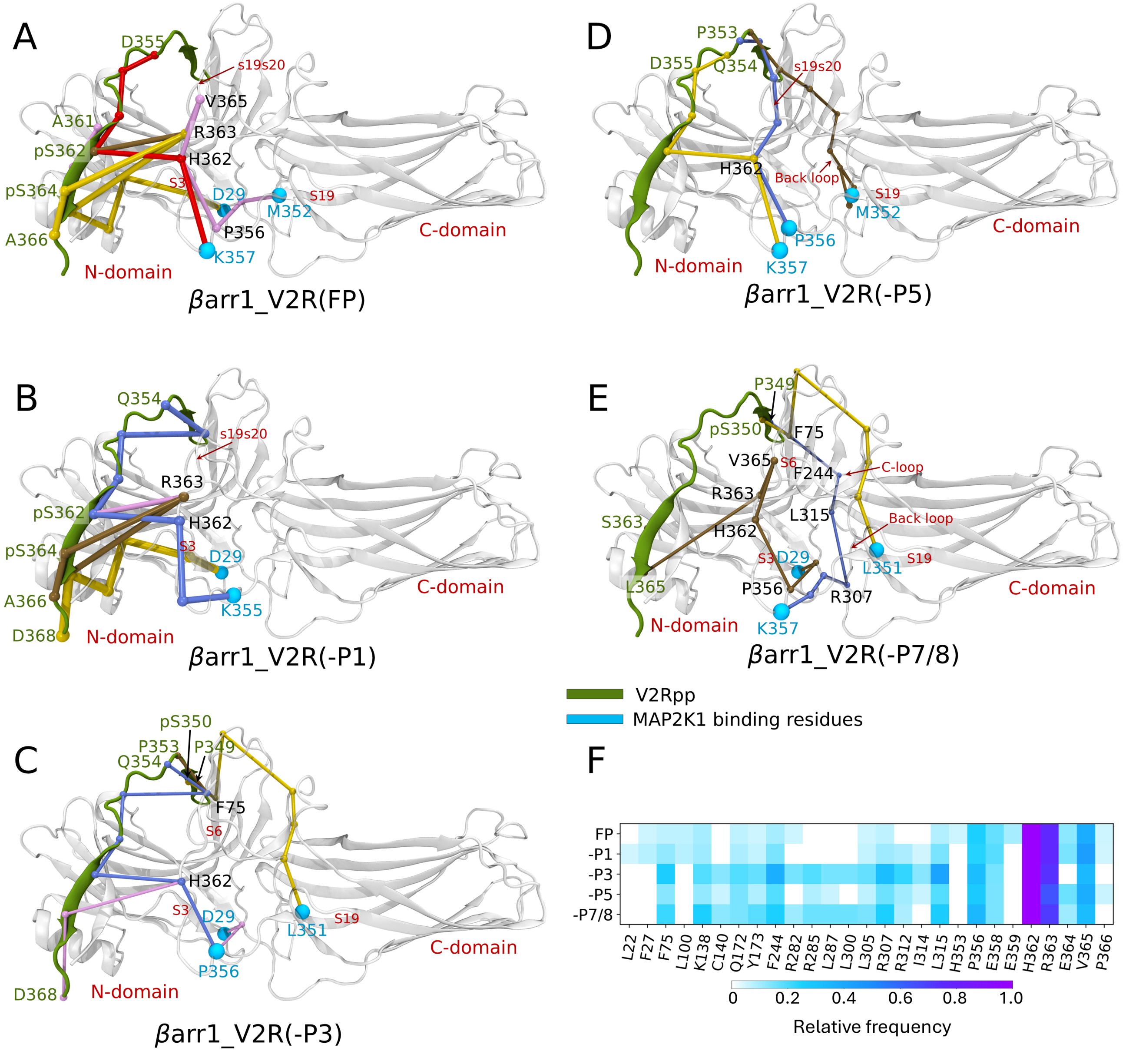


Figure S21: **(A-E)** Allosteric communication pathways from V2Rpp to the MAP2K1 interface at $\beta$arr1. The top five suboptimal paths ranked by thickness are shown in yellow, blue, brown, red, and pink (note that some of them overlap at non-terminal residues). V2Rpp is shown in dark green, and MAP2K1 interface residues in cyan. The representations are based on the fully phosphorylated structure (PDB ID 4JQI), with missing residues modeled. **(F)** The top 20 residues, ranked by their occurrence in suboptimal paths, are shown as heatmaps for each system. Key residues, identified by a relative frequency $\geq 0.2$, are highlighted in Figures A-F.

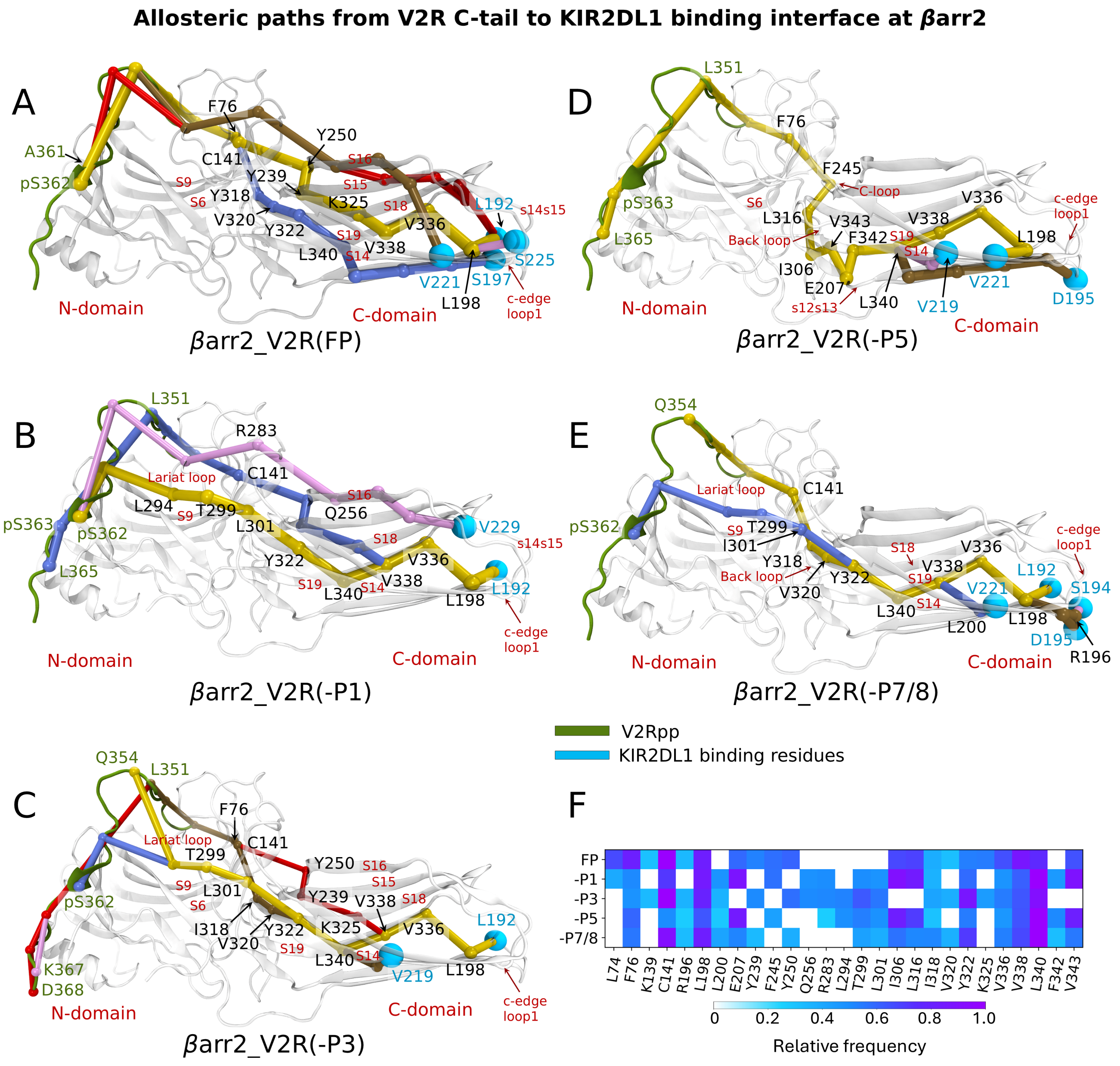


Figure S22: **(A-E)** Allosteric communication pathways from V2Rpp to the KIR2DL1 interface at $\beta$arr2. The top five suboptimal paths ranked by thickness are shown in yellow, blue, brown, red, and pink (note that some of them overlap at non-terminal residues). V2Rpp is shown in dark green, and KIR2DL1 interface residues in cyan. The representations are based on the fully phosphorylated structure (PDB ID 8I10), with missing residues modeled. **(F)** The top 20 residues, ranked by their occurrence in suboptimal paths, are shown as heatmaps for each system. Key residues, identified by a relative frequency $\geq 0.2$, are highlighted in Figures A-F.

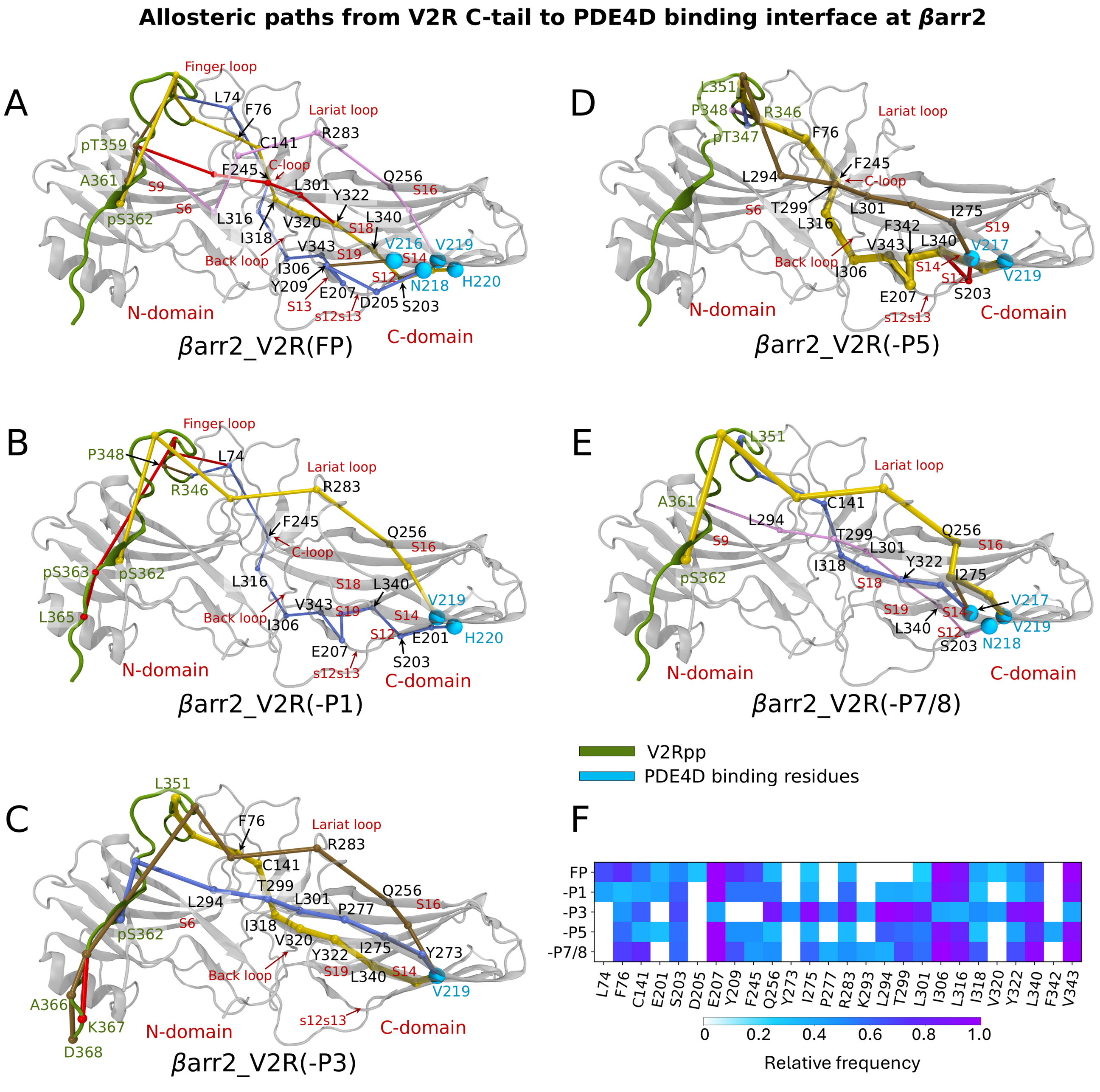


Figure S23: **(A-E)** Allosteric communication pathways from V2Rpp to the PDE4D interface at $\beta$arr2. The top five suboptimal paths ranked by thickness are shown in yellow, blue, brown, red, and pink (note that some of them overlap at non-terminal residues). V2Rpp is shown in dark green, and PDE4D interface residues in cyan. The representations are based on the fully phosphorylated structure (PDB ID 8I10), with missing residues modeled. **(F)** The top 20 residues, ranked by their occurrence in suboptimal paths, are shown as heatmaps for each system. Key residues, identified by a relative frequency $\geq 0.2$, are highlighted in Figures A-F.

**Allosteric paths from V2R C-tail to PTGDS binding interface at $\beta$arr2**

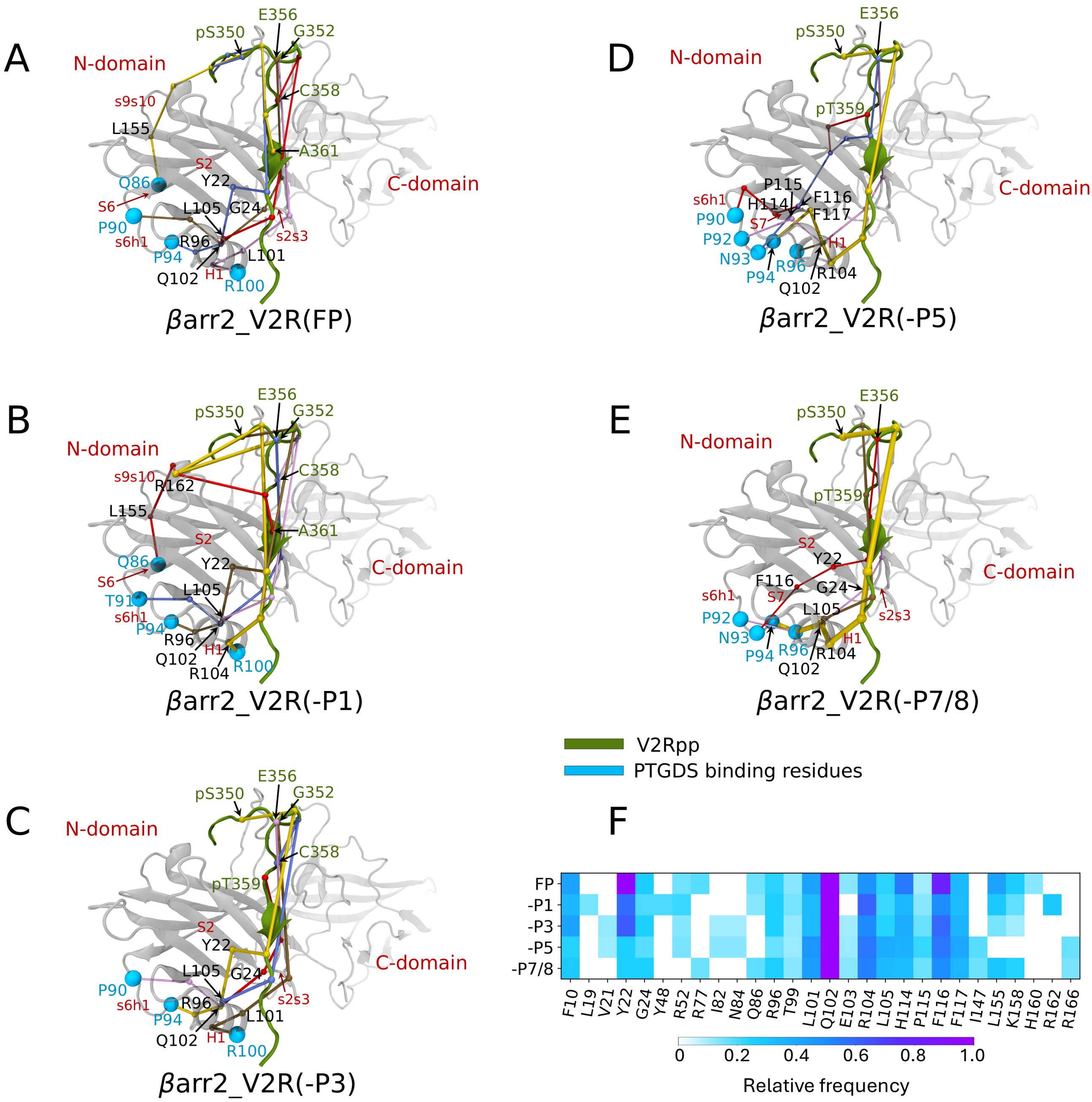


Figure S24: **(A-E)** Allosteric communication pathways from V2Rpp to the PTGDS interface at $\beta$arr2. The top five suboptimal paths ranked by thickness are shown in yellow, blue, brown, red, and pink (note that some of them overlap at non-terminal residues). V2Rpp is shown in dark green, and PTGDS interface residues in cyan. The representations are based on the fully phosphorylated structure (PDB ID 8I10), with missing residues modeled. **(F)** The top 20 residues, ranked by their occurrence in suboptimal paths, are shown as heatmaps for each system. Key residues, identified by a relative frequency $\geq 0.2$, are highlighted in Figures A-F.

Table S1: Occupancy of specific interactions present in the fully phosphorylated system compared to those in the –P1 system in site 1, highlighting the impact of the absence of phosphorylation at T347 in the latter for $\beta$arr1.

| Interaction | $\beta$arr1 residue | –P1 residue | Occupancy (%) | FP residue | Occupancy (%) |
|---|---|---|---|---|---|
| H-bond | Y63 | T347 | - | pT347 | 82.14 |
| | R65 | | - | | 85.19 |
| | K77 | | - | | 79.56 |
| Anion-$\pi$ interactions | F75 | | - | | 16.93 |
| Polar interactions | R76 | | - | | - |
| Hydrophobic interaction | F75 | | - | | 44.75 |
| | R76 | | - | | 71.41 |
| Charge-charge interactions | R65 | | - | | 84.45 |
| | K77 | | - | | 63.18 |
| vdW interactions | Y63 | | - | | 82.35 |
| | R65 | | - | | 89.46 |
| | F75 | | - | | 73.37 |
| | R76 | | 54.56 | | 93.32 |
| | K77 | | - | | 82.27 |
| | K147 | | 51.49 | | - |

Table S2: Occupancy of specific interactions present in the fully phosphorylated system compared to those in the –P3 system in site 3, highlighting the impact of the absence of phosphorylation at S357 in the latter for $\beta$arr1.

| Interaction | $\beta$arr1 residue | –P3 residue | Occupancy (%) | FP residue | Occupancy (%) |
|---|---|---|---|---|---|
| H-bond | K11 | S357 | 20.15 | pS357 | 64.96 |
| | K160 | | - | | 64.96 |
| | R165 | | - | | 85.69 |
| Polar interactions | K11 | | - | | - |
| | K160 | | - | | - |
| Charge-charge interactions | K11 | | - | | 54.90 |
| | K160 | | - | | 56.32 |
| | K138 | | - | | - |
| | R165 | | - | | 79.23 |
| vdW interactions | K11 | | 40.75 | | 65.49 |
| | K160 | | - | | 66.42 |
| | R165 | | 16.20 | | 85.69 |

Table S3: Occupancy of specific interactions present in the fully phosphorylated system compared to those in the –P5 system in site 5, highlighting the impact of the absence of phosphorylation at T360 in the latter for $\beta$arr1.

| Interaction | $\beta$arr1 residue | –P5 residue | Occupancy (%) | FP residue | Occupancy (%) |
|---|---|---|---|---|---|
| H-bond | K11 | pS357 | | pS357 | 64.06 |
| | K160 | | 14.15 | | 64.96 |
| | R165 | | 14.29 | | 85.69 |
| Polar interactions | K160 | | - | | - |
| Charge-charge interactions | K138 | | - | | - |
| | K11 | | - | | 54.90 |
| | K160 | | 13.31 | | 56.32 |
| | R165 | | 14.32 | | 79.23 |
| vdW interactions | K11 | | - | | 65.49 |
| | K160 | | 14.17 | | 66.42 |
| | R165 | | 14.29 | | 85.69 |
| | K292 | | 45.16 | | |
| | K294 | | 42.36 | | 16.78 |
| H-bond | K11 | pT359 | 75.83 | pT359 | - |
| | K294 | | 21.34 | | 77.76 |
| Charge-charge interactions | K11 | | 66.78 | | - |
| | R25 | | - | | - |
| | K294 | | 13.84 | | 62.01 |
| H-bond | K11 | T360 | - | pT360 | 97.63 |
| | R25 | | 25.00 | | 100.00 |
| | K294 | | - | | 78.52 |
| Polar interactions | R25 | | - | | - |
| Hydrophobic interaction | L166 | | 32.76 | | 59.15 |
| Charge-charge interactions | K11 | | | | 71.53 |
| | R25 | | | | 79.49 |
| | K294 | | | | 62.83 |
| vdW interactions | K10 | | 67.65 | | 71.95 |
| | K11 | | 32.79 | | 98.46 |
| | D25 | | 64.71 | | 100.00 |
| | K294 | | - | | 63.46 |
| | L166 | | 53.94 | | 79.23 |

Table S4: Occupancy of specific interactions present in the fully phosphorylated system compared to those in the –P7/8 system in site 7/8, highlighting the impact of the absence of phosphorylation at S363 and S364 in the latter for $\beta$arr1.

| Interaction | $\beta$arr1 residue | –P7/8 residue | Occupancy (%) | FP residue | Occupancy (%) |
|---|---|---|---|---|---|
| H-bond | R7 | S363 | - | pS363 | - |
| | V8 | | 99.77 | | 99.85 |
| | K10 | | - | | 99.47 |
| | Y21 | | - | | 37.62 |
| | K107 | | - | | 96.46 |
| Polar interactions | R7 | | - | | - |
| | V8 | | 94.04 | | - |
| Charge-charge interactions | K10 | | - | | 81.14 |
| | K107 | | - | | 69.41 |
| Hydrophobic interaction | R7 | | 31.96 | | 50.37 |
| vdW interactions | R7 | | 88.13 | | 95.44 |
| | V8 | | 99.94 | | 99.98 |
| | K10 | | 33.26 | | 99.62 |
| | K107 | | - | | 96.55 |
| H-bond | T6 | S364 | - | pS364 | - |
| | K107 | | - | | 87.13 |
| | R7 | | 14.11 | | 81.15 |
| Polar interactions | R7 | | - | | - |
| Charge-charge interactions | R7 | | - | | 69.00 |
| | K107 | | - | | 11.80 |
| vdW interactions | R7 | | 50.36 | | 94.31 |
| | K107 | | 22.27 | | 93.95 |
| | T6 | | 52.36 | | 11.31 |

Table S5: Occupancy of specific interactions present in the fully phosphorylated system compared to those in the –P1 system in site 1, highlighting the impact of the absence of phosphorylation at T347 in the latter for $\beta$arr2.

| Interaction | $\beta$arr2 residue | –P1 residue | Occupancy (%) | FP residue | Occupancy (%) |
|---|---|---|---|---|---|
| H-bond | R63 | T347 | - | pT347 | 77.95 |
| | R77 | | - | | 80.10 |
| | R148 | | - | | 99.71 |
| Anion-$\pi$ interactions | F76 | | - | | - |
| Polar interactions | R77 | | - | | - |
| Charge-charge interactions | R63 | | - | | 76.21 |
| | R77 | | - | | 73.88 |
| | R148 | | - | | 95.19 |
| Hydrophobic interaction | F76 | | - | | - |
| vdW interactions | R63 | | - | | 78.46 |
| | R77 | | - | | 80.14 |
| | R148 | | - | | 99.76 |

Table S6: Occupancy of specific interactions present in the fully phosphorylated system compared to those in the –P3 system in site 3, highlighting the impact of the absence of phosphorylation at S357 in the latter for $\beta$arr2.

| Interaction | $\beta$arr2 residue | –P3 residue | Occupancy (%) | FP residue | Occupancy (%) |
|---|---|---|---|---|---|
| H-bond | K12 | S357 | - | pS357 | - |
| Polar interactions | K12 | | - | | - |
| | K161 | | - | | - |
| Charge-charge interactions | K139 | | - | | - |
| | R166 | | - | | - |

Table S7: Occupancy of specific interactions present in the fully phosphorylated system compared to those in the –P5 system in site 5, highlighting the impact of the absence of phosphorylation at T360 in the latter for $\beta$arr2.

| Interaction | $\beta$arr2 residue | –P5 residue | Occupancy (%) | FP residue | Occupancy (%) |
|---|---|---|---|---|---|
| H-bond | K12 | pS357 | 51.89 | pS357 | - |
| Polar interactions | K161 | | - | | - |
| Charge-charge interactions | K139 | | - | | - |
| | R166 | | - | | - |
| vdW interactions | K12 | | 53.05 | | - |
| H-bond | K12 | pT359 | - | pT359 | 96.24 |
| | R26 | | 77.61 | | - |
| | R170 | | 46.68 | | - |
| | R162 | | - | | 51.40 |
| | K295 | | 35.19 | | - |
| Charge-charge interactions | R26 | | 77.53 | | - |
| | K12 | | - | | 69.64 |
| | R162 | | - | | 51.91 |
| vdW interactions | K12 | | - | | 99.60 |
| | R162 | | - | | 51.79 |
| H-bond | K12 | T360 | - | pT360 | 89.98 |
| | R26 | | 28.96 | | 100.00 |
| | K295 | | - | | 36.16 |
| | R66 | | - | | 42.86 |
| Polar interactions | R26 | | - | | - |
| Charge-charge interactions | K12 | | - | | 61.08 |
| | R26 | | - | | 65.46 |
| vdW interactions | K11 | | 57.11 | | 35.25 |
| | K12 | | 10.52 | | 97.47 |
| | R26 | | 60.72 | | 100.00 |

Table S8: Occupancy of specific interactions present in the fully phosphorylated system compared to those in the –P7/8 system in site 7/8, highlighting the impact of the absence of phosphorylation at S363 and S364 in the latter for $\beta$arr2.

| Interaction | $\beta$arr2 residue | –P7/8 residue | Occupancy (%) | FP residue | Occupancy (%) |
|---|---|---|---|---|---|
| H-bond | R8 | S363 | - | pS363 | - |
| | V9 | | 96.11 | | 99.96 |
| | K11 | | - | | 99.78 |
| | Y22 | | - | | 52.71 |
| | K108 | | - | | 98.54 |
| Polar interactions | R8 | | - | | - |
| | V9 | | - | | - |
| Charge-charge interactions | K11 | | - | | 83.23 |
| | K108 | | - | | 68.14 |
| Hydrophobic interaction | R8 | | - | | 55.66 |
| vdW interactions | R8 | | 91.21 | | 92.04 |
| | V9 | | 79.06 | | 100.00 |
| | K11 | | 30.90 | | 99.83 |
| | Y22 | | - | | 53.06 |
| | K108 | | - | | 98.87 |
| H-bond | T7 | S364 | - | pS364 | - |
| | K108 | | - | | 93.33 |
| | R8 | | - | | 51.99 |
| Polar interactions | R8 | | - | | - |
| Charge-charge interactions | R8 | | - | | 42.26 |
| | K108 | | - | | 35.70 |
| vdW interactions | R8 | | - | | 90.05 |
| | K108 | | - | | 98.77 |

Table S9: Residues in $\beta$arr1 that interact with various effectors.

| Effector | $\beta$arr1 residues and Structural Element | | Effector | $\beta$arr1 residues and Structural Element | |
|---|---|---|---|---|---|
| MAP2K4 | 66, 67, 69, 77, 78 | s5s6 (Finger loop), S6 | RAF1 | 302-308 | s17s18 (back loop) |
| MAP3K5 (ASK1) | 6-35 | S1, s1s2, S2, s2s3, S3, s3s4 | SRC | 75, 77-80, 314, 87-89, 91 | S6, s17s18 (back loop), s6h1 |
| MAP2K1 | 26, 29, 351-357 | S3, S19, s19s20 | CLTC | 188-193, 326-332, 334-339 | S11, s11s12 (c-edge loop1), S18, s18s19 (c-edge loop2) |
| CALM1 | 70, 71, 73, 167, 191, 234, 238, 246, 257, 308 | S6, S10, S15, s15s16 (C-loop), S16, s17s18 (back loop) | NLS | 157-161, 169–170 | s9s10, S10 |
| GNAS | 33 | s3s4 | | | |

Table S10: Residues in $\beta$arr2 that interact with various effectors.

| Effector | $\beta$arr2 residues | Struct. element | Effector | $\beta$arr2 residues | Struct. element |
|---|---|---|---|---|---|
| MAP3K5 (ASK1) | 9-23, 25, 27, 29, 30 | S2, s2s3, S3 | PDE4D | 18, 20, 25, 26, 215-220, 286, 291 | s1s2, S2, s2s3, S14, s17s18 (lariat loop) |
| MAPK1 | 285, 286, 295 | s17s18 (lariat loop) | PTGDS | 86-100 | S6, s6h1, H1 |
| MAPK10 (JNK3) | 9-23, 196-198, 208, 215, 216, 264, 278, 280, 343, 350-353 | S1-S3, s11s12 (c-edge loop1), S12, s13s14, s16s17, S19, s17s18 (lariat loop), s19s20 | KIR2DL1 | 185-240 | S11, s11s12 (c-edge loop1), S12, s12s13, S13, s13s14, S14, s14s15, S15 |
| SLC9A5 (NHE5) | 52-78, 133-183, 318-332 | S5, s5s6, S6, S9, s9s10, S10, s10s11, S18, s18s19 | RAPGEF3 | 286-306 | s17s18 (lariat loop) |

### References


[1] Latorraca, N. R.; Masureel, M.; Hollingsworth, S. A.; Heydenreich, F. M.; Suomivuori, C.-M.; Brinton, C.; Townshend, R. J.; Bouvier, M.; Kobilka, B. K.; Dror, R. O. *Cell* **2020**, *183*, 1813–1825.

[2] Roe, D. R.; Cheatham III, T. E. *Journal of chemical theory and computation* **2013**, *9*, 3084–3095.

[3] McGibbon, R. T.; Beauchamp, K. A.; Harrigan, M. P.; Klein, C.; Swails, J. M.; Hernández, C. X.; Schwantes, C. R.; Wang, L.-P.; Lane, T. J.; Pande, V. S. *Biophysical journal* **2015**, *109*, 1528–1532.

[4] Latorraca, N. R.; Wang, J. K.; Bauer, B.; Townshend, R. J.; Hollingsworth, S. A.; Olivieri, J. E.; Xu, H. E.; Sommer, M. E.; Dror, R. O. *Nature* **2018**, *557*, 452–456.

[5] Madhu, M. K.; Shewani, K.; Murarka, R. K. *Journal of Chemical Information and Modeling* **2024**, *64*, 449–469.

[6] Venkatakrishnan, A. J.; Fonseca, R.; Ma, A. K.; Hollingsworth, S. A.; Chemparathy, A.; Hilger, D.; Kooistra, A. J.; Ahmari, R.; Babu, M. M.; Kobilka, B. K.; Dror, R. O. *BioRxiv* **2019**,

[7] Darbellay, G. A.; Vajda, I. *IEEE Transactions on Information Theory* **1999**, *45*, 1315–1321.

[8] Chen, K. Neoneuron/minfo: Mutual information estimator with adaptive partitioning algorithm (C++/OpenMP accelerated). `https://github.com/NeoNeuron/minfo`, 2025; Accessed: 2025-01-05.

[9] Press, W. H.; Flannery, B. P.; Teukolsky, S. A.; Vetterling, W. T. Numerical Recipes: The art of scientific computing (Cambridge. 1986.

[10] Wold, S.; Esbensen, K.; Geladi, P. *Chemometrics and intelligent laboratory systems* **1987**, *2*, 37–52.

[11] Schölkopf, B.; Smola, A.; Müller, K.-R. Kernel principal component analysis. International conference on artificial neural networks. 1997; pp 583–588.

[12] Antony, J.; Bardhan Anand, R.; Kumar, M.; Tiwari, M. K. *Journal of Manufacturing Technology Management* **2006**, *17*, 908–925.

[13] Schölkopf, B. Learning with kernels: support vector machines, regularization, optimization, and beyond. 2002.

[14] Van der Maaten, L.; Hinton, G. *Journal of machine learning research* **2008**, *9*.

[15] Appadurai, R.; Koneru, J. K.; Bonomi, M.; Robustelli, P.; Srivastava, A. *Journal of chemical theory and computation* **2023**, *19*, 4711–4727.

[16] Belkina, A. C.; Ciccolella, C. O.; Anno, R.; Halpert, R.; Spidlen, J.; Snyder-Cappione, J. E. *Nature communications* **2019**, *10*, 5415.

[17] McInnes, L.; Healy, J.; Melville, J. *arXiv preprint arXiv:1802.03426* **2018**,

[18] He, Q.-T.; Xiao, P.; Huang, S.-M.; Jia, Y.-L.; Zhu, Z.-L.; Lin, J.-Y.; Yang, F.; Tao, X.-N.; Zhao, R.-J.; Gao, F.-Y.; others *Nature Communications* **2021**, *12*, 2396.

[19] Crépieux, P.; Poupon, A.; Langonné-Gallay, N.; Reiter, E.; Delgado, J.; Schaefer, M. H.; Bourquard, T.; Serrano, L.; Kiel, C. *Frontiers in Endocrinology* **2017**, *8*, 32.

[20] Paszke, A.; Gross, S.; Massa, F.; Lerer, A.; Bradbury, J.; Chanan, G.; Killeen, T.; Lin, Z.; Gimelshein, N.; Antiga, L.; others *Advances in neural information processing systems* **2019**, *32*.

[21] Fey, M.; Lenssen, J. E. *arXiv preprint arXiv:1903.02428* **2019**,

[22] Sethi, A.; Eargle, J.; Black, A. A.; Luthey-Schulten, Z. *Proceedings of the National Academy of Sciences* **2009**, *106*, 6620–6625.

[23] Rivalta, I.; Sultan, M. M.; Lee, N.-S.; Manley, G. A.; Loria, J. P.; Batista, V. S. *Proceedings of the National Academy of Sciences* **2012**, *109*, E1428–E1436.

[24] VanWart, A. T.; Eargle, J.; Luthey-Schulten, Z.; Amaro, R. E. *Journal of chemical theory and computation* **2012**, *8*, 2949–2961.

[25] Eargle, J.; Luthey-Schulten, Z. *RNA interaction networks. Bioinformatics* **2012**, *28*, 3000–3001.

[26] Humphrey, W.; Dalke, A.; Schulten, K. *Journal of molecular graphics* **1996**, *14*, 33–38.
